# PSMtags improve peptide sequencing and throughput in sensitive proteomics

**DOI:** 10.1101/2025.05.22.655509

**Authors:** Harrison Specht, Sarah Sipe, Maddy Yeh, David Barnes-Seeman, Mark Adamo, Kevin McDonnell, Michael P. Agius, Corinna Friedrich, Helena Rittenhouse, Joshua J. Abbott, Wai Kit Pang, Yanchang Huang, Kasa Shiva Raju, Wayne Vuong, Michael Alan Lee, Ahmet Yesilcimen, Joseph S. Brown, Chao Wang, Ryan T. Kelly, Bradley L. Pentelute, Nikolai Slavov

## Abstract

Mass spectrometry-based proteomics enables comprehensive characterization of proteins, but label-free approaches are challenging to scale to thousands of samples per day (SPD). Multiplexed data-independent approaches (plexDIA) can increase throughput but lack the mass tags to enable it. Here, we systematically screened a library of 576 compounds and identified several small molecules that, when conjugated to peptides, improve their detection and sequence identification. The lead molecule, PSMtag, substantially increases the detection of fragment b-ions, which increases the confidence of sequence identification and enhances *de novo* sequencing. PSMtags allow 9-plexDIA, using only stable isotopes of carbon, oxygen and nitrogen. As a result, it allows increasing sample throughput for plexDIA workflows. We demonstrate 1,130 SPD with 9-plexDIA from single-cell-level human proteomes, acquiring 27,520 protein datapoints in the same time label-free methods acquire 3,216. Our approach constitutes an expandable framework for developing mass tags to overcome existing limitations in multiplexed proteomics.

## Introduction

Mass spectrometry-based proteomics has a long history of using isotopes for increasing sample throughput via isotopic labeling. Isotopes can be introduced via metabolic labeling^1,2^ or as mass tags that covalently modify proteins or peptides, as reviewed by Pappireddi, Martin, and Wuhr^3^. Mass tags can be used with many types of samples and have been adopted for clinical studies, such as the National Cancer Institute’s Clinical Proteomic Tumor Analysis Consortium (CPTAC).

Early examples of mass tags include isotope-coded affinity tags^4^ and tandem mass tags^5^. The advantages of mass tags motivated successful commercialization, including isobaric and non-isobaric tags for relative and absolute quantitation (iTRAQ and mTRAQ, respectively)^6,7^ that allowed for combining up to four different proteome samples for simultaneous analysis. More recently, tandem mass tags (TMT, TMTpro)^5,8–10^ multiplex up to 35 samples. Some labeling strategies have been achieved with readily available reagents to achieve dimethylation^11,12^ or diethylation^13,14^ of tryptic or LysC-derived peptides. Other mass tags and amino acid reagents for multiplexing have highlighted do-it-yourself synthesis (DiLeu)^15,16^, low energy fragmentation of isobaric tags (EASI-tag)^17^ or increased isotopic multiplexing by using mass defects^18,19^. These chemical labeling strategies have allowed researchers to compare multiple biological conditions simultaneously, offering deeper insights into cellular processes, disease mechanisms, and biomarker discovery.

Despite these benefits, labeling with popular tags, such as mTRAQ and TMT/TMTpro, reduces peptide identification^20–22^. In some cases, labeling favors the detection of b-ions^20^, but it generally reduces the number of confidently identified peptides^20,21^. Since previous reports support that at least some types of labeling alter peptide fragmentation, we reasoned other chemical structures might hold the potential to further enhance peptide sequence identification through improved production of annotatable fragment ions. Towards this goal, we started exploring the large chemical space that has not yet been evaluated for mass tags.

Unlike isobaric mass tags, nonisobaric mass tags increase the number of distinct mass peaks, which raises the potential for interference to affect quantification. This effect is likely to be less pronounced in single-cell proteomics than in bulk samples, since fewer precursors are quantifiable from single cells, limiting the opportunities for peak interference. Previous work with 3-plexDIA of bulk samples has shown that the increased interference results in only a modest decrease in quantitative accuracy^21^ and smaller still at the single-cell level^12,23,24^. Nonetheless, the interference at higher plex has not been rigorously tested, and this motivated us to carefully benchmark the impact of nonisobaric labeling on single-cell quantification.

Advances in short separation times^25^ and algorithms interpreting data from peptides collected in parallel by LC-MS/MS (data-independent acquisition, or DIA)^26–28^ support increased throughput. Mass tags compatible with DIA can further increase throughput, but they remain limited^12,21,29,30^. Simultaneously increasing the number of protein data points measured per sample per unit time by LC-MS/MS requires engineering new mass tags that incorporate more isotopes and preserve or enhance the quality of peptide identification. Using combinatorial chemistry and high-throughput screening, we developed a mass tag, called PSMtag, with the potential to simultaneously increase the number of detectable peptide fragments and sample throughput for DIA workflows without compromising quantitative accuracy. For instance, 9-plex PSMtag allows 34,037 protein data points to be acquired over a 30-minute LC gradient, while a label-free workflow acquires 5,060 protein data points from single-cell level proteomes over the same time.

## Results

### Combinatorial synthesis

We synthesized a range of chemical structures as candidate mass tags and systematically evaluated their effect on peptide identification quality when conjugated to peptides. Using readily-available building blocks, we employed on-peptide split-pool compound synthesis^31^ to create hundreds of candidates. We synthesized these candidates directly on two model peptides (AVQVAH-WAWSNEK, TLESSGENLRLQK) immobilized on resin to limit the impact of synthetic yield and leftover reagents on analysis. The split-pool approach used sixteen trifunctional cores, five boronic acids, and five carboxylic acids as shown in Fig. 1a and Supplementary Fig. 1. This design allowed a combinatorial synthesis of 576 compounds (including expected products of partial synthesis) directly on the two model peptides.

**Figure 1.**
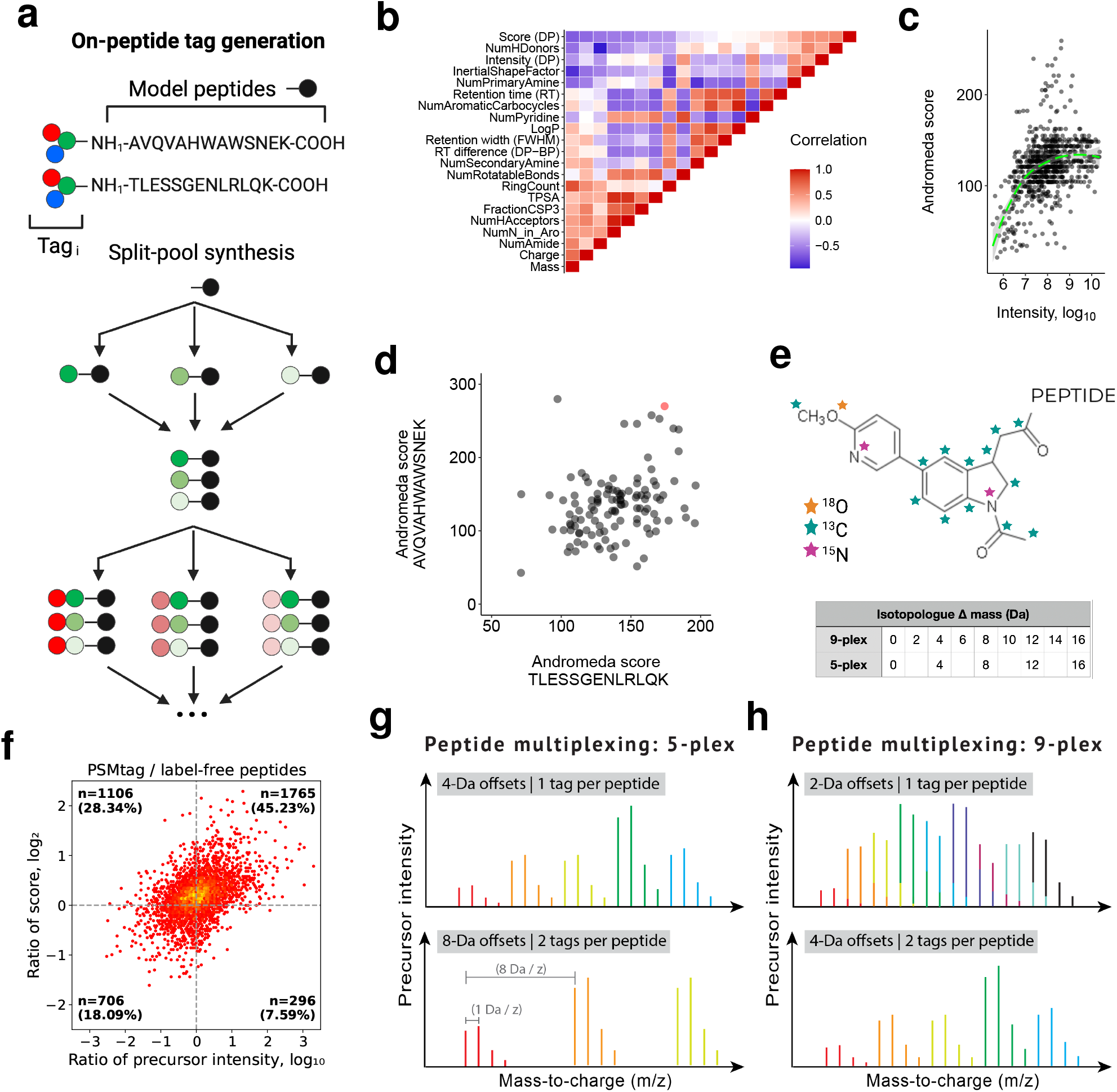
A systematic approach to find tags that improve peptide identification and throughput in proteomics workflows. **a**, Two model peptides were synthesized on resin and tag candidates were combinatorially synthesized directly on the model peptides using a split-pool approach. **b**, All pairwise correlations between the chemical and MS-measured properties of the screened compounds conjugated to model peptides. The MS properties are calculated by MaxQuant v2.7^32^ and the chemical properties are estimated via RDKit^33^ from SMILES representation of unconjugated tags. For an extended array of features, see Extended Data Fig. 1. **c** The Andromeda scores as a function of MS-measured intensity. **d**, A lead candidate was identified for further evaluation (data point colored red) based on its score on both model peptides. **e**, The structure of the lead candidate, with sites of isotopic doping of carbon, oxygen, and nitrogen indicated. A 9-plex set of reagents separated by 2 Da and a 5-plex subset of reagents separated by 4 Da were synthesized. **f**, The lead candidate was synthesized as an NHS-ester and applied to a human proteome (K-562 cancer cell line, tryptic digest) for comparison of peptide sequence identification, as measured by ratio of score (Sage^34^ hyperscore), conditioned on the ratio of precursor intensity. **g, h**, Illustration of isotopic envelopes for peptides from different samples multiplexed by PSMtag. Each color corresponds to a sample. Overlapping signal may be deconvolved *in silico* by search algorithms that model the linear superposition, such as JMod^28^.

### Compound screen for beneficial properties

The relationship between the chemical properties of tags and their effect on the LC-MS properties of tag-peptide conjugates is explored in Fig. 1b and Extended Data Fig. 1, as well as the relationship between the chemical properties of tags and holistic metrics like Andromeda score, a value summarizing the similarity between empirical mass spectra and predicted mass spectra for a given peptide sequence^35^. Andromeda scores correlate with the intensity of tag-peptide conjugates but reach an asymptote around the middle of the intensity range, Fig. 1c. The highest-scoring tag candidates are not the most intense ones (Fig. 1c). Since most labeled peptides are sampled for fragmentation near the apex of their elution profile (Supplementary Fig. 2), apex offsets in sampling are unlikely to strongly influence the Andromeda scores. Of the 576 compounds, 389 were confidently identified on at least one model peptide, while 111 were identified on both model peptides and evaluated for their effect on the confidence of matching mass spectra to peptide sequences (Fig. 1d). This confidence was measured by the Andromeda score. A lead candidate tag was identified (Fig. 1d, red) to maximize the Andromeda scores for both model peptides. We named it Parallel Squared Multiplexing tag (PSMtag) and synthesized multiple isotopologues by incorporating heavy atoms in the positions shown in Fig. 1e. These isotopologues allow for 9 tags spaced by 2 Da, and 5 tags spaced by 4 Da.

### Validation on the human proteome

To further evaluate the properties of PSMtag beyond the model peptides, we characterized its effect on a broad range of endogenous peptides from the human proteome. To facilitate further characterization, the enantiomers of the racemic synthesis of PSMtags were separated by chiral chromatography. Only enantiopure tags were used for all following analysis in this study. To reliably label many peptides, PSMtag was functionalized with NHS-ester chemistry to label peptides derived from human proteome samples. NHS-functionalized PSMtags allowed for efficient and specific labeling of N-terminal and lysine amine groups, Supplementary Fig. 3 and Supplementary Fig. 4, respectively.

Byproducts of NHS-ester labeling can be separated from tagged peptides in ion mobility and *m/z* space as shown in Extended Data Fig. 2. Some data in this study were acquired with solutions by MS manufacturers originally developed to remove similar, unrelated byproducts from proteomic analysis by ion mobility and *m/z*. These solutions include high field asymmetric wave-form ion mobility spectrometry (FAIMS) and trapped ion mobility spectrometry (TIMS). TIMS allows selective acquisition of peptide ions from tag byproducts acquiring only the signal for fragmentation and identification shown in the black boxes (”diaPASEF windows”). FAIMS effectively reduces the signal from tag byproducts several orders of magnitude as shown in Extended Data Fig. 2. Peptide ion transmission can also be impacted, but signal-to-noise improvements with FAIMS have been thoroughly discussed^36,37^.

To facilitate reliable comparisons of spectral quality, we examined the same peptide ions with and without tag while controlling for changes in their abundance (as estimated by precursor intensity). This systematic view is presented in Fig. 1f. The majority of peptides scored more favorably with the tag attached, with 73% of all peptides having higher scores when tagged across all peptide abundances. Even when the tagged peptides were apparently less abundant than their label-free counterparts, still more peptides scored higher when tagged (28% vs. 18%).

### PSMtags improve the identification of peptide spectra

#### Database-matching sequence identification

We sought to identify the mechanism increasing scores for tagged peptides compared to label-free peptides. Given that peptide sequence identification relies on detecting peptide fragments, we expected that the improved scores could be driven by increased observation of peptide fragments for tagged peptides. Tagged and label-free human proteome standards (K-562 cancer cell line, tryptic digest) were diluted to 2 ng input levels and analyzed by data-dependent acquisition on an Orbitrap Astral mass spectrometer. Tagged and label-free spectra for peptide ion FEELNADLFR (2+) exemplified an increase in Sage hyperscores^34^ for the tagged peptide ion, primarily through the stabilization of the b-ion series (Fig. 2a). Scores for both the label-free and labeled peptide allowed confident assignment of the peptide sequence to the spectra. A main reason for moving away from Andromeda scoring was that MaxQuant (by default) has stricter score cutoffs for reporting modified vs. unmodified peptides. We consider this inappropriate for comparing proteomes of tagged samples and label-free samples directly, as it would artificially filter out PSMtag peptide-spectral matches (PSMs).

**Figure 2.**
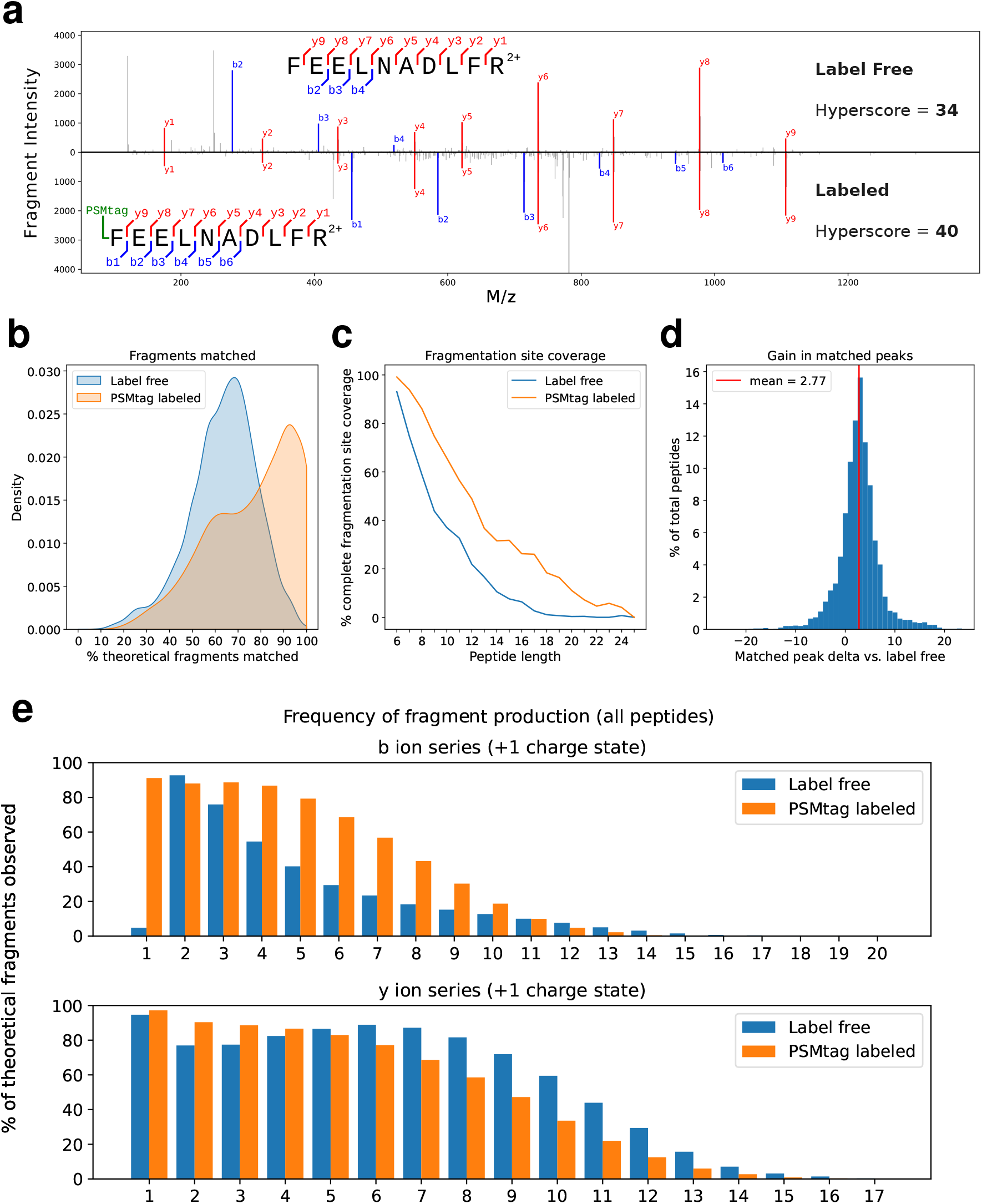
Exploring improvements to peptide sequencing. **a**, Tagged and label-free spectra for peptide ion FEELNADLFR (2+) exemplifying an increase in score (Sage hyperscore) for the tagged peptide ion, primarily through the stabilization of the b-ion series. **b**, Distribution of the percent of theoretically-expected fragments from intersected peptide sequences (n=3902) observed in their empirical spectra for both tagged and label-free samples. **c**, Percent of predicted fragmentation sites covered for tag (n=13888) and label-free (n=14748) spectra conditioned on peptide length. **d**, Distribution of the relative number of peaks in individual peptide spectra (intersected) that can be assigned to theoretically-expected fragments. **e**, For all peptides confidently assigned in tagged and label-free populations, the percent of theoretical fragments observed according to their ion type: b-ions (containing N-terminus of peptide) and y-ions (containing C-terminus of peptide). The number series indicates the number of amino acids in that ion. Data for all comparisons was acquired on the Thermofisher Astral platform using collision energies and FAIMS CV values optimized to maximize peptide identifications from the different sample types.

To systematically characterize the effect of the tag on peptide spectra, we examined the percent of theoretical fragments expected per peptide that were observed empirically in the tagged and label-free spectra for the same peptides in Fig. 2b. The tagged peptides exhibited greater representation of expected fragments. Indeed, this allowed for greater overall fragmentation site coverage across all peptide lengths, as shown in Fig. 2c. We conditioned on peptide length for this examination because longer peptides have more possible fragments. For the set of peptides confidently assigned from both tagged and label-free samples, tagged peptides had an average of 2.77 more observed fragments per spectra annotated to an expected fragment (Fig. 2d). A primary difference in spectra between tagged and label-free peptides was the greater rate of observation of b-ions for tagged peptides, as shown in Fig. 2e.

#### *De novo* sequence identification

We also sought to determine if the mechanism increasing scores for database-matching approaches to peptide spectra identification would extend to modern *de novo* sequencing approaches^38^. One potent application of *de novo* sequencing is the effective definition of a minimal non-canonical search space in immunopeptidomics^39^ to directly inform T-cell antigen discovery^40^. This approach achieves sensitive and confident identification of cryptic peptides without requiring next-generation sequencing data, which is usually required in the field^41,42^. Restricting the canonical search space for increased sensitivity has also been successfully applied to immunopeptidomics data from mono-allelic cell lines^43^. This indicates the potential for *de novo*-sequencing to tackle large search spaces which pose a challenge in immunopeptidomics^44^.

In contrast to tryptic peptides, immunopeptides are short and typically don’t bear basic, positively charged C-termini. This leads to poorly visible fragments that make MS2 spectra challenging to *de novo* sequence. We evaluated the possibility of enhanced *de novo* sequencing through PSMtag with peptides produced by GluC and ArgC, which, for analytical purposes, model the class I immunopeptidome well. For this reason, we chose proteomes digested by these alternative proteases to compare the efficacy of *de novo* algorithms to identify label-free and PSMtagged spectra.

A set of high-confidence peptide sequences in the samples was determined by taking the peptide sequences identified by a database matching approach for each protease-digested proteome, both labeled and label-free (Sage search engine, q-value *<* 0.01). This set of high-confidence peptide sequences for each sample type and their intersect is depicted in Fig. 3a,b. We used the *de novo* search engine PointNovo^45^ to identify sequences from the same samples that were processed by database matching. PointNovo models for label-free and PSMtag-labeled spectra were generated using equally sized training sets of validated PSMs (50,000 training, 5,000 validation) selected from tryptic DDA data collected on the Orbitrap Astral. The numbers of peptide spectra that were assigned identical sequences by both *de novo* sequencing and database search for each sample type are shown in Fig. 3a,b in dashed lines for different peptide length bins. To control for labeling-specific sampling effects on the peptide spectra acquired, peptide-spectral matches were additionally subsetted to include only peptides identified by database search in both labeled and label-free samples. Numbers of matching *de novo* and database-search sequence assignments for this intersected peptide subset are shown in solid lines. When not restricting sequences to those identified both labeled and label-free, more short (*≤* 9 AA) and long (*≥* 14 AA) peptides were identified *de novo* by application of PSMtag for both protease digests. For the GluC digest, more medium-length peptides were observed to correspond to database results. The subset of sequences from the intersect of both labeled and label-free database results showed more similar rates of correspondence, but the quantity and quality of *de novo* sequence assignments were improved for both intersected and non-intersected comparisons, as indicated in Fig. 3c,d. Similar to the mechanism improving database-matching results, the improvements are likely driven by increased b-ion production, as visualized in Fig. 3e,f. The relative gain in accuracy for individual amino acid assignments increased with proximity to the N-terminus, where a PSMtag is conjugated. Results were qualitatively similar using a different *de novo* search engine, Novor^46^ via novor.cloud, shown in Extended Data Fig. 3.

**Figure 3.**
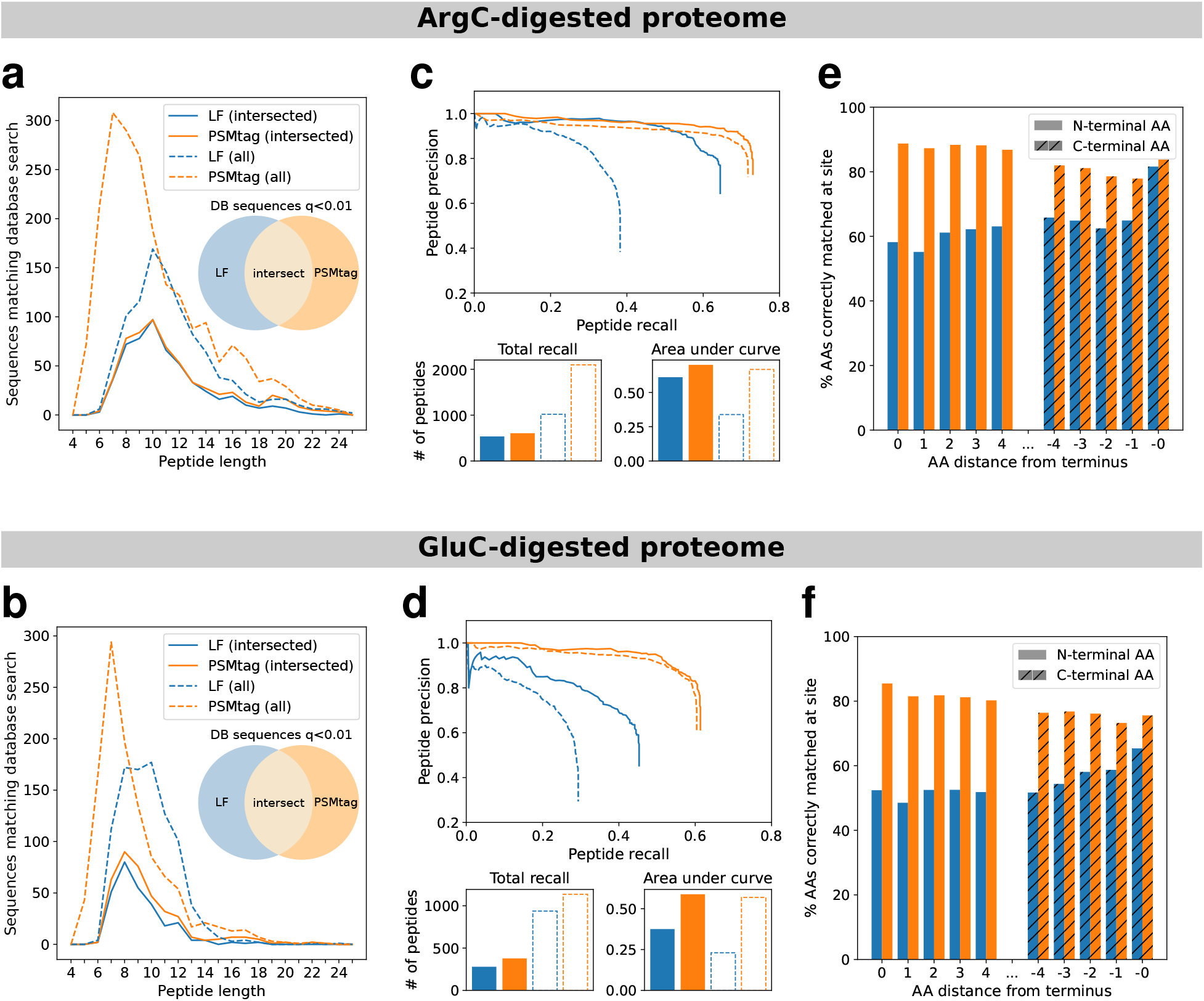
Improvements to PointNovo *de novo* sequencing with ArgC- and GluC-derived peptides. **a, b.** Total numbers of correct *de novo* peptide-spectral matches conditioned on peptide length. PointNovo sequence assignments in agreement with validated Sage PSMs (spectrum q-value *<*0.01) are considered correct *de novo* matches. **c, d.** Peptide-level precision-recall curves for tagged and label-free samples with measurements of total numbers of peptides recalled and area under each precision-recall curve. **e, f.** Overall accuracy of *de novo* amino acid assignments for residues with a distance of up to 4 AA from the peptide N- or C-terminus. Amino acids in the *de novo* sequence are considered to be correctly assigned if the database sequence contains a counterpart amino acid within 0.02 Da in mass with a prefix mass (sum of all preceding amino acids) within 0.1 Da.

#### Descriptive statistics of DDA spectra

Data-dependent workflows for bottom-up proteomics aim to acquire spectra of peptides individually. This allowed for the comparison of spectra for the same peptides tagged and label-free in Fig. 2. Overall, these low-input samples produced comparable numbers of peptide-spectra-matches (PSMs), unique sequence assignments, and proteins supported by the unique sequence assignments as shown in Extended Data Fig. 4a, whether tagged or label-free. The counts are broken down by sequences containing at least one arginine or sequences containing at least one lysine. This breakdown was chosen to support the investigation of the different effects on spectral quality induced by the conjugation of one tag (most arginine-terminating peptides) versus two tags (most lysine-terminating peptides). Small differences between the count statistics for arginine- vs. lysine-terminating sequences were observed for this data acquired by a DDA workflow. The distribution of scores (Extended Data Fig. 4b) was likewise comparable with some benefit afforded by doubly-tagged sequences. The effect on peptide chromatographic retention time of applying one versus two tags was significant and depicted in Extended Data Fig. 4c. DDA workflows, because they do not rely on predicting peptide chromatographic retention time, are well-suited to account for changes in peptide retention time without modification. Along with differences in retention time, the specific peptides yielded from the same sample by tagged and label-free preparations were different. The consistency of peptides and proteins observed in replicate label-free analyses is shown in Extended Data Fig. 4d,e. To account for the different fragmentation energy parameter used for the tagged peptides, the label-free replicates were acquired with different normalized collision energies on the Orbitrap Exploris platform (NCE = 24, NCE = 28), one determined to maximize label-free PSMs (NCE = 28) and the other determined to maximize tagged PSMs (NCE = 24). Even accounting for the different collision energies, the populations of tagged and label-free peptides and proteins were less similar, as shown in Extended Data Fig. 4f,g. Little off-target and other labeling products of PSMtag were found, as shown in Supplementary Fig. 4 and Extended Data Fig. 5a, respectively. The relative amino acid frequencies of confidently identified peptides in the label-free vs. PSMtag data was also examined in Extended Data Fig. 5b and found to be similar. The combination of tagged and label-free replicate preparations may allow greater protein sequence coverage and proteome coverage than label-free replicate preparations of the same input.

### Proteome coverage and plexDIA multiplexing with PSMtags

#### Isotopic doping for increasing throughput

We sought to maximize the number of samples that could be isotopically encoded by PSMtag without constraining the cycle of data acquisition or undermining protein quantitation. Based on previous observations, we had a strong expectation that encoding separated by 4 Da or more would allow this^21^. Furthermore, 2-Da multiplexing has been demonstrated for data-dependent workflows^14^, so we aimed to enable that encoding for plexDIA^28^ and take advantage of its greater multiplexing. The structure of PSMtag with the implemented sites of isotopic doping is shown in Fig. 1e. The sites of isotopic doping allowed the creation of a set of nine reagents separated by 2 Da. Using a subset of those reagents separated by 4 Da allowed multiplexing five samples, an increase over the state of the art (three) for plexDIA of proteomes digested by trypsin. The masses of the reagents are precisely described in Extended Data Fig. 7. The retention time consistency of the reagents is depicted in Extended Data Fig. 8, where the +16 Da reagent, the only reagent doping the pyridine nitrogen, is observed to elute earlier than other reagents.

Tags are conjugated to peptides in a sample-specific but not peptide-specific manner. Stated another way, all the peptides in a given sample are labeled by the same tag. The naturally-occurring isotopic envelope (attributable largely to natural ^13^C abundance in bio-molecules) for a given peptide is shown in Fig. 1g,h with offsets induced in the *m/z* domain by sets of tags, allowing the encoding of 5 or 9 samples, respectively. When the mass offset contributed by the tags is smaller than the width of the isotopic envelope of a peptide, the isotopic peaks of neighboring channels partially overlap. This overlap can be modeled as a linear superposition and deconvolved *in silico*. More specifically, the 9-plex described in Fig. 1h induced spacing of 2 Da / z for peptides conjugated to one tag, spacing of 4 Da / z for peptides conjugated to two tags, and 6 Da / z for three, *etc*. Enzymes commonly used in proteomics allow attaching one or two tags per peptide (trypsin) or at least two tags per peptide (LysC). Peptides resulting from incomplete cleavage by either enzyme may allow three or more tags to be conjugated.

### Evaluating identification rates and quantitation quality

Next we evaluated the amino acid sequence identification of PSMtagged peptides analyzed by DIA and throughput gains realized by PSMtags used for plexDIA. Sequence identification by DIA is highly dependent on having spectral libraries that faithfully represent peptide retention times and fragmentation patterns. Such spectral libraries are available for label-free analysis but not for PSMtags. Thus, we used DIA data of PSMtagged peptides to train fragmentation and retention time prediction models for PSMtags with DIA-NN v2.5.1^26,47^. Deeper coverage of the tagged proteome will allow training more accurate models, creating more accurate spectral libraries and better interpreting DIA spectra from peptides labeled with PSMtags.

To rigorously compare sequence coverage for single-cell level samples, we analyzed 200-pg samples from K-562 cells either as label-free samples or labeled with PSMtags using data-independent acquisition on the Bruker timsTOF SCP and Thermo Fisher Orbitrap Astral mass spectrometers. Yeast proteome digests were also spiked in at amounts described in Fig. 4f to allow for simultaneous evaluation of quantitation quality. Only human peptides and proteins were considered for evaluation of identification rates in Fig. 4a-d. Sample D was used for label-free and single-channel evaluation. Protein FDR was computed separately within each sample type (label-free, single-plex, 5-plex and 9-plex).

**Figure 4.**
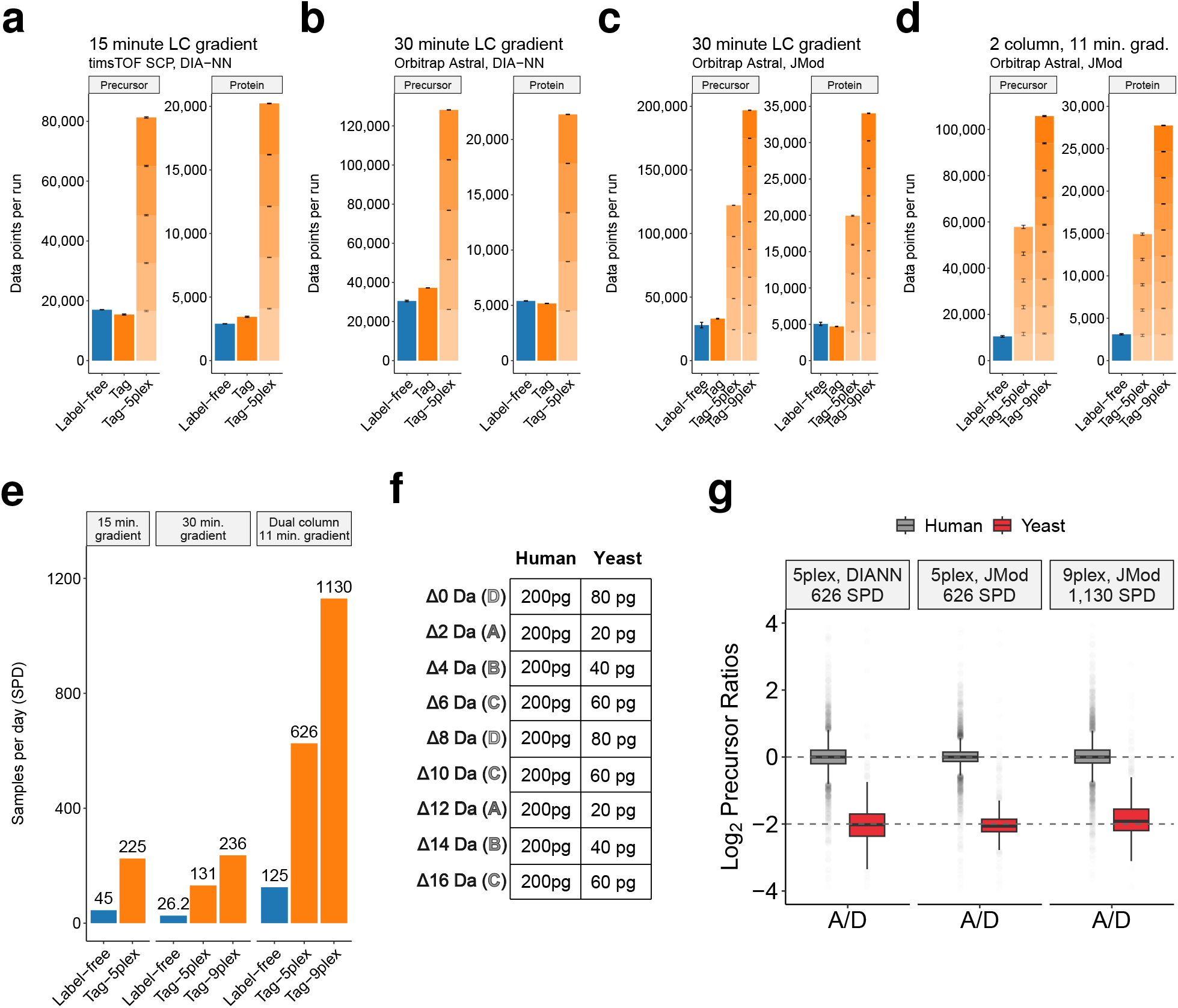
Identification rates for tagged and label-free single-cell-level (200 pg) proteomes by DIA. **a-d**, Numbers of human precursors and proteins identified in replicate injections of samples containing 200 pg human proteome (K562 cancer cell line) and yeast proteomes spiked as described in **f**. Samples were unlabeled, labeled with a single PSMtag isotopologue, or with the 5-plex PSMtag set and multiplexed. Label-free and single-plex samples used sample D diluted to 200 pg of human material. Plexed samples were diluted to 200 pg of human material per channel. All precursors and proteins are reported at 1% FDR. **a**, Data acquired using timsTOF SCP and analyzed with DIA-NN 2.5. **b**, Data acquired with Orbitrap Astral and search with DIA-NN 2.5. **c**, The data from **b** plus a 9-plex set were searched with JMod. **d**, Data acquired using Orbitrap Astral and a dual column setup with 11-minute gradients^25^ and searched with JMod. **e**, The number of samples per day (SPD) that can be analyzed using each approach. Full run-to-run time, including sample loading, active gradient, wash, and equilibration steps, is considered in the calculation. **f**, The experimental design for samples in this figure, prepared in bulk quantities and diluted to 200 pg of human input and 20-80 pg of yeast input. **g**, The measured versus expected ratio for yeast spike-ins in samples comprising 200 pg of human proteome and 20-80 pg of yeast proteome at 626 and 1,130 SPD. More ratios also are examined in Extended Data Fig. 6. Precursors intersected across all three conditions visualized.

When searched with DIA-NN^26^ or JMod^28^ using libraries fine-tuned as described^47^, singly-tagged, arginine-terminating peptides had similar or improved rates of identification, while the identification rate for doubly-tagged, lysine-terminating peptides was preserved or diminished, Extended Data Fig. 9a,b,c. Generally, the number of confident peptide and protein identifications was maintained or increased relative to label-free counterparts when considering a single tag applied to single-cell level proteomes, and coverage by sample decreased slightly in the plexed samples. Specifically, as shown in Fig. 4a, the number of confident protein data points per run identified by DIA-NN 2.5 from data acquired using a 15 minute gradient on the timsTOF SCP platform is 2,901 when label-free, 3,444 when using a single PSMtag, and 20,228 when using 5-plex PSMtag. As shown in Fig. 4b, the number of confident protein data points per run identified by DIA-NN 2.5 from data acquired using a 30 minute gradient on the Astral platform is 5,400 when label-free, 5,173 when using a single PSMtag, and 22,268 when using 5-plex PSMtag. As shown in Fig. 4c, the number of confident protein data points per run identified by JMod 2.0 from data acquired using a 30 minute gradient on the Astral platform is 5,060 when label-free, 4,696 when using a single PSMtag, 19,944 when using 5-plex PSMtag, and 34,037 when using 9-plex PSMtag. As shown in Fig. 4d, the number of confident protein data points per run identified by JMod 2.0 from data acquired using a 11-minute gradient on the Astral platform is 3,216 when label-free, 14,868 when using 5-plex PSMtag, and 27,520 when using 9-plex PSMtag. The applied gradients demonstrated a number of different throughputs. The 5-plex tags demonstrated a 225 samples-per-day (SPD) rate on 15 minute gradients and the 9-plex tags demonstrated a 236 SPD on 30 minute gradients that allow greater depth of proteome coverage, as shown in Fig. 4e. SPD of 1,130 was achieved using a dual column setup^48^ and 11-minute gradients with 9-plex tags.

The quantitation accuracy achieved by plexDIA using PSMtags was benchmarked using the proteome mixing ratios described in Fig. 4f and found comparable to that achieved by matched label-free analysis of the same samples, as shown in Extended Data Fig. 6a and Fig. 4g. The results indicate that PSMtags can achieve high accuracy and precision across a range of mixing ratios, Extended Data Fig. 6b. Quantification with PSMtag of larger mixing ratios, up to 1:32, was benchmarked by McDonnell *et al.*^28^ These results indicate that PSMtags increase throughput without significant adverse impact on depth of coverage or quantitative accuracy.

#### Predicting spectral libraries for new mass tags

While spectral libraries can be well predicted for unlabeled tryptic peptides^49,50^, tagging peptides alters their properties, including retention time (RT) (Fig. 5a-c), fragmentation spectra (Fig. 5d) and ion mobility (IM) (Fig. 5e) for the corresponding predictions of the default models trained on label-free data. These differences can be reduced by fine-tuning the models, which is increasingly integrated into leading DIA search engines^26,47,51^ by using empirical DIA data from tagged peptides. Here, we used DIA-NN to fine-tune the models and predict improved spectral libraries as previously described^47^. Specifically, we used a random 50:50 training-validation split of PSMtag labeled peptides (stripped sequences) from single replicate runs on Orbitrap Astral or timsTOF SCP instruments. Evaluating the tuned predictions on the validation set indicates significant improvements. Before tuning, the default label-free model predicted the RT of PSMtag peptides poorly (Fig. 5a), with large iRT errors whose direction differed between K- and R-peptides (Fig. 5c). Tuning brought predicted RT into close agreement with empirical values (Fig. 5b,c), albeit RT deviations remain larger than for unlabeled peptides. Tuning likewise increased the spectral angle similarity between predicted and empirical spectra (Fig. 5d) and removed the systematic IM errors (Fig. 5e). These improvements in the predicted spectral library increased the number of precursors and proteins identified on both instruments (Fig. 5f). Despite these substantial improvements, libraries predicted for PSMtag peptides remain less accurate than those for unlabeled peptides: Even after tuning, PSMtag spectra have lower spectral angle similarity, and the RT and IM errors are more dispersed, i.e., differ more from empirical measurements (Fig. 5c-e), leaving space for further improvements in the predictions, which may further increase peptide sequence identification for labeled peptides.

**Figure 5.**
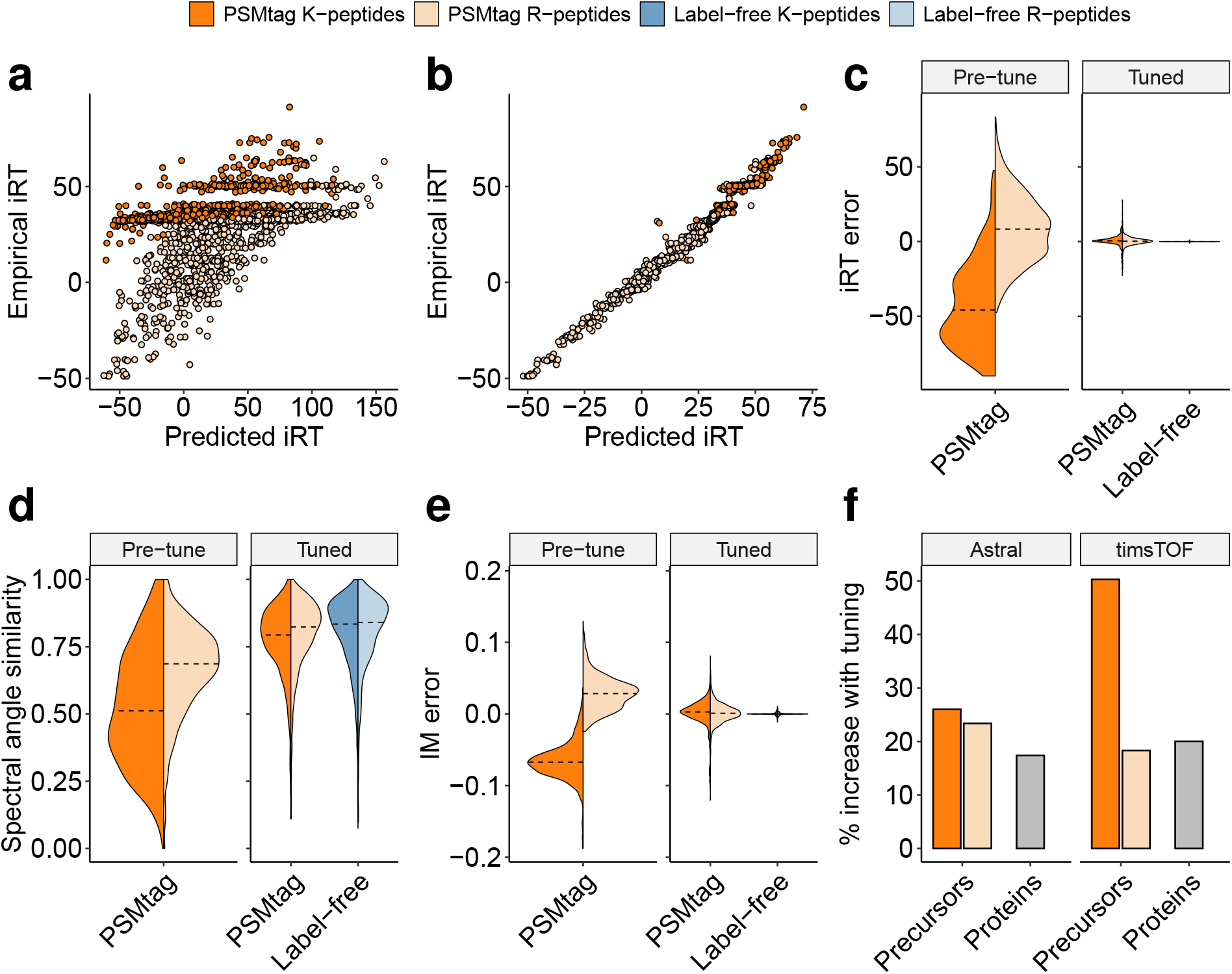
Tuning spectral libraries for PSMtag identification. **a**, The predicted and empirical standardized retention times (iRT) for a single PSMtag run. Predicted values are generated with the internal DIA-NN default label-free model. Data acquired using the timsTOF SCP and analyzed with DIA-NN 2.5.1. For panels **a-f**, distributions, data points, and bars are colored by labeling state (PSMtag or Label-free) and whether the given precursor contains lysine (K-peptides) or not (R-peptides). **b**, The same empirical values shown compared to values predicted by DIA-NN 2.5.1 following a published tutorial.^47^ **c**, The distribution of standardized retention time errors pre- and post-tuning. Training and validation performed on a random 50:50 split of stripped peptide sequences. Comparisons on tuned values for **c-e** use the validation set only. **d**, The distribution of spectral angle similarity (as defined by Gessulat *et al.*^49^) of pre- or post-tuned spectra to their empirical counterparts. Data acquired using the Astral. **e**, The distribution of ion mobility (IM) error. Data acquired using the timsTOF SCP. **f**, The percent increase above the default DIA-NN 2.5.1 label-free model for tuning libraries independently for Astral and timsTOF data.

#### Evaluating in-plex coverage and quantitation across diverse cell types

Most applications of multiplexing proteomics samples involve comparisons of a given species across different experimental conditions, which we aim to capture with three diverse but common human cell lines (K562, HeLa, U937) in the experimental design for Fig. 6a. To evaluate data completeness, we obtained replicate measurements for each cell line using either label-free DIA or 9-plexDIA and estimated the intersection and union of proteins across the 3 replicates x 3 cell lines for the label-free and labeled experiments, Fig. 6b,c. The union of proteins for the label-free approach across the 9 total samples is larger (9,439) than the union for the 9 labeled samples (6,262). The intersect of proteins (those identified in all samples) for the 9-labeled samples is larger (5,996) than the intersect for the label-free samples (4,887). Displayed in Fig. 6d, the number of protein data points measured per run for the 9 label-free samples has a mean of 6,962, while the number of protein data points measured per run for the 9 labeled samples is 56,358. The protein-level coefficient of variation (%) is calculated for proteins observed in all three replicates (n=3) for a given label-free cell type or a given label channel. The distribution is shown in Fig. 6e. The labeled samples show longer tails, leading to more proteins with CV above 25% than the label-free samples.

**Figure 6.**
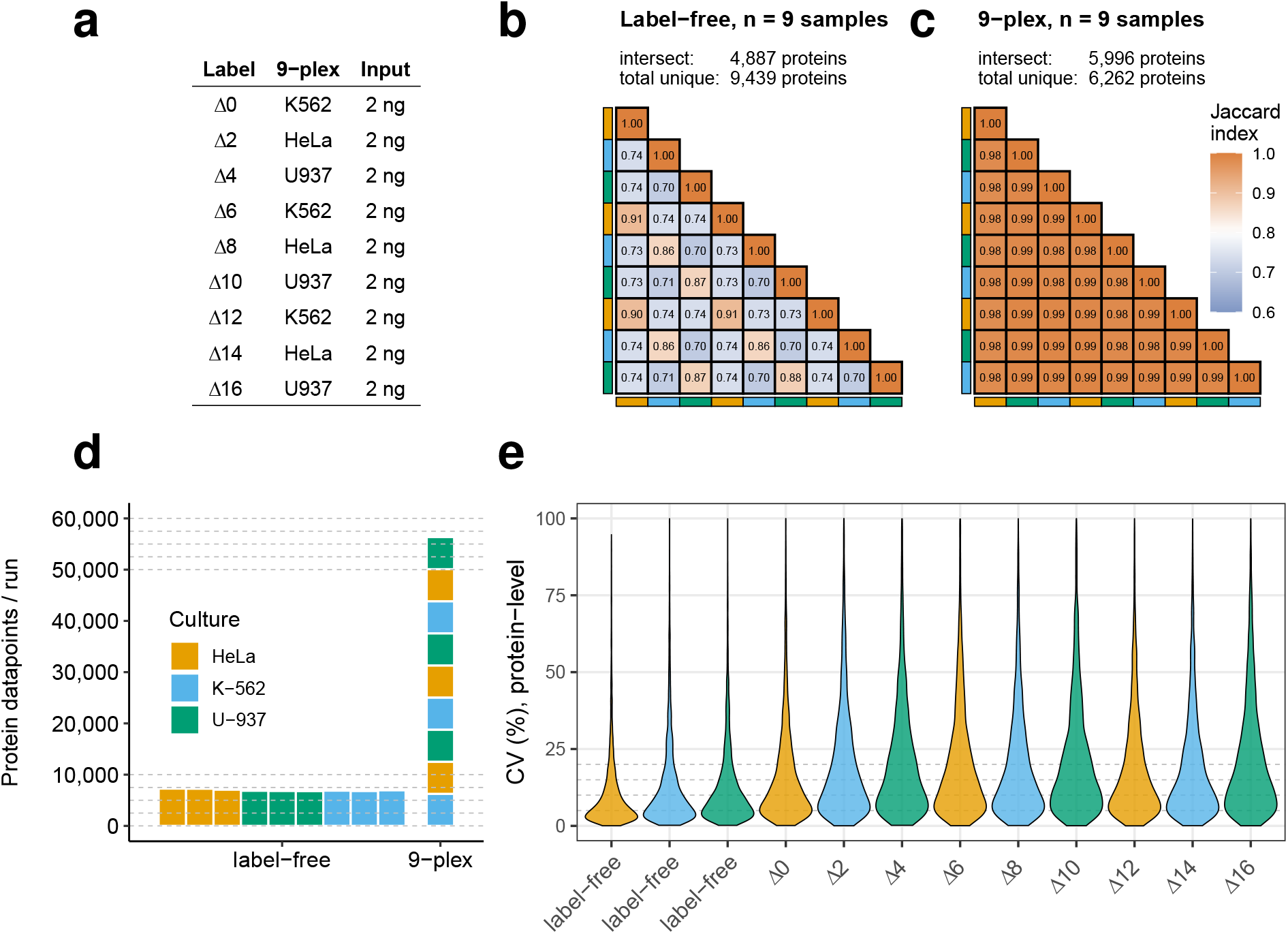
Evaluating protein coverage and quantitation across diverse cell types. **a**, The experimental design: K-562, HeLa, and U-937 cell lines were lysed, digested, labeled, cleaned up with magnetic amine beads, combined at 2ng per channel and analyzed as described in the Methods. Similarly, 3 replicate measurements of each of the same 3 cell lines label-free at the same input level of 2ng were also taken. All data was acquired on Orbitrap Astral and searched with JMod using a fine-tuned library generated by DIA-NN. Nine label-free runs are compared to one run or 9-plex PSMtag unless otherwise stated. **b**, The Jaccard indices of each pairwise comparison of protein sets confidently identified (1% FDR). The intersect and union of proteins confidently identified across the 9 label-free samples. **c**, The Jaccard indices of each pairwise comparison of protein sets confidently identified (1% FDR). The intersect and union of proteins confidently identified across the 9 labeled samples. **d**, The confident protein data points per run per cell type per labeling condition. **e**, The protein-level coefficient of variation (CV, %) calculated requiring three replicates per channel or per label-free cell type. For the labeled samples, this involves three independent runs to obtain three replicates per individual channel. For the label-free samples, this involves 9 independent runs to obtain three replicates per cell type. Less than 0.5% of data points across all samples in the plot area are above 100% and thus not displayed.

## Discussion

We present what may be the first systematic approach to discovering and validating new mass tags for bottom-up proteomics by LC-MS/MS. From an initial screen of hundreds of compounds (Fig. 1a), we select and demonstrate a new mass tag (Fig. 1e) that improves peptide identification (Fig. 1f) and throughput (Fig. 4e) for sensitive proteomics applications. Specifically, the 5-plex and 9-plex sets of reagents allow for accurate quantification of 14,868 and 27,520 protein datapoints in the same time a label-free experiment analyzes 3,216 protein datapoints (Fig. 4d).

The detection of peptide fragment ions is improved by tag conjugation, as demonstrated in Fig. 2b,c and evident when controlling for both protein input amount and peptide ionization (Fig. 1f). These improvements are driven by stabilization of b-ions (Fig. 2e for both lysine- and arginine-terminating peptides, Extended Data Fig. 10, and higher charge state fragments, Supplementary Fig. 6). We detail possible mechanisms for b-ion stabilization in Supplementary Note 1.

The benefits for sequencing short, non-tryptic peptides with PSMtag is consistent with previous findings of enhancing sensitivity for immunopeptidomics with labeling strategies^52^. Thus, PSMtag can extend these tagging benefits to DIA analysis of non-tryptic peptides, such as immunopeptidomics.

### Exploring chemical properties correlated with positive tag performance

One goal of rational mass tag design is to understand and subsequently leverage the relationship between a sufficient subset of chemical properties/descriptors of small molecules and their effect on peptide fragmentation. In Fig. 1b and Extended Data Fig. 1, we present relevant chemical descriptors and LC-MS metrics across the observed tag candidates. Due to high interdependence of these variables, and the comparatively low number of candidates measured relative to the number of these variables, we suggest the relationships revealed by the data in this study be used for hypothesis generation to be tested in further studies. We discuss more limitations and opportunities in rational tag design in Supplementary Note 2.

### Further potential for multiplexing with isotopic doping

Not all possible sites for stable isotopic incorporation for PSMtag were used due to budget and time constraints. Additionally, this study restricted itself to multiplexing using peptides derived from biological samples using the enzyme trypsin to digest proteins due to its near universal adoption by the bottom-up proteomics field. However, less-frequently used enzymes, such as LysC and LysN^12,13^, create multiple sites per peptide for tag conjugation, thus requiring fewer isotopes to be incorporated to differentiate between reagent channel. Deuterium incorporation was not considered to achieve multiplexing in this study, but recent advances start to explore its promise and limitations^10^ for use in the context of mass tags. Even without deuterium incorporation, 1000-plex mass tags may be designed by incorporating the differential mass defect to create mDa differences between plex channels^53^, but full realization requires sufficiently-high MS resolution.

### Benchmarking single-cell standards at high SPD

Single-cell proteomics is an application area demanding high throughput that can participially benefit from PSMtags^54^. Using mixed species standards, we find that 9-plex PSMtag are compatible with high accuracy and precision at single-cell input (Fig. 4g, Extended Data Fig. 6c), but also conclude they are marginally more susceptible to noise and ratio compression in an intensity-dependent manner than 5-plex or label-free approaches (Extended Data Fig. 6c,d). This intensity-dependence of accuracy and precision is observed in other studies using the mixed species standards^48,55^. Controlling for precursors in PSMtag and label-free samples, some degree of ratio compression and increased variance can be observed for the 9-plex samples run at 1,130 SPD. This trade-off in precision and accuracy can be weighed against the increased throughput. Crucially, the reduced precision for the 9-plex is relative to methods with lower throughput, and we could not compare to 1,130 SPD label-free method as such methods have not been implemented.

### Limitations and opportunities in discovering new mass tags

#### This first screen represents a limited range of properties

The 576 compounds studied represent a limited range of chemical properties, for example: logP (Supplementary Fig. 7). While the screen resulted in at least one compound to improve peptide identification, the quenched and hydrolysis products of the compound (logP approx. 1.5) coelute with peptides on standard C18 reverse-phase chromatography (Extended Data Fig. 2c). While the two compounds are chromatographically well-resolved and omitted from accumulation in the mass spectrometer by nature of being lower *m/z* than most tagged peptides, the peptides with which they coelute may experience ionization suppression. The degree to which this affects peptides is mostly dependent on their relative abundance: highly abundant ions may tolerate ionization suppression without loss of sequencing or quantitative accuracy, while less abundant ions will experience more deleterious effects. Additionally, both model peptides contain lysine at the C-terminus. A more comprehensive screen would increase peptide diversity.

### Impact of heavy isotope doping on retention time

Peptide analytes incorporating ^15^N can elute earlier than the same peptide analytes incorporating ^14^N^56,57^. The broad adoption and utility of multiplexing reagents which incorporate ^15^N motivated approaches to accurately quantify peptide analytes with such isotopic enrichment^58–60^. Similarly, isotopologues of PSMtag with ^15^N incorporated into the indoline-like portion of the structure show some elution time differences in Extended Data Fig. 8a, while the isotopologue (Δ16) with ^15^N incorporated in the pyridine ring structure shows a larger effect. The effect size is also larger with two tags per peptide than with one, as shown in Extended Data Fig. 8b,c. Future tag designs^53^ can consider the incorporation of ^15^N at moieties that induce minimal elution time differences, in turn, minimizing the number of coeluting peptide peaks.

### Removal of byproduct and leftover tag reagent prior to LC-MS analysis

Minimization or removal of tag byproducts is desirable. We took the approach of minimizing PSMtag byproduct by using anhydrous peptide labeling conditions to reduce reagent hydrolysis during labeling and thus allow less reagent. Additionally, ion mobility separation of the byproduct from the peptides (Extended Data Fig. 2b-e) has been used to achieve some results in this study, but not all mass spectrometers are equipped with such ion mobility separation devices (i.e. FAIMS or TIMS). For mass spectrometers with or without ion mobility separation, we introduce a magnetic bead-based clean-up for removing unreacted (NHS-ester-functionalized) tag from solution prior to LC-MS analysis to allow the super-stoichiometric amount of tag to be isolated from peptides by means other than ion mobility. While this approach was fruitfully applied in Fig. 6, optimization of reaction conditions for the range of inputs commonly used in bottom-up proteomics is outstanding. We characterize and discuss the performance of the amine-bead quenching protocol for removal of NHS-ester PSMtag and recovery of PSMtag-labeled peptides in Supplementary Note 3.

### Conclusion

With the first screen of 576 compounds, we uncovered insights into characteristics that may lend to improved LC-MS sequencing. Such insights provide guidance for designing new compounds and interrogating new chemical spaces, enabling a cycle of discovery that will allow strides to proteomics depth and throughput in parallel.

## Acknowledgments

PTI is a Convergent Research Focused Research Organization (FRO) and has received support from Eric and Wendy Schmidt as well as Griffin Catalyst. We thank Jason Derks for helpful discussions and advice on plexDIA analysis and we thank Jane Nagel for unifying language in the Methods section. We thank engineers Rob Thomas, Mark Trahan, Vlad Ondruska, John Draper, Michael Krawitzky and Chase Laxdal for maintaining LC-MS instrumentation at top performance.

## Data availability

Data, raw and processed, for reproducing figures is available through data repositories MassIVE MSV000097968, MSV000103730 and ProteomeXchange PXD064191, PXD086351. Processed data organized by contributing figure is available at data repository Zenodo under DOI https://doi.org/10.5281/zenodo.23675860.

## Code availability

The code used for data analysis and figures is at: github.com/ParallelSquared/PSMtag

## Competing Interests

H.S., M.Y., S.S., D.B.S., M.A., W.V., A.Y., M.A.L., J.S.B., B.P. and N.S. are listed as inventors on a patent application for the tags described in this paper. R.T.K. have a financial interest in MicrOmics Technologies, whose LC workflow was used for the 1130 SPD demonstration. PTI has a collaborative research agreement with Thermo Fisher Scientific, the manufacturer of mass spectrometry instrumentation used in this research.

## Author contributions

**Experimental design**: H.S., M.Y., S.S., B.P. and N.S.

**Data analysis**: H.S., M.A., K.M., S.S., M.Y., H.R., J.J.A. and N.S.

***De novo* data analysis**: M.A.

**LC-MS/MS**: S.S., M.Y., C.F., C.W. and R.T.K.

**Isotopic incorporation route design:** D.B.S.

**Compound synthesis**: D.B.S., W.K.P., Y.H., K.S.R., W.V., A.Y., J.S.B. and M.A.L.

**Proteomic sample preparation**: S.S., M.Y., M.P.A., H.R. and J.J.A.

**Raising funding**: N.S. and H.S.

**Writing**: H.S., S.S., M.A., M.Y. and N.S. wrote the first draft.

All authors approved the final manuscript.

## Methods

### Model peptide synthesis

AAs were randomly selected, with replacement, from biological distribution of AAs and terminated with lysine for ease of coupling to resin for synthesis while retaining similarity to tryptic peptides. Model peptide length (13 AAs) is longer than the median tryptic peptide length (10-12) to balance opportunities for mass tag candidates to enhance peptide fragment production with producing the minimum number of fragments for consistent peptide spectral matching.

Model peptides were synthesized via manual flow solid phase peptide synthesis (SPPS) as previously described^61^. For each peptide, this involved the use of 500 mg of NH_2_-Tentagel resin (500 mg, 0.23 mmol/g loading, 300 *µ*m monosized). The sequences synthesized using this resin consisted of either:

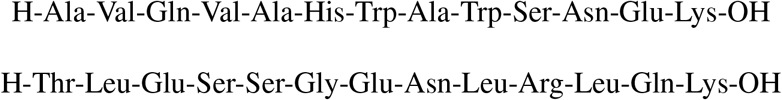

The resin was first prepared and functionalized for use via attachment of a 4-(4-hydroxymethyl-3-methoxyphenoxy)-butyric acid (HMPB) linker and Fmoc-Lys(Boc)-OH (fluorenylmethoxycarbonyl and tert-butoxycarbonyl protecting groups, respectively) as follows: NH_2_-Tentagel resin (500 mg, 0.23 mmol/g loading, 300 *µ*m monosized) was initially swelled using N,N-dimethylformamide (DMF) in a fritted Torviq syringe. Next, a solution consisting of HMPB (276.28 mg, 0.575 mmol, 5.0 equiv) dissolved in a hexafluorophosphate azabenzotriazole tetramethyl uronium (HATU) solution (0.38 M in DMF, 2.878 mL, 4.75 equiv) and activated with diisopropylethylamine (DIPEA) (0.601 mL, 15.0 equiv) was added. The reaction mixture was allowed to react for 1 hour prior to draining, and the resin was washed with DMF (3 x 10 mL). Next, a solution of Fmoc-Lys(Boc)-OH (538.82 mg, 1.15 mmol, 10 equiv) and DMAP (1.405 mg, 0.0115 mmol, 0.1 equiv) in DMF (2.875 mL) was added to the resin, followed by addition of N,N’-diisopropylcarbodiimide (DIC) (90.03 *µ*L, 0.575 mol, 5 equiv). The syringe was then capped and allowed to mix overnight on a nutating mixer. The solution was subsequently drained, and the resin washed with DMF (3 x 10 mL).

After resin preparation, peptide synthesis was conducted in the following manner: The remaining residues (as appropriate based on sequence) were sequentially coupled in a C-to-N direction via flow SPPS, which included an initial deprotection step to remove the Fmoc group from Lys(Alloc) (Allyloxycarbonyl protecting group)^61^.

### On-peptide compound synthesis

Small molecule probes were attached to the free N-terminus of the peptide via procedures based on previous peptide encoded library efforts^31^. The resin batch was evenly split into 16 different fritted Torviq syringes. To each portion of resin was coupled a different trifunctional building block (10 equiv each) as detailed in Supplementary Fig. 1.

Each trifunctional building block was dissolved in a solution of HATU in DMF (0.38 M, 239.58 *µ*L, 9.58 *µ*mol, 9.5 equiv) and activated with DIPEA (50.08 *µ*L, 0.2875 mmol, 30 equiv) prior to addition to resin. The reaction was then allowed to proceed for 1 h. Afterwards, specific tubes were combined separately due to the isobaric nature of some of the building blocks. Recombination was performed based on the color coding shown in Supplementary Fig. 8, referencing the compound IDs in Supplementary Fig. 1, generating four separate mini-libraries which were then each washed with DMF (3 x 3 mL). Trifunctional building blocks A and C were omitted. Fmoc deprotection was then performed using a solution of 20% piperidine in DMF (3 x 3 mL, 5 min each) before being washed again with DMF (3 x 3 mL). Each mini-library was then individually subjected to the following procedures: Each batch of resin was equally split into five portions and transferred to smaller fritted Torviq syringes. To each portion of resin was coupled a different carboxylic acid (10 equiv each, twice) as detailed in Supplementary Fig. 1. Each carboxylic acid was dissolved in a solution of HATU in DMF (0.38 M, 143.75 *µ*L, 54.625 *µ*mol, 9.5 equiv) and activated with DIPEA (30.05 *µ*L, 0.1725 mmol, 30 equiv) prior to addition to resin. The reaction was then allowed to proceed for 1 h before being drained, and the coupling step repeated once more. The resin portions were then pooled and washed with DMF (3 x 3 mL), then THF (3 x 3 mL). For Pd-mediated cross-coupling, microcentrifuge tubes (5 x 1.5 mL) were each charged with a different boronic acid (5 equiv each) as detailed in Supplementary Fig. 1.

To each tube was added Pd G4 XPhos (5.937 mg, 6.9 *µ*mol, 1.2 equiv) and XPhos (3.29 mg, 6.9 *µ*mol, 1.2 equiv). Next, resin suspended in THF (2.59 mL) was then thoroughly mixed and evenly dispensed into each microcentrifuge tube in 12 equal portions, each containing an identical final volume of THF (517.5 *µ*L). Finally, a solution of K_3_PO_4_ (0.5 M, 57.5 *µ*L, 28.8 *µ*mol, 5 equiv) was added to each microcentrifuge tube and thoroughly mixed via pipetting. The headspace of each microcentrifuge tube was then purged with a stream of N_2_ and left to react on a nutating mixer for 24 h at 37 °C. Finally, each portion of resin was combined, washed with THF (3 x 3 mL), DCM (3 x 3 mL), DMF (3 x 3 mL), H_2_O (3 x 3 mL), DMF (3 x 3 mL), diethyldithiocarbamate (DTC) solution until original resin color restored (0.022 M DTC in DMF, prepared via 100 mg DTC per 20 mL DMF, 3 x 3 mL), and DMF (3 x 3 mL). The resin was then washed with DCM (3 x 3 mL) and dried under vacuum prior to cleavage from the resin and global deprotection, furnishing the desired peptide-bound small-molecule library.

Each batch of resin was transferred to a 50 mL conical vial and cleaved and deprotected by the addition of 5 mL of cleavage cocktail (95/2.5/2.5 mixture of TFA/H_2_O/TIPS), followed by agitation using a nutating mixer for 3 h at room temperature. The desired material was subsequently precipitated by the addition of cold Et_2_O (−78 °C, to a total volume of 50 mL) then vortexed to ensure thorough mixing. This suspension was then centrifuged at 20,800 x g for 10 minutes to yield the desired material as a pellet. The supernatant was decanted and the pellet resuspended with an additional portion of cold Et_2_O (−78 °C, 25 mL) prior to being centrifuged once more at 20,800 x g for 10 minutes. The supernatant was discarded, and the pellet containing both resin and peptide was dissolved in a solution of 50/50 H_2_O/ACN + 0.1% TFA (15 mL) prior to filtration through a fritted syringe to remove the resin. The filtrate was then frozen and lyophilized to furnish the desired material as a crude white powder.

The crude material was weighed to determine the crude mass prior to loading onto Discovery® DSC-18 SPE Tubes (aiming for sample loading of max 10% of SPE bed wt.). The peptide sample was first dissolved in a minimal volume of DMF (approx. 200 *µ*L), prior to diluting with 5/95 H_2_O/ACN + 0.1% TFA (with a final DMF content of no more than 2% v/v). SPE was then performed using the following procedure as outlined:

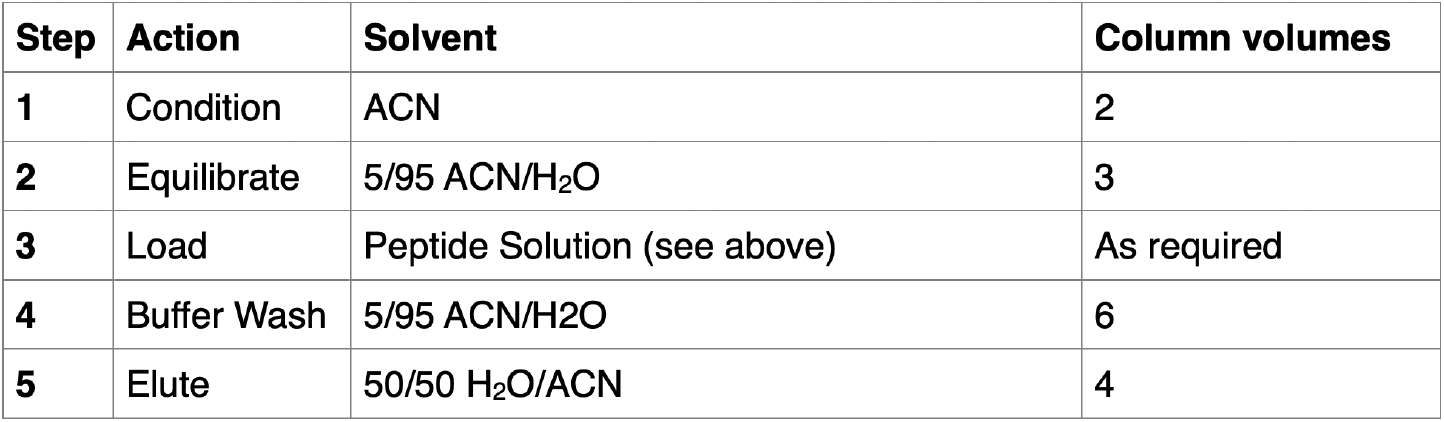

Eluted volumes were then transferred into pre-weighed 15 mL conical vials before being flash frozen and lyophilized to furnish the desired mini-library.

### Compound-peptide library data acquisition with Thermo Scientific Orbitrap Exploris 240

A pool of 576 initial compounds synthesized by split-pool chemistry were conjugated to the model peptides and separated via online nanoflow liquid chromatography (nLC) with a Thermo Vanquish Neo set in trap-and-elute mode and analyzed by a Thermo Scientific Orbitrap Exploris 240 mass spectrometer. Approximately 10 ng of the sample was loaded onto a 0.5 x 0.3 cm trapping column (Thermo Scientific, 174500) and separated with a 25 cm x 75 *µ*m IonOpticks Aurora Series UHPLC analytical column (IonOpticks, AUR2-25075C18A). Mobile phase A (MPA) was 0.1% formic acid (FA) in water and mobile phase B (MPB) was 80% acetonitrile (ACN) with 0.1% FA. A constant flow rate of 300 nL/min was used throughout sample loading and separation with a column temperature of 55°C. Samples were loaded onto the column at 6% MPB. The active gradient was ramped to 29% MPB over 23 min then increased to 44% B over 7 min. MPB increased to 99% over 0.2 min and was maintained for 4 min before returning to 6% MPB in 0.1 min for column reequilibration. Using the Exploris 240, electrospray voltage was set to 1900 V.The temperature of the ion transfer tube was 275°C. All MS1 scans were collected at a resolution of 60,000 over *m/z* 380-900 with a maximum fill time of 100 ms, and an AGC of 1e6. The RF lens was set to 80% for MS1 and MS2. Following the MS1 scan, the top 20 ions above the intensity threshold of 5e3 with charge states 2-5 were selected for MS2 fragmentation and measurement. Dynamic exclusion was used with a count of 2 and duration of 60 s with a mass tolerance of 10 ppm. For MS2 scans, ions were isolated with a 1 *m/z* window, 0.7 Da offset, and fragmented with 23% normalized collision energy (NCE) based on “Standard” AGC and “Auto” max injection time mode settings. The fragments were analyzed at 30,000 resolving power with the first mass fixed at *m/z* 120.

### Compound-peptide identification

Mass tag candidates synthesized directly onto model peptides were identified and scored by the dependent-peptide approach described by Savitski *et al.*^62^ using 10 ppm tolerance to match the expected mass of the synthesized candidates. Isobaric compound sets were acquired in separate LC-MS runs to allow for unambiguous identity assignment.

### Cell culture

Human K562, myelogenous leukemia cells (ATCC, CCL-243) were grown as suspension cultures in RPMI high glucose media (Sigma-Aldrich D5796) with the addition of 100 U/mL Penicillin-Streptomycin (Sigma-Aldrich P0781) and supplemented with 10% fetal bovine serum (FBS, Millipore Sigma F4135). U-937 (ATCC, CRL-1593.2) cells were grown as suspension cultures in RPMI medium (HyClone 16777-145) supplemented with 10% fetal bovine serum (FBS, Millipore Sigma F4135) and 1% penicillin-streptomycin (pen/strep, Thermo Fisher 15140122). Cells were passaged when a density of 10^6^ cells/mL was reached, approximately every 2 days. Pierce™ HeLa Protein Digest Standard (Thermofisher 88329) was used for experiments involving HeLa material.

### Cell harvest

Cultured cells were collected in 15 mL conical tubes by centrifugation at 500 g for 10 minutes. An aliquot of 20 *µ*L was taken before centrifugation and used for counting. The cells were washed twice with PBS in the 15 mL tube then transferred in PBS to a 1.5 mL Eppendorf tube, spun in the microcentrifuge at 500 g for 5 min then quickly washed one time with LC-MS water and spun for 2 minutes. Supernatant was removed by vacuum suction, and the cell pellet disrupted and resuspended in LC-MS water at 2000 cells/*µ*L then stored at −80°C until ready for use.

### Cell lysis and digestion with trypsin

Intact human and yeast proteins (Promega V6941 and V7341, respectively) were digested via manufacturer’s suggestions. Briefly, proteins that are presolubilized by the manufacturer (in 6.5 M urea in 50 mM Tris-HCl, pH 8 at a protein concentration of 10 mg/mL), were reduced with dithiothreitol (DTT, Pierce A39271) to a final concentration of 5 mM for 30 min at 37°C, and then alkylated using iodoacetamide (IAA, Pierce A39270) to a final concentration of 15 mM for 30 min in the dark at room temperature. The urea solution was diluted with 100 mM triethylammonium bicarbonate (TEAB, Thermo Scientific P90114) and trypsin was added at a mass ratio of 1:40 enzyme:protein and allowed to digest overnight at 37°C. The digestion was quenched to 1% FA, and desalted using C18 solid phase extraction with Oasis HLB columns (Waters WAT094225) using manufacturer’s protocol.

Frozen, harvested K562 cells were lysed and digested using the minimal proteOmic sample preparation (mPOP) method. Briefly, frozen cells taken directly from −80°C were heated to 90°C in a thermal cycler for 10 minutes to lyse. TEAB was added to a final concentration of 100 mM (pH 8.5) for buffering and Trypsin Gold (Promega V5280) was added to a final trypsin concentration of 10 ng/*µ*L, followed by digestion at 37°C for 4 h in a thermal cycler. Digested proteins were then quenched to 1% FA, dried *in vacuo* and stored until ready for future use.

### Anhydrous labeling

The following protocol has been optimized for labeling and quenching dried peptides within the range of 4-24 *µ*g. Labeling of human and yeast peptides was carried out anhydrously to minimize competition of hydrolysis with aminolysis at the primary amines of peptide N-termini and lysine residues. Dried samples were resuspended in a mixture of DIPEA and PSMtag in DMSO. The final sample solution contained peptides and PSMtag with a mass ratio of 1:3 in DMSO with 2% v/v DIPEA. The solution was mixed via pipetting to ensure sample is completely dissolved, spun down, and incubated at room temperature (RT) for 1-2 hours.

#### Simultaneous quench and removal of excess reagent

The size and hydrophobicity of PSMtag render C18-based clean ups of the quenched or NHS-ester species ineffective, thus we leveraged the NHS-ester chemistry to both quench and remove unreacted tag simultaneously. Amine-functionalized paramagnetic beads (MagReSyn, MR-AMN005) were washed once with water, then twice with DMSO prior to resuspension in DMSO to two times their original concentration (i.e. 40 mg/mL). The beads were then added at a 20:1 bead:PSMtag mass ratio to the peptide-tag mixture and allowed to incubate for 15-30 min at room temperature. Peptides were recovered by adding 4x the reaction volume of 500 mM TEAB, centrifuging briefly with a bench-top centrifuge, and placing samples on a magnetic rack to extract the peptide solution. The entire sample was then dried by vacuum centrifugation and either stored at −20°C or reconstituted in an aqueous solution of 30% acetonitrile, 0.05% DDM, 2% formic acid and analyzed using LC-MS/MS. A concentration of 0.015% DDM is typical for label-free single-cell LC-MS/MS. We found that at least 0.03% yielded improved solubility of labeled peptides and 0.05% is used routinely without notable impact on LC-MS/MS performance^63^.

### Evaluation of PSMtag for *de novo* sequencing

Training data was collected using a Vanquish Neo LC coupled to the Orbitrap Astral (Thermo Scientific) using gas-phase fractionation for greater depth using a data-dependent acquisition method. Six injections across an *m/z* range from 500-1000 as follows: 500-592, 588-650, 646-702, 698-762, 758-832, and 828-1000. LC conditions remained consistent for all injections: the gradient started at 30% MPB and increased to 38% in 0.1 min then ramped to 62% over 49 min for the active gradient. MPB was then increased to 75% MPB over another 10 min before washing and reequilibration. FAIMS was used for all analyses set to −50 CV. Resolution was set to 240K for MS1 with AGC of 300% and max injection time of 10 ms. The *m/z* range was limited to the range of the gas-phase fraction *±*20 Th. The Astral analyzer was used for MS2 limited to the *m/z* range noted above. Dynamic exclusion was set to Auto, AGC was set to 200%, and max injection time set to 100 ms. Collision energy was set dependent on precursor charge with 2+ precursors activated with 24 NCE, 3+ with 26, and 4+ with 28. This was done with the intent of DDA fragmentation spectra to be more similar to DIA spectra that does not normalize collisional energy to each charge state independently.

#### Analysis of non-tryptic peptides

Proteins from K562 cells were digested either with ArgC or GluC and subsequently labeled with PSMtag or left label-free (LF) as a control. All samples were again analyzed using a Vanquish Neo LC coupled to an Orbitrap Astral equipped with FAIMS. The label-free samples were loaded using 6% MPB and the gradient started at 0.1 min at 10% MPB, increased to 35% in 24.5 min, then to 42% in another 5.4 min. PSMtag-labeled samples were loaded at 30% MPB with the gradient starting at 0.1 min with 38%, then increasing to 62% in 24.5 min and finishing at 75% for the remaining 5.4 min. In both cases, the flow rate was maintained at 100 nL/min. FAIMS was used for all analyses set to −50 CV. Resolution was set to 240K for MS1 with AGC of 300% and max injection time of 100 ms. The *m/z* range for LF samples was 380-900 and 480-1000 for labeled. The Astral analyzer was used for MS2 with an AGC of 300% and max injection time of 100 ms. The collision energy was set to 24 NCE for GluC peptides and 26 NCE for ArgC peptides. Optimal collision energy was determined for each samples individually as assessed by maximum ID rate.

### Single-cell level standard data acquisition with timsTOF SCP

Samples were analyzed on a Bruker timsTOF SCP mass spectrometer coupled to a Vanquish Neo (Thermo Scientific) nLC system. Labeled peptides were diluted with 30% ACN, 0.18% dodecyl beta-D-maltoside (DDM), 2% FA and 200 pg were injected to mimic a single-cell level input. The samples were separated on a 25 cm x 75 *µ*m IonOpticks Aurora Ultimate CSI C18 column over a 19 min gradient. MPA remained at 0.1%FA (in water) and MPB at 80% ACN, 0.1% FA. The flow rate was kept constant at 250 nL/min with a column temperature of 55°C. The active gradient started at 30% MPB and 300 nL/min flow. %B was increased to 38% over 0.1 min and 62% over 12.9 min. At this point, the flow rate was reduced to 250 nL/min. MPB was then increased to 75% over 2 min and maintained for 3.1 min while the flow rate was increased to 300 nL/min again. Finally, MPB was ramped up to 99% over 0.1 min and maintained for 0.8 min. For long term sample acquisition, 0.05% DDM is optimal.

Unlabeled peptides were reconstituted in 0.015%DDM, 2% FA. The label-free samples were separated with an identical IonOpticks C18 column over a 19 min gradient (15 min active gradient). The active gradient started at 6% MPB and 300 nL/min flow. %B was increased to 10% over 0.1 min and 35% over 12.9 min. At this point, the flow rate was reduced to 250 nL/min. MPB was then increased to 42% over 2 min and to 75% over 0.1 min. Here it maintained for 3.1 min while the flow rate was increased to 300 nL/min again. Finally, MPB was ramped up to 99% over 0.1 min and maintained for 0.8 min.

The timsTOF SCP was operated in dia-PASEF mode with the following settings: MS1 *m/z* range of 100 - 1700, mobility scan range of 0.64 - 1.42 V.s/cm2, ramp and accumulation times of 100 ms. Capillary voltage was 1500 V, dry gas flow 3 L/min, and dry temp was 200°C. The minimum *m/z* that passes the quadrupole (low mass) was set to 380 and high sensitivity mode was enabled. The dia-PASEF windows covered an *m/z* range from 350 - 1050 and mobility range from 0.64 - 1.22 V.s/cm2. The collision energy was set to 24 - 64 eV as a function of the inverse mobility of the precursor. Each cycle consisted of 1 MS1 scan, 4 dia-PASEF cycles, 1 MS1 scan, 4 more dia-PASEF cycles for a total cycle time of 1.07 s. The dia-PASEF windows were 25 Th wide.

### Data acquisition with Orbitrap Astral (30-minute acquisitions)

Samples were separated via online nLC with a Thermo Vanquish Neo configured for trap-and-elute. Eluted peptides were introduced to an Orbitrap Astral mass spectrometer (Thermo Scientific) fitted with a FAIMS Duo Pro interface and an Easy Spray electrospray ionization (ESI) source. For all experiments, FAIMS was operated at −50 CV with a carrier gas flow set to 3.6 L/min, a spray voltage of 1900 V, and an ion transfer tube temperature 275°C. Samples (prepared identically as described above) were loaded onto the 5 mm x 0.3 mm trapping column (IonOpticks NS1-50100C18-3NZ) and analyzed with a 25 cm x 75 *µ*m IonOpticks Aurora Series UHPLC column (AUR3-25075C18-XT).

For all label-free samples, both DDA and DIA, initial conditions were 6% MPB at 300 nL/min. Flow rate was reduced to 100 nL/min, then ramped to 10% B over 0.1 min. The active gradient began at 10% MPB and increased to 35% over 24.5 min, followed by ramping to 42%B over 5.5 min. The column was then washed with 99% MPB for 4 min, whereupon the flow rate was returned to 300 nL/min prior to column re-equilibration. For the PSMtag samples, the same LC method structure was used as for the label free samples with the following exceptions. For PSMtag samples, initial conditions were 30% MPB at 300 nL/min and the active gradient occurred from 38% to 62% MPB with a 100 nL/min flow rate over 24.5 min, followed by an increase to 75%B over 5.5 min.

#### Data-dependent acquisition

For DDA experiments, 2 ng of human proteome were injected for analysis. MS1 scans were collected at a resolution of 240,000 over an *m/z* range of 380-900 with a maximum fill time of 100 ms, and an AGC of 3e6. The RF lens was set to 50%. Following the MS1 scan, ions with charge states 2-5 and *m/z* 400-900 for label-free or *m/z* 500-1000 for labeled peptides were selected based on a cycle time of 1 second for MS2 fragmentation and measurement in the Astral detector. Dynamic exclusion was implemented, set to the “auto” setting and the MS2 fragment scan range was set to 120-1800 *m/z*. Ions were isolated with a 1 *m/z* window, 0.2 *m/z* offset. AGC was set to 2e4 and maximum injection time of 50 ms. Peptides were fragmented with 24, 26, 28, or 30 NCE for experiments determining optimal collision energy for each sample type.

#### Data-independent acquisition of single-cell inputs

For mixed-proteome DIA injections, 200 pg of human proteome were analyzed with variable yeast proteome spike-ins. MS1 scans were collected at a resolution of 240,000 over an *m/z* range of either 380-900 (label-free) or 480-1000 (labeled) with a maximum fill time of 150 ms and an AGC of 300%. The RF lens was set to 50%. DIA spectra were collected with the Astral detector using 20 windows of variable size as listed in Table 1. The fragment scan range was set to 150-2000 with an acquisition loop set to 0.6 sec. Collision energy was set to 28% (label-free) or 24% NCE (labeled) with an AGC set to 500% and maximum injection time of 60 ms. LC parameters were identical to DDA injections.

**Table 1.** 20 Window DIA Method.

| Label-free | PSMtagged |
|---|---|
| 400-438.5 | 500-538.5 |
| 438-466.5 | 538-566.5 |
| 466-490.5 | 566-590.5 |
| 490-511.5 | 590-611.5 |
| 511-533.5 | 611-633.5 |
| 533-549.5 | 633-649.5 |
| 549-563.5 | 649-663.5 |
| 563-574.5 | 663-674.5 |
| 574-585.5 | 674-685.5 |
| 585-600.5 | 685-700.5 |
| 600-616.5 | 700-716.5 |
| 616-628.5 | 716-728.5 |
| 628-643.5 | 728-743.5 |
| 643-654.5 | 743-754.5 |
| 654-671.5 | 754-771.5 |
| 671-689.5 | 771-789.5 |
| 689-716.5 | 789-816.5 |
| 716-747.5 | 816-847.5 |
| 747-793.5 | 847-893.5 |
| 793-900 | 893-1000 |

#### Data-independent acquisition of mixed human proteomes

For analysis of diverse human cell lines, 2 ng each were injected either label-free or labeled with 9-plex PSMtags. The gradients were identical as to what is described immediately above, but carried out with an active gradient flow rate of 250 nL/min. The MS1 resolution was maintained at 240,000 with a max injection time of 20 ms. The AGC for LF was 300%, but was increased for 5-plex and 9-plex injections with the intent to scale the ion population with the plex size. The maximum setting, however, is 1000%, which was used for both 5- and 9-plex samples. The MS1 *m/z* range was 350-900 for label-free samples with DIA windows spanning 400-900 and 450-1050 for PSMtag samples with DIA windows spanning 500-1000. MS2 AGC was scaled with the number of samples analyzed per run with 300% for LF, 1500% for 5-plex, and 3000% for 9-plex. Optimal window schemes were determined for each sample independently: LF samples were acquired with 5-Th fixed window widths with 8-ms injection times; 5-plex samples were acquired with 2-Th fixed window widths and 3-ms injection times; and 9-plex samples were acquired with 2-Th fixed window widths and 10-ms injection times. Further, 28 NCE was used for LF analyses and 24 NCE for PSMtagged. FAIMS was not used for this analysis.

### Data acquisition with Orbitrap Astral (11-minute acquisitions)

Samples were separated via a dual-column online nLC as described by Wang *et. al*^25^, with the exception of the analytical column, here using an analytical column with dimensions 75*µ*m ID x 20cm packed with 1.6*µ*m 100 Å C18 (Micromics). The 11-minute acquisition is manually-positioned on different portions of the overall gradient for label-free or PSMtag-labeled samples to optimize total ion current for these different sample types. MS parameters remain as described for the 30-minute acquisition, with the exception that the ion transfer tube temperature was 200 °C, The spray voltage was 2400 V and MS1 scan range was 380-1100 m/z (label-free) or 480-1200 m/z (PSMtag).

### Peptide-spectra-matching parameters

#### MaxQuant 1.6/2.7

Raw data from the candidate mass tag screen were processed with MaxQuant (version 1.6.0.16 or 2.7.0.0) using the integrated Andromeda search engine^32,35,64,65^. MS/MS spectra were searched against the 2 model peptide sequences, supplemented with the MaxQuant common contaminants database and reversed-sequence decoys. Enzyme specificity was set to Trypsin/P, allowing up to two missed cleavages and a minimum peptide length of seven residues. Carbamidomethylation of cysteine was a fixed modification. Methionine oxidation and protein N-terminal acetylation were variable modifications, with a maximum of five modifications per peptide. Precursor mass tolerances were 20 ppm for the first search and 4.5 ppm for the main search after nonlinear mass recalibration, and fragment ion tolerance was 20 ppm. False discovery rates (FDR) at the peptide-spectrum match, protein, and site levels were controlled at 1% using the target-decoy approach.

To discover unanticipated modifications, the dependent peptide (DP) search was enabled^62^. In this mode, MS/MS spectra not assigned in the main search were compared with spectra of identified, unmodified base peptides. The precursor mass difference was treated as an unknown modification. DP results were extracted from the allPeptides.txt output table and matched to possible mass tag candidate masses according to experimental design (isobaric candidates were acquired in separate LC-MS acquisitions) and at mass accuracy of *±*10 ppm.

#### Sage 0.14.6

Label-free and PSMtag DDA data were searched with Sage version 0.14.6. A Trypsin/P digest was configured with up to 2 missed cleavages, a peptide length range of [3, 50] AA, a peptide mass range of [300, 6000] Da, and a fragment m/z range of [50, 2000]. Precursor charges [2, 4], fragment charges [1, 3], and precursor isotope errors [*−*1, 3] were considered. A precursor mass tolerance of *±*20 ppm and fragment mass tolerance of *±*10 ppm were used. Cysteine carbamidomethylation and PSMtag were set as static modifications, with methionine oxidation as a variable modification. Sage LFQ was enabled with the following parameters: *{peak scoring:“Hybrid”, integration:“Sum”, spectral angle:0.7, ppm tolerance:15.0, combine charge states:false}*. Spectra were not deisotoped.

#### PointNovo

The pytorch implementation of PointNovo (github.com/irleader/PointNovo) was used for model training and sequence prediction. The default configuration in config.py was edited to set *PRECURSOR MASS PRECISION TOLERANCE=0.05*; *spectrum reso=50*; and to add PSMtag as a static modification mass on N-term and K.

#### Novor

Novor search parameters were specified as follows: Precursor error tolerance: 25 ppm; Fragment error tolerance: 0.02 Da; PSMtag as static PTM on K, n-term; Variable N deamidation; Variable M oxidation; Variable C carbamidomethylation; MS2 mass analyzer: Orbitrap; Fragmentation: HCD; FDR: 1

#### DIA-NN 2.5

DIA-NN^26^ version 2.5.1 was used to search the PSMtag data using library fine tuning^47^. PSM-tag was searched as a fixed modification using *{–fixed-mod tag, 308.1160923903, nK –channels tag,d0,nK,0:0; tag,d4,nK,4.01096:4.01096;tag,d8,nK,8.026839:8.026839; tag,d12,nK,12.033938:12.033938; tag,d16,nK,16.038574:16.038574 –original-mods}*. Quantitation was set to *–high-acc*. The normalization strategy was set to *–channel-spec-norm*.

#### JMod 2.0

JMod^28^ version 2.0.0 was used to search the PSMtag data. The same library as DIA-NN was used for all searches. The PSMtag plex size (*PSMtag, PSMtag 5plex, PSMtag 9plex*) was selected for each respective sample and all were searched using *{--apex jitter 3 --free apex --additional scans 3 --no ms1 req --use emp rt}* with other parameters set to default.

## Code availability

Data analysis was performed using the programming language R and packages: pak, devtools, reshape2, ggridges, tidyverse, ggpubr, ggpattern, arrow, gridExtra, grid, eulerr, ggh4x, xml2, googledrive, readxl, dplyr, patchwork, stringr, ggsignif, proteomicsCV, and diann. Exact package versions are specified in the GitHub repository provided. Data analysis was also performed using the programming language python and packages: pandas, numpy, matplotlib, matplotlibvenn, seaborn, nltk. Exact package versions are specified in the GitHub repository provided. All code for reproducing figures available at: github.com/ParallelSquared/PSMtag

## Data availability

Data, raw and processed, for reproducing figures is available through data repositories MassIVE MSV000097968, MSV000103730 and ProteomeXchange PXD064191, PXD086351.

Processed data organized by contributing figure is available at data repository Zenodo under DOI https://doi.org/10.5281/zenodo.23675860.

## Extended Data Figures

**Extended Data Fig. 1.**
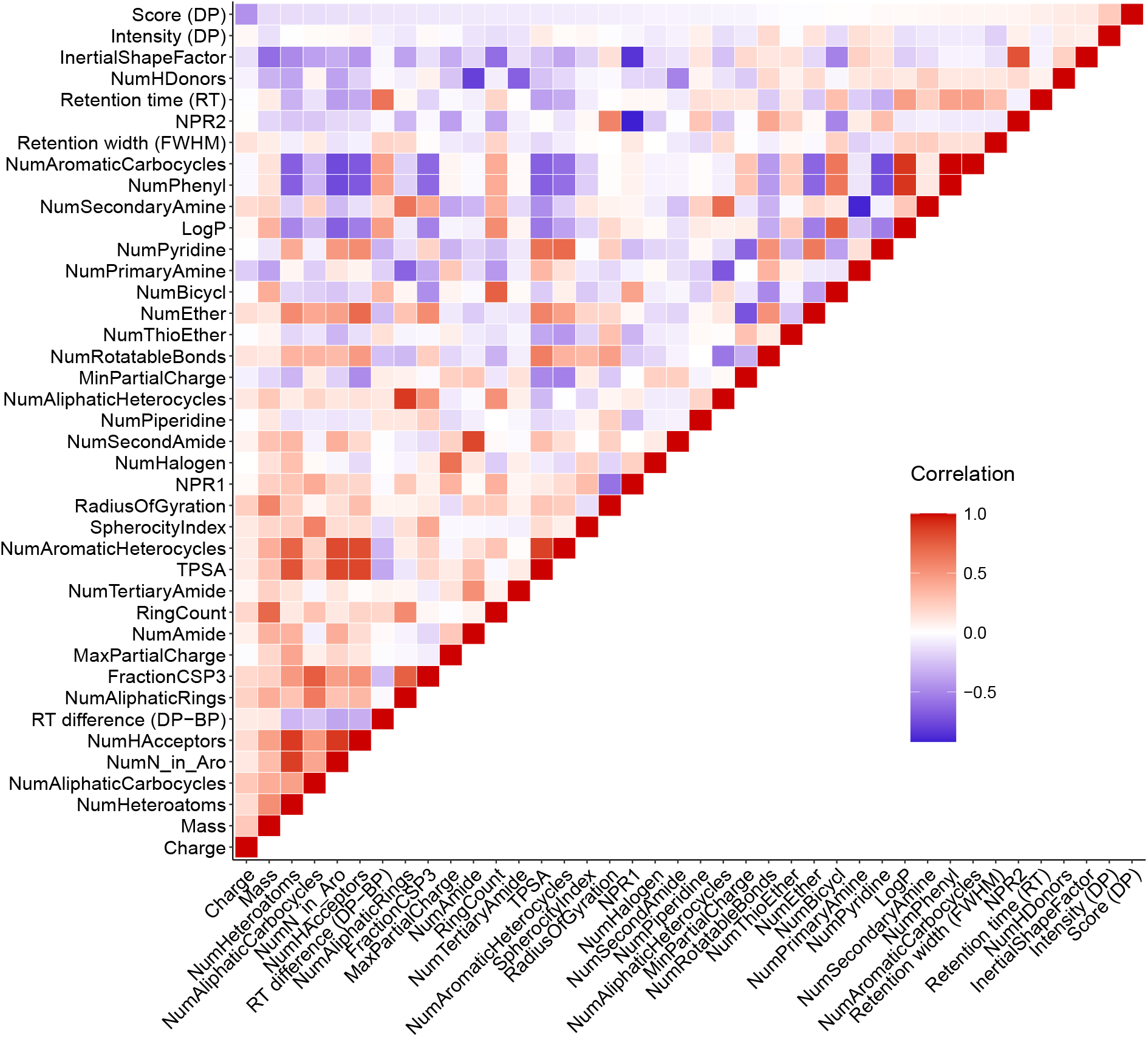
A more comprehensive comparison of chemical and LC-MS features of the tag screen. For tags observed on at least one model peptide, their chemical properties are estimated from their unconjugated SMILES string by RDKit^33^ and their LC-MS features are calculated by MaxQuant v2.7^32^. The features are ordered by Spearman correlation to Dependent Peptide Score (Score (DP)), the primary metric to assess quality of spectra produced by tag-peptide conjugates. Redundant information in the lower triangle is omitted.

**Extended Data Fig. 2.**
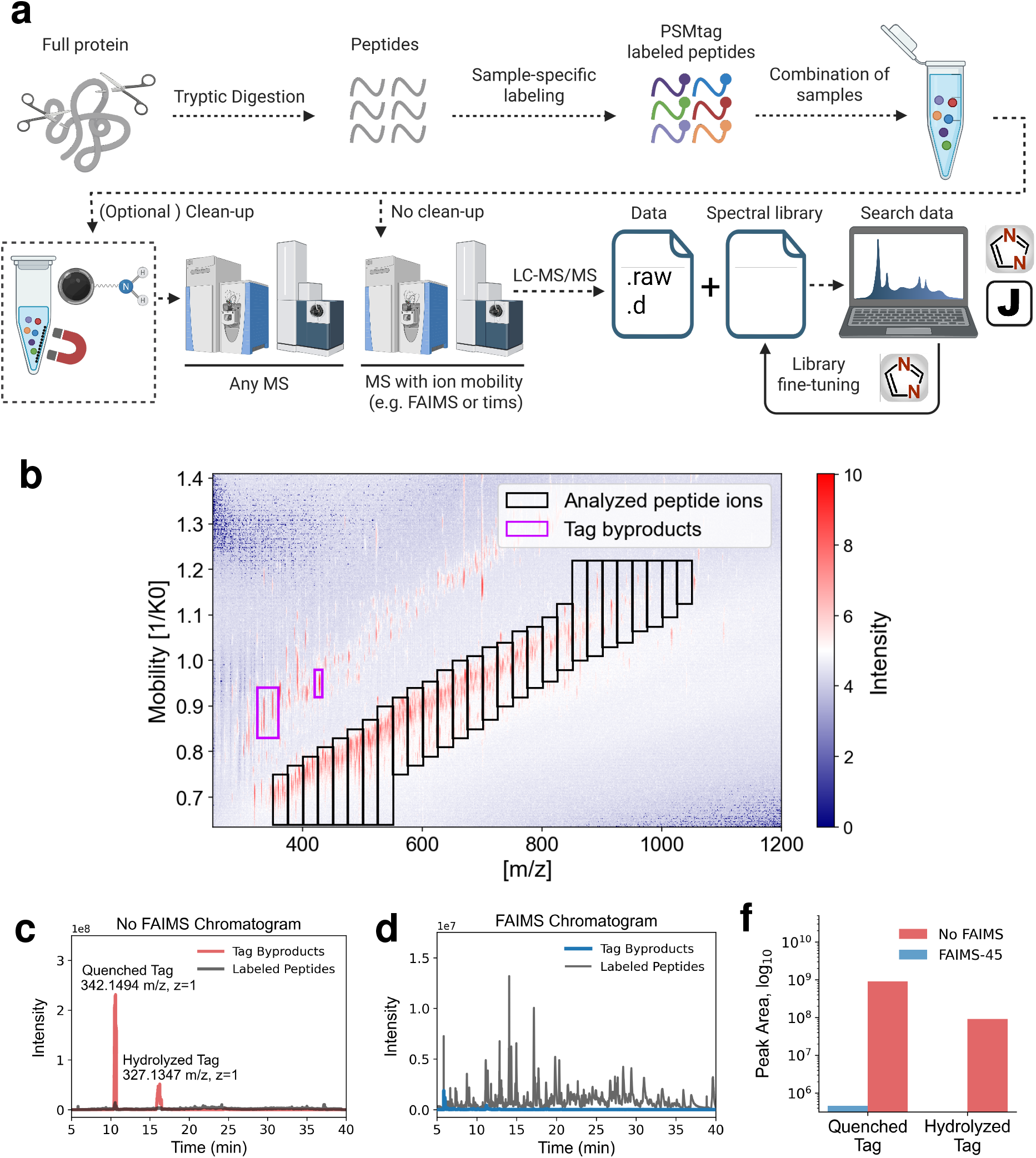
Full PSMtag workflows that remove tag by-products by magnetic bead clean-up or use ion mobility properties to separate co-eluting tag byproducts and tagged peptides. **a**, The current state of the PSMtag workflow for sample multiplexing and analysis. Created in BioRender. Specht, H. (2026) https://BioRender.com/lyrgcdy. **b**, The excess tag remaining after conjugation to peptides (as a quenched or hydrolyzed product, shown in pink box) is separable from peptides by ion mobility as implemented by trapped ion mobility (TIMS) (shown) or field asymmetric ion mobility spectrometry **c**, The relative extracted total ion currents for quenched (m/z 342.15) and hydrolyzed (m/z 327.13) tag byproducts (red) and tagged peptides (grey) without using FAIMS Pro. **d**, The relative extracted total ion currents for tag byproducts (red) and tagged peptides (grey) using FAIMS Pro at −45 V. **e**, The total extracted total ion currents for tag byproducts without using FAIMS (red) and using FAIMS (blue).

**Extended Data Fig. 3.**
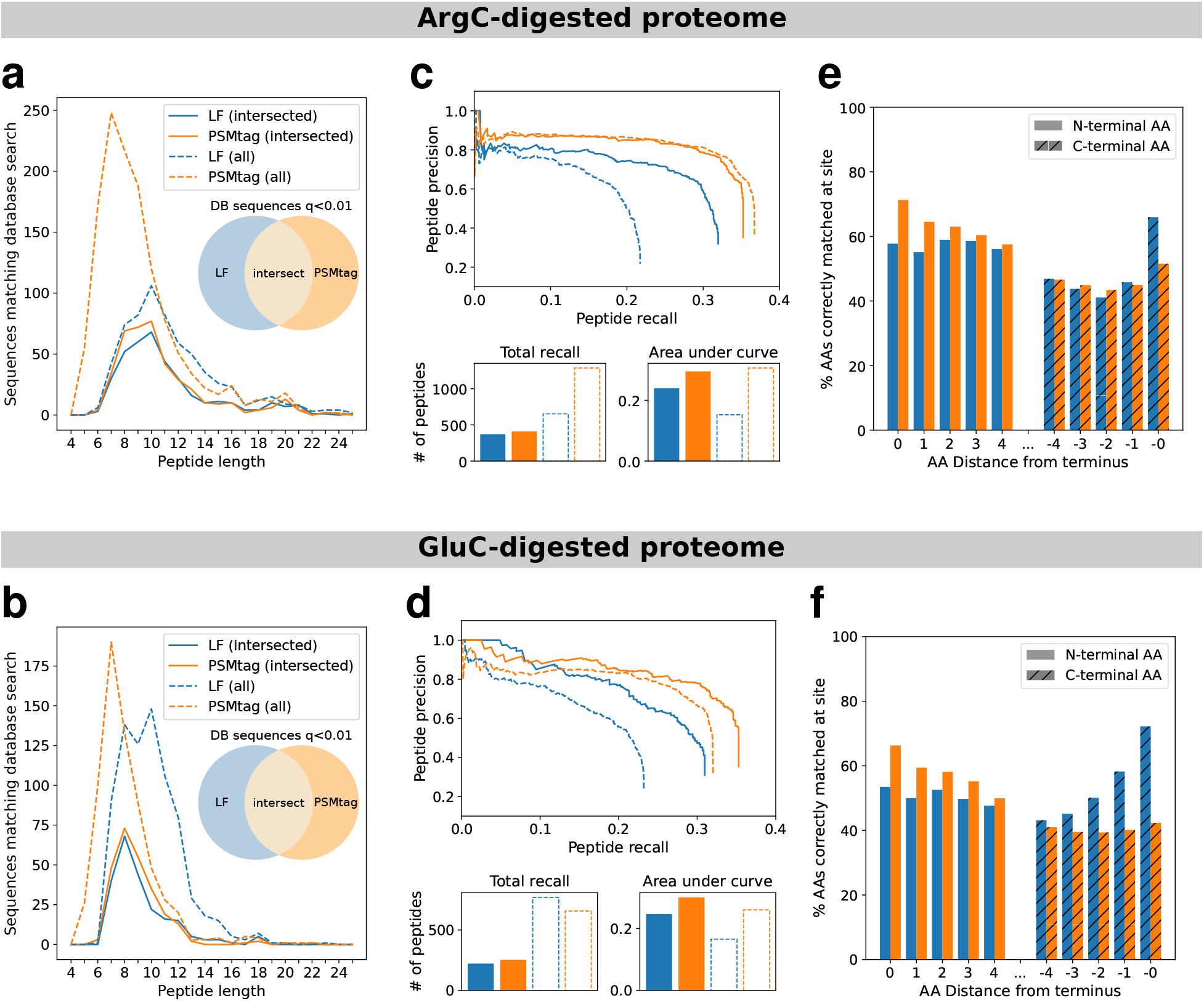
Improvements to Novor *de novo* sequencing with ArgC- and GluC-derived peptides. **a, b.** Total numbers of correct *de novo* peptide identifications, conditioned on peptide length. Peptide spectra assigned the same sequence by both Novor and Sage (at Sage spectrum q-value *<*0.01) are considered correct *de novo* matches. **c, d.** Peptide-level precision-recall curves for tagged and label-free samples, with measurements of total numbers of peptides recalled and area under each precision-recall curve. **e, f.** Overall accuracy of *de novo* amino acid assignments for residues with a distance of up to 4 AA from the peptide N- or C-terminus. Amino acids in the *de novo* sequence are considered to be correctly assigned if the database sequence contains a counterpart amino acid within 0.02Da in mass, with a prefix mass (sum of all preceding amino acids) within 0.1Da.

**Extended Data Fig. 4.**
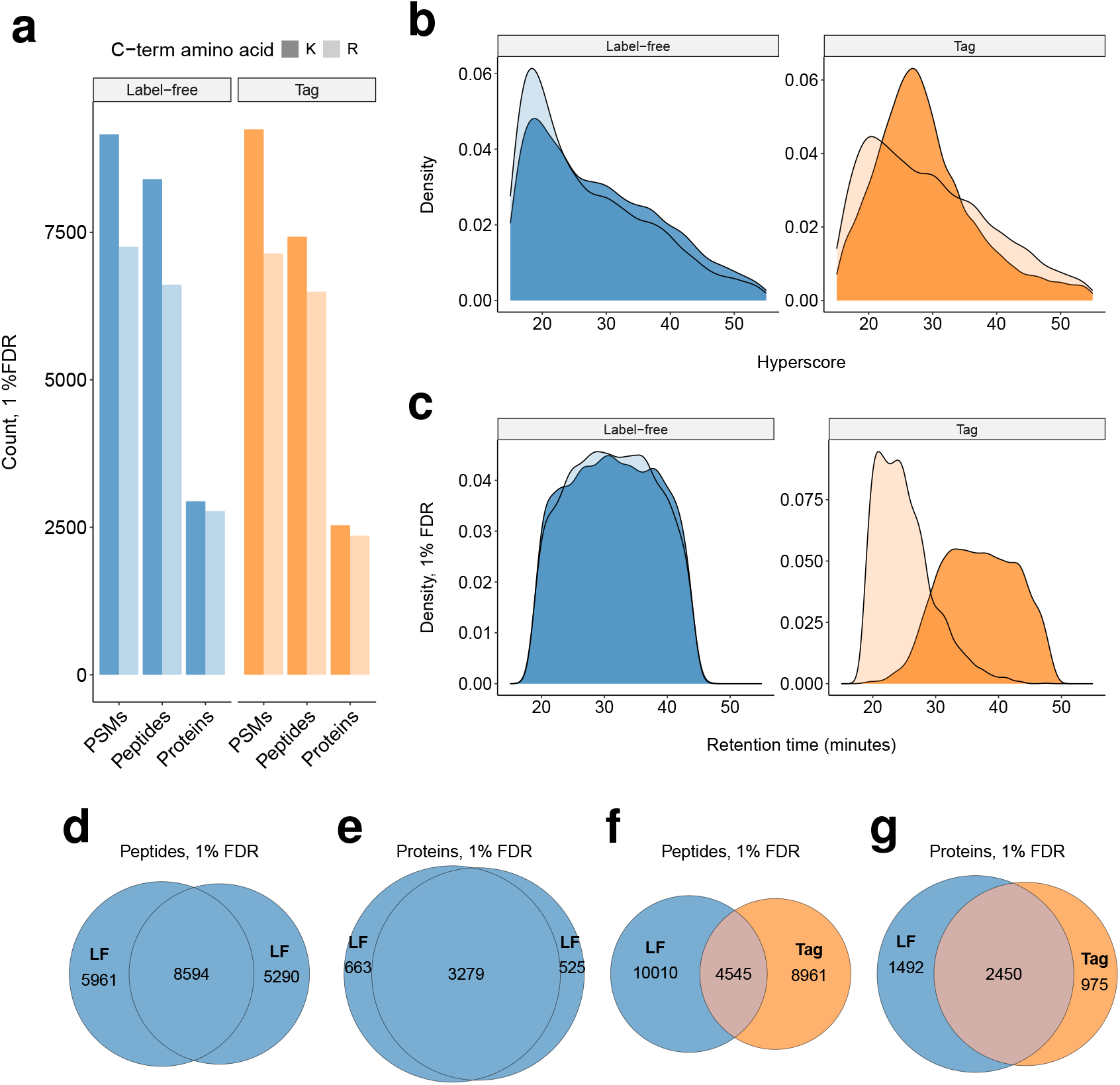
Descriptive statistics on peptides and proteins identified from 2ng tagged and label-free samples by DDA. All data was acquired on the Thermofisher Astral platform. **a**, The counts at 1% FDR (spectrum Q-value calculated by Sage): peptide-spectra-matches (PSMs), peptides identified with or without quantitative information extracted by Sage, peptides identified with quantitative information extracted by Sage, and likewise for proteins. Counts supported by a sequence with at least one arginine are lighter shades. Counts supported by a sequence with at least one lysine are darker shades. **b**, The density distributions of peptides’ scores. Peptides with arginine on the C-terminus are the lighter shades. Peptides with lysine on the C-terminus are darker shades. **c**, The density distributions of peptide retention times. While most arginine and lysine terminating peptides elute together when label-free, the lysine peptide elute significantly later due to carrying an extra tag. **d**, Peptides observed in common or uniquely for two label-free replicates, one replicate at normalized collision energy (NCE) 28, the other at NCE = 24. **e**, Proteins observed in common or uniquely for two label-free replicates, one replicate at normalized collision energy (NCE) 28, the other at NCE = 24. **f**, Peptides observed in common or uniquely for tagged (NCE=24) and label-free samples (NCE=28). **g**, Proteins observed in common or uniquely for tagged (NCE=24) and label-free samples (NCE=28).

**Extended Data Fig. 5.**
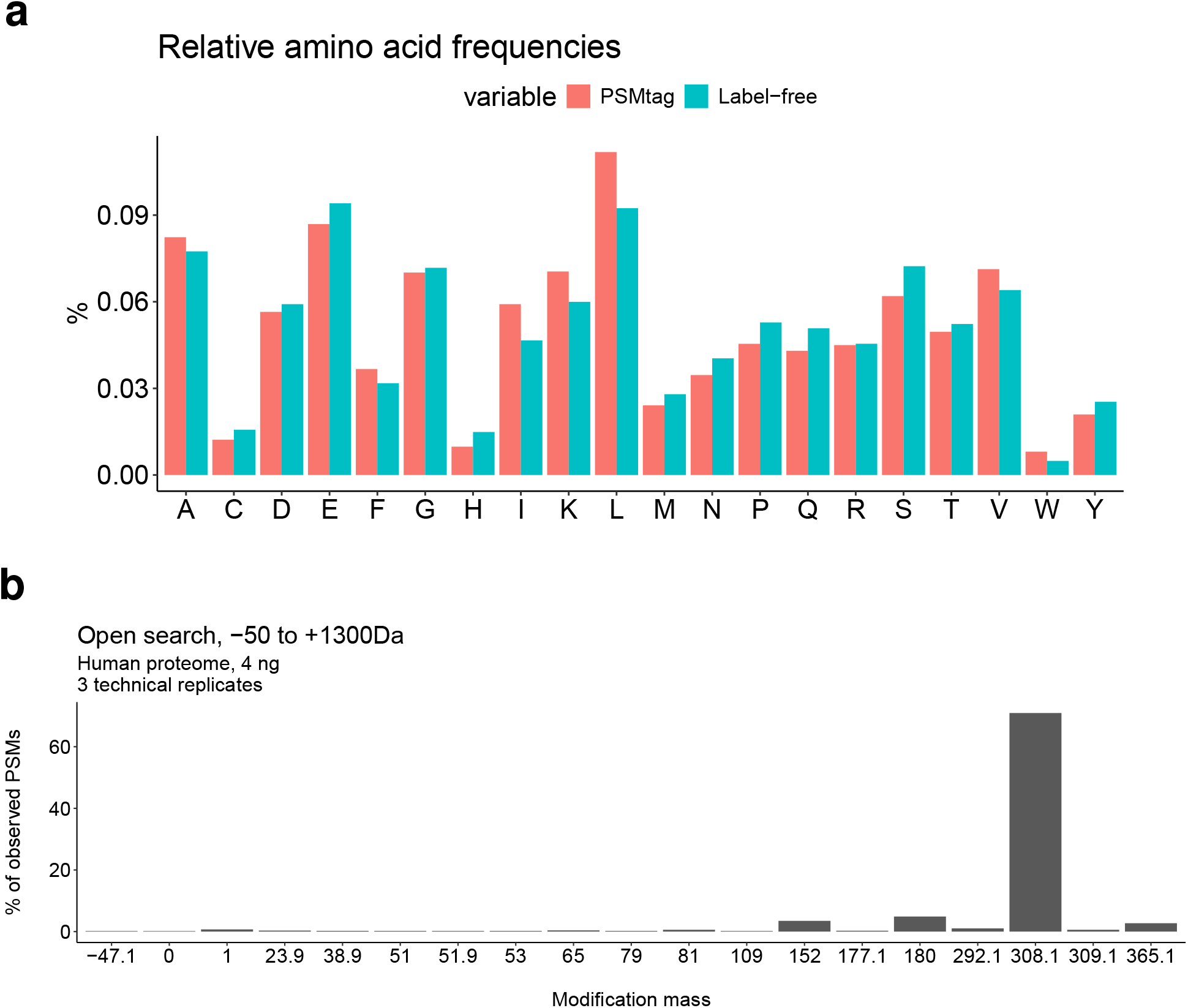
Other labeling products of PSMtag and the relative observation of amino acids in PSMtag vs. label-free proteomes. **a**, Three replicate injections of PSMtag labeled human proteomes (K-562 cell lines) of 4 ng were acquired by the Orbitrap Astral and analyzed by MSFragger using the open search workflow over a range of −50 Da to 1300 Da. **b**, The DDA data used in Extended Data Fig. 4 was examined for the relative observation frequency of amino acids and labeled and label-free peptides to understand the source of different peptides observations Extended Data Fig. 4f.

**Extended Data Fig. 6.**
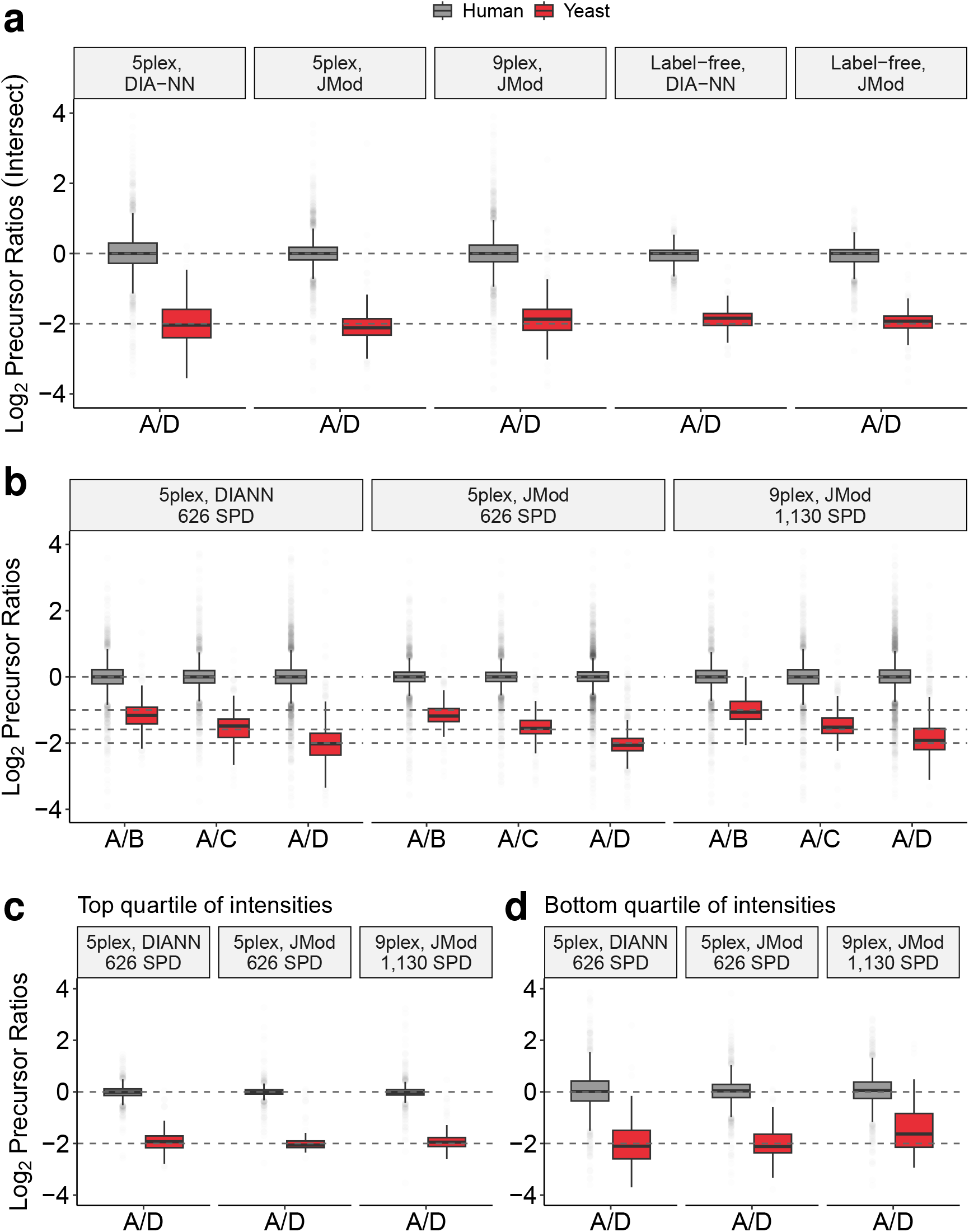
Extended examination of yeast spike in ratios. All data acquired using the 11-minute gradient LC-MS acquisition described in Methods. **a**, Examining the same mixing ratios from Fig. 4g, adding in a label-free benchmark using the same experimental design and visualizing only precursors in common between the sample types. **b**, Examining more mixing ratios as described in Fig. 4g. Measured ratios for the most abundant precursors (top quartile) in **c** and the least abundant precursors (bottom quartile shown) in **d**. Fourteen replicates per sample type were used for PSMtag only comparisons. Seven replicates per sample type were used for PSMtag and label-free comparisons to control for data availability of label-free. PSMtag replicates were randomly sampled down to seven.

**Extended Data Fig. 7.**
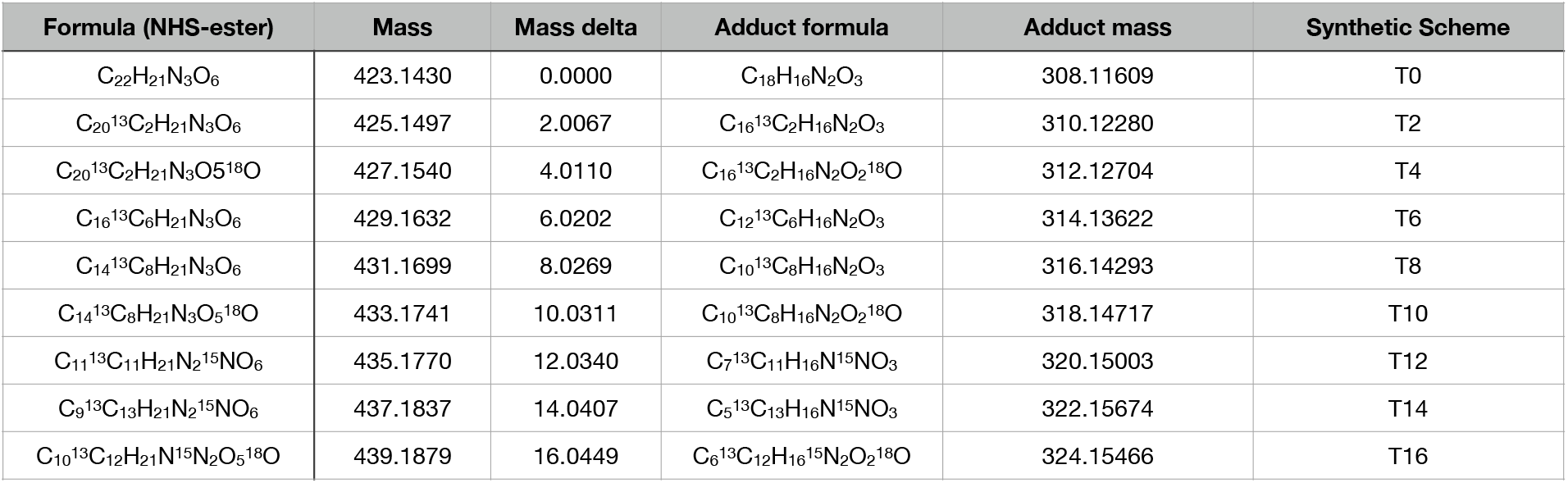
Empirical formulas and masses for tag reagents. The empirical formulas for the isotopically-doped reagents functionalized with NHS-ester allowing 5-plex and 9-plex with their corresponding monoisotopic masses (Mass) and differences in mass from the lowest mass isotopomer (Mass delta). The nomenclature for the reagents as used in the Synthetic Scheme is included (”T0”-”T16”).

**Extended Data Fig. 8.**
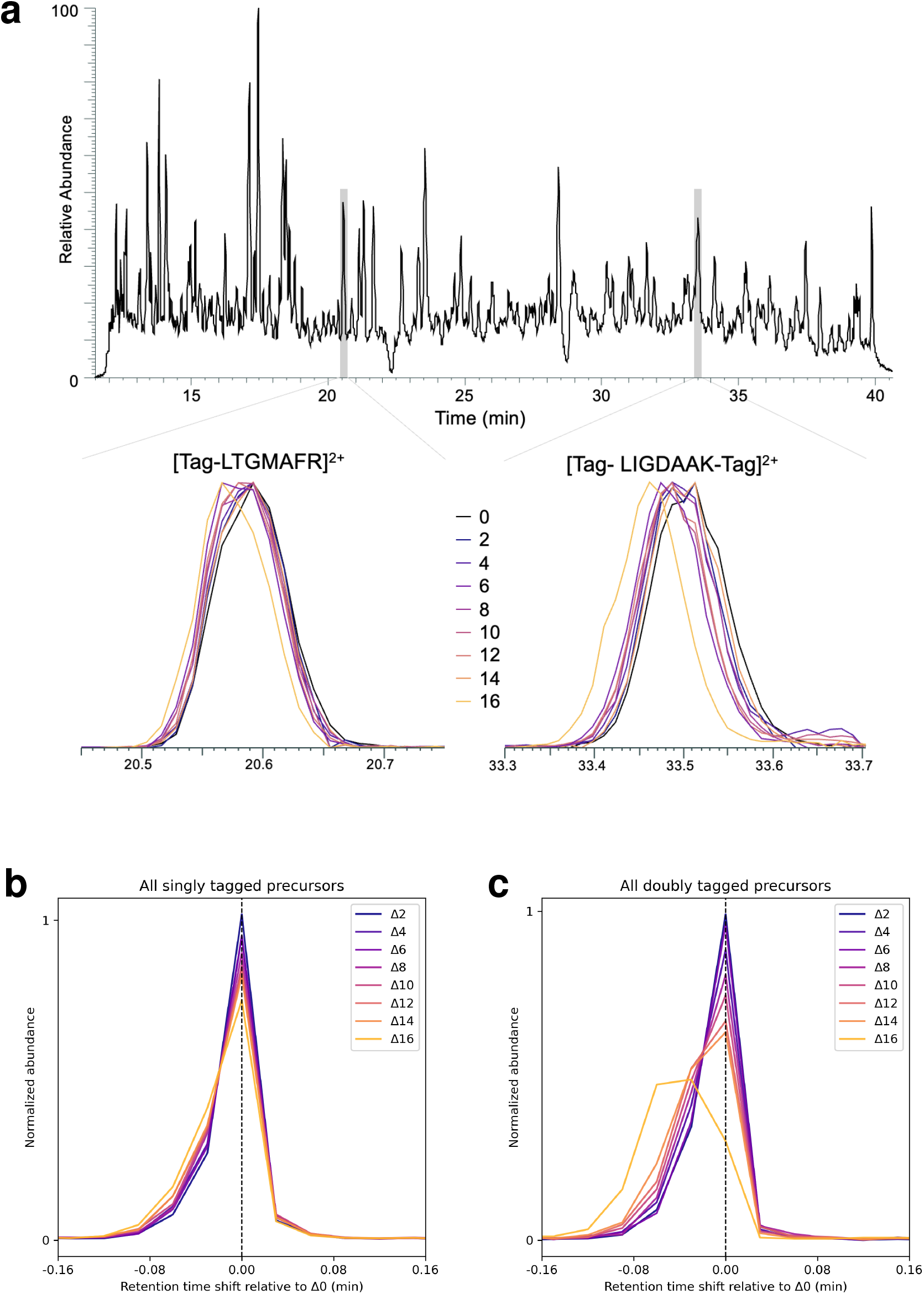
The effect of isotopic doping on tagged peptide retention time. **a**, The extracted signal for select high-abundance tagged peptides and their isotopologues. **b**, The differences in retention time for all confidently-identified singly tagged peptide isotopologues from their respective Δ0 channel. **c**, The differences in retention time for all confidently-identified doubly tagged peptide isotopologues from their respective Δ0 channel.

**Extended Data Fig. 9.**
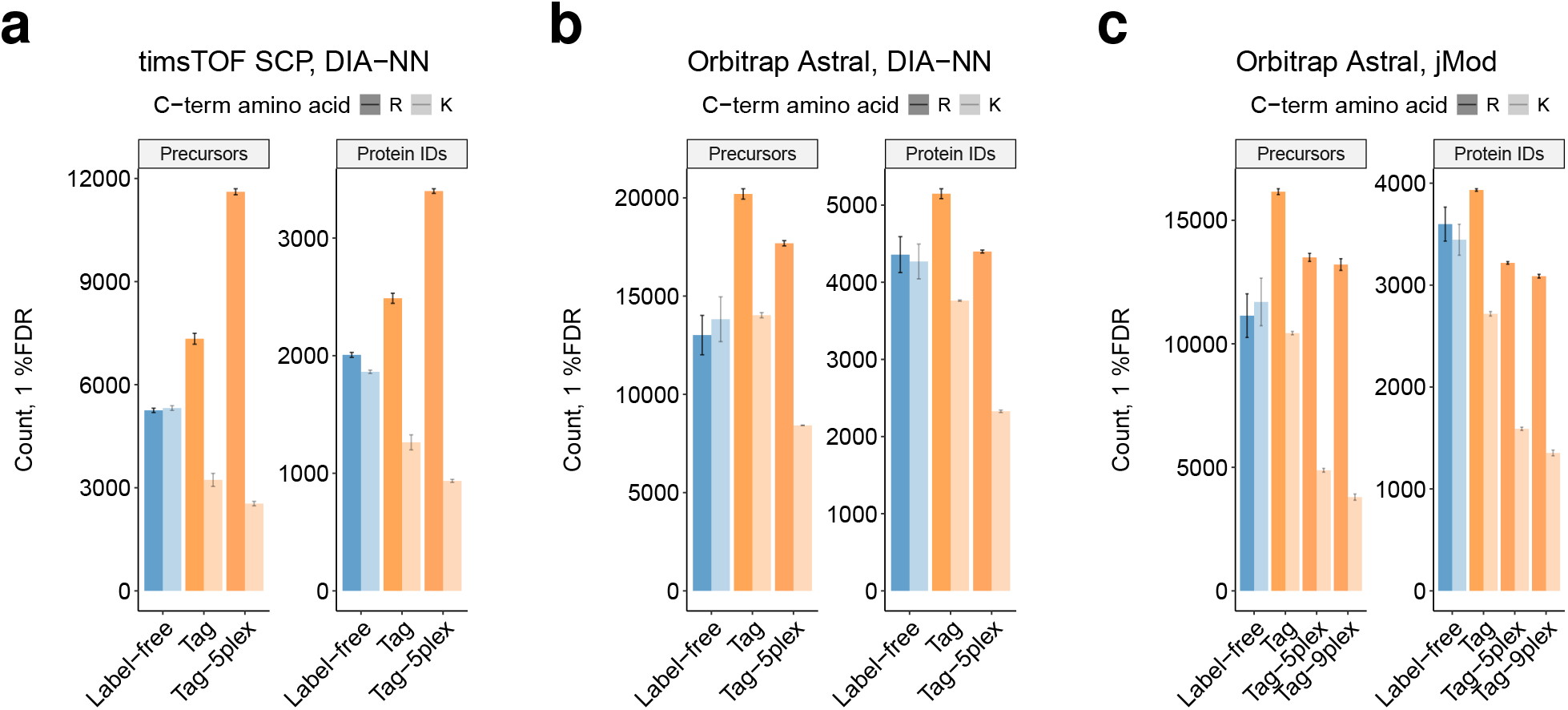
Identification rates for tagged and label-free single-cell-level (200pg) proteomes by DIA. **a**, The number precursors containing at least one lysine or arginine and proteins supported by these precursors identified in replicate injections of a 200pg human proteome labeled, unlabeled, and labeled with 5-plex reagents. Data acquired using the timsTOF SCP mass spectrometer and analyzed with search engine DIA-NN 2.5. **b**, The same, but data acquired with Orbitrap Astral mass spectrometer. **c**, The same, but data (Astral) analyzed with JMod.

**Extended Data Fig. 10.**
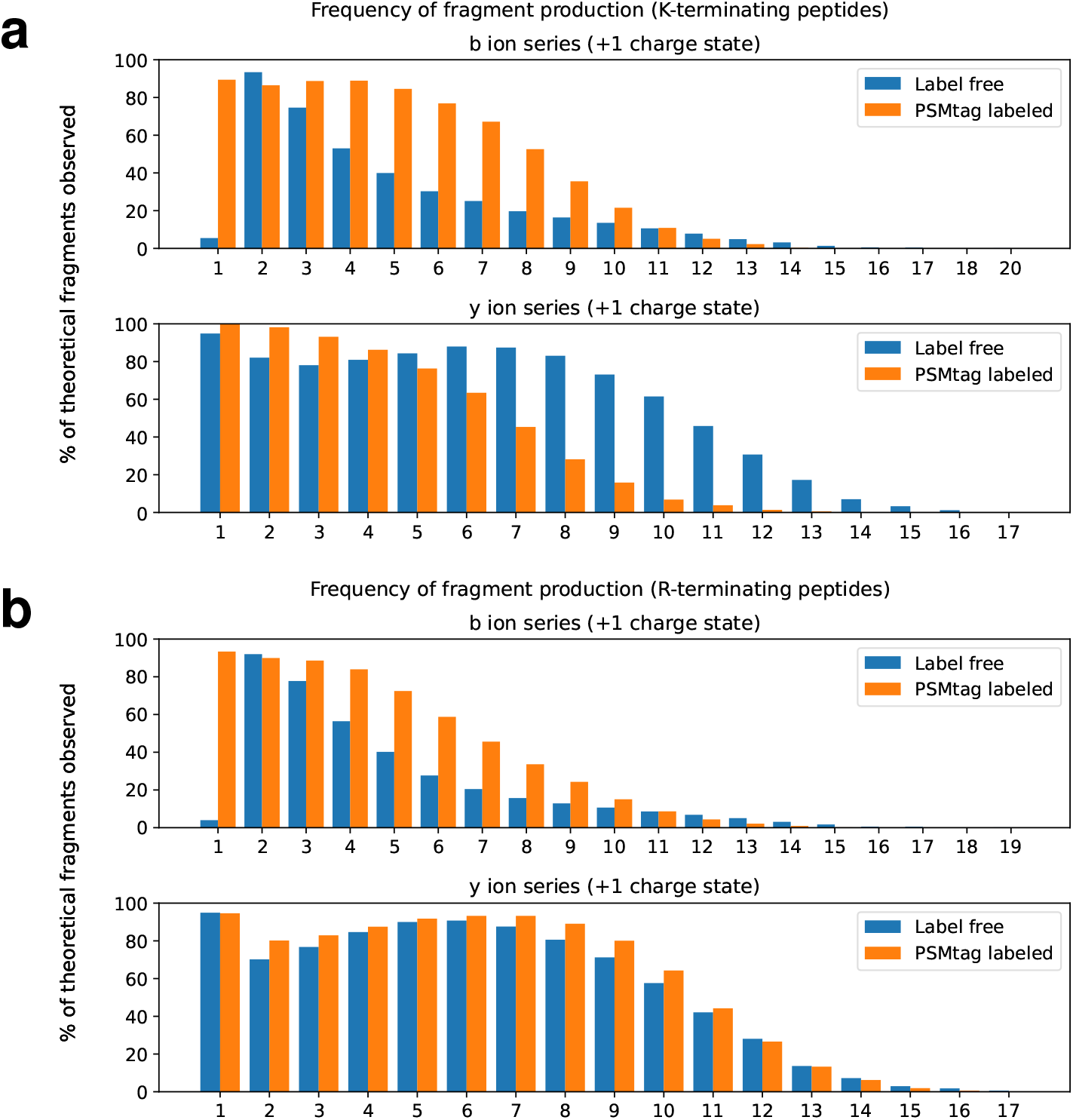
Fragment ion series with one positive charge for tagged and label-free peptides. **a**, The frequency of observing b-ions and y-ions with one positive charge for all peptides confidently assigned to terminate in lysine. **b**, The frequency of observing b-ions and y-ions with one positive charges for all peptides confidently assigned to terminate in arginine. Data was acquired on the Orbitrap Exploris 240 platform using collision energies and FAIMS CV values optimized to maximize peptide identifications from the different sample types, label-free and labeled.

## Supplemental

**Supplemental Fig. 1.**
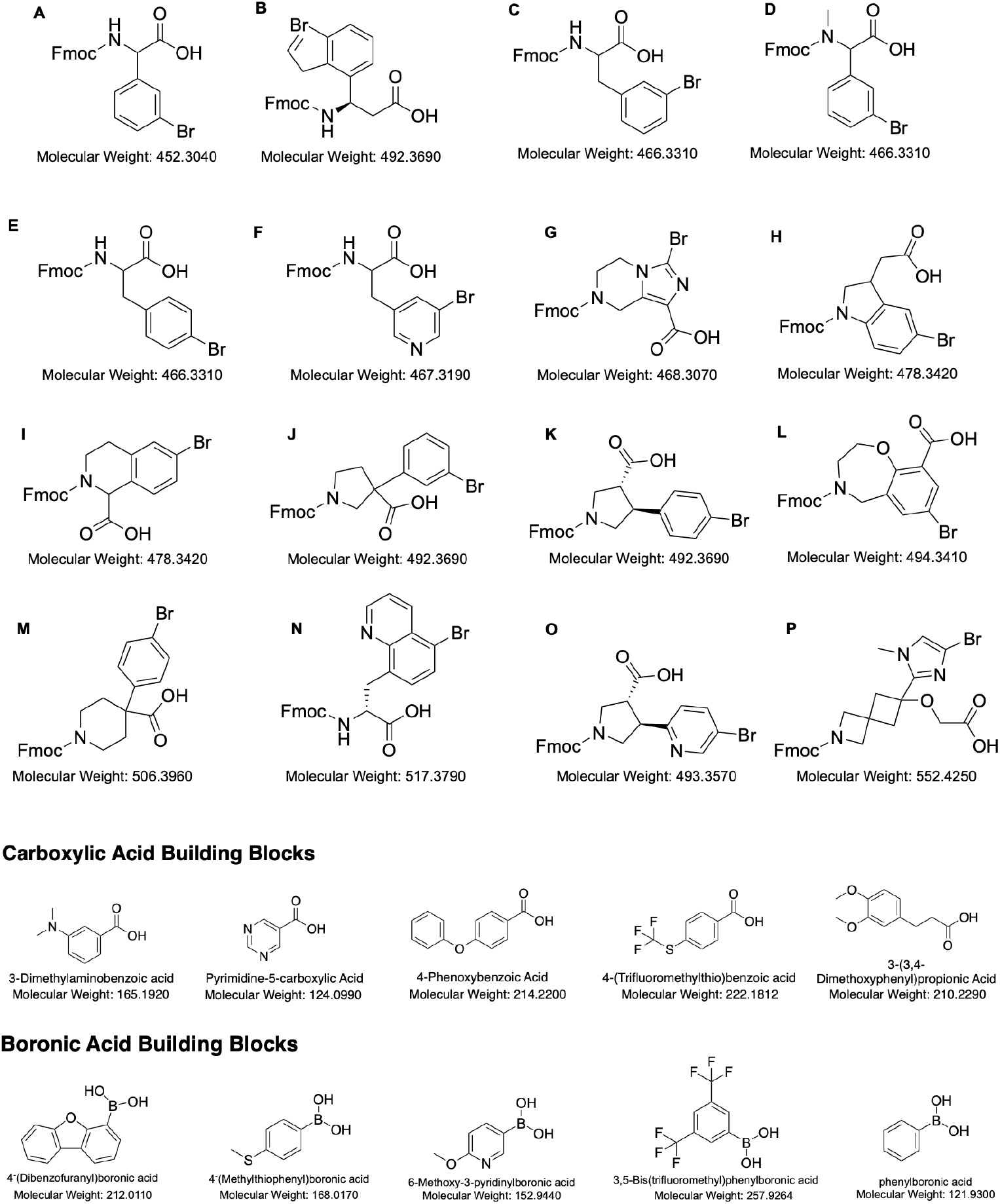
Building blocks for split-pool synthesis of mass tag candidates. The boronic acids, carboxylic acids and trifunctional cores used.

**Supplemental Fig. 2.**
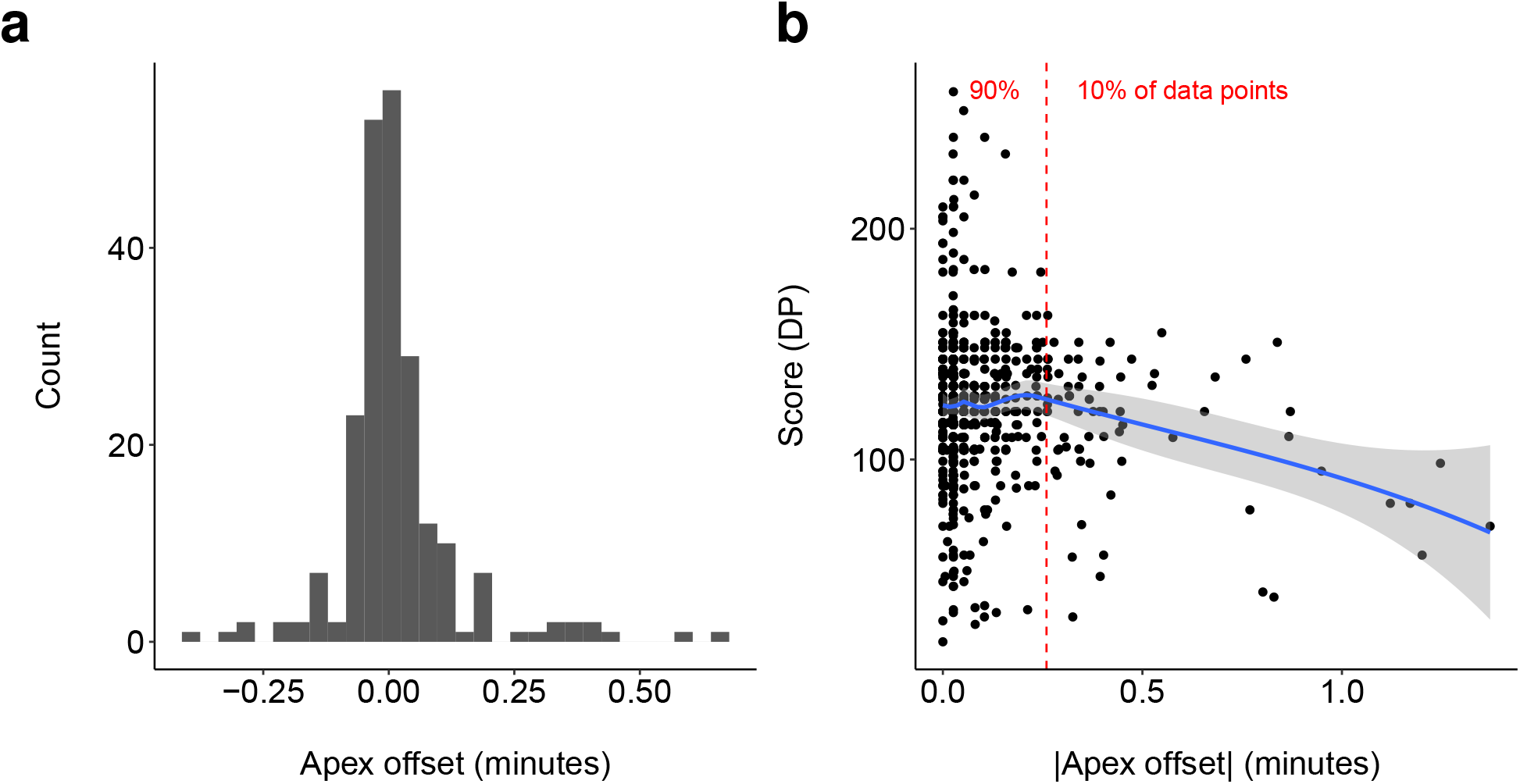
Apex offset for tag-peptide conjugates in combinatorial screen. **a**, The apex offset, the time from the recognized apex of the elution profile of each identified tag-peptide conjugate to the time it was sequenced by the MS detector. **b**, The absolute value of the apex offset. The dependent peptide Andromeda score is shown on the y-axis. Loess regression (blue) and a 95% confidence band (gray) estimate the trend.

**Supplemental Fig. 3.**
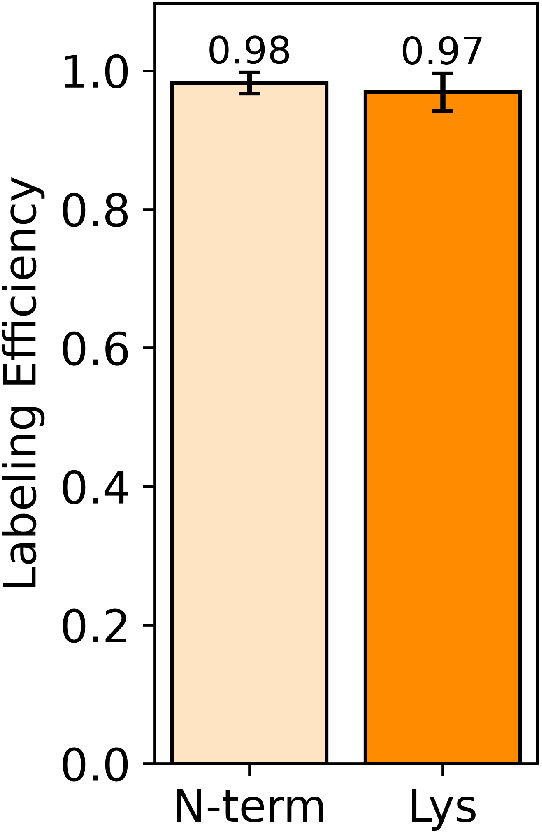
The labeling efficiency achieved labeling human proteomes with PSMtag. Average labeling efficiency calculated from triplicate, 2-ng injections of human proteome totaling 7,827 confident identifications.

**Supplemental Fig. 4.**
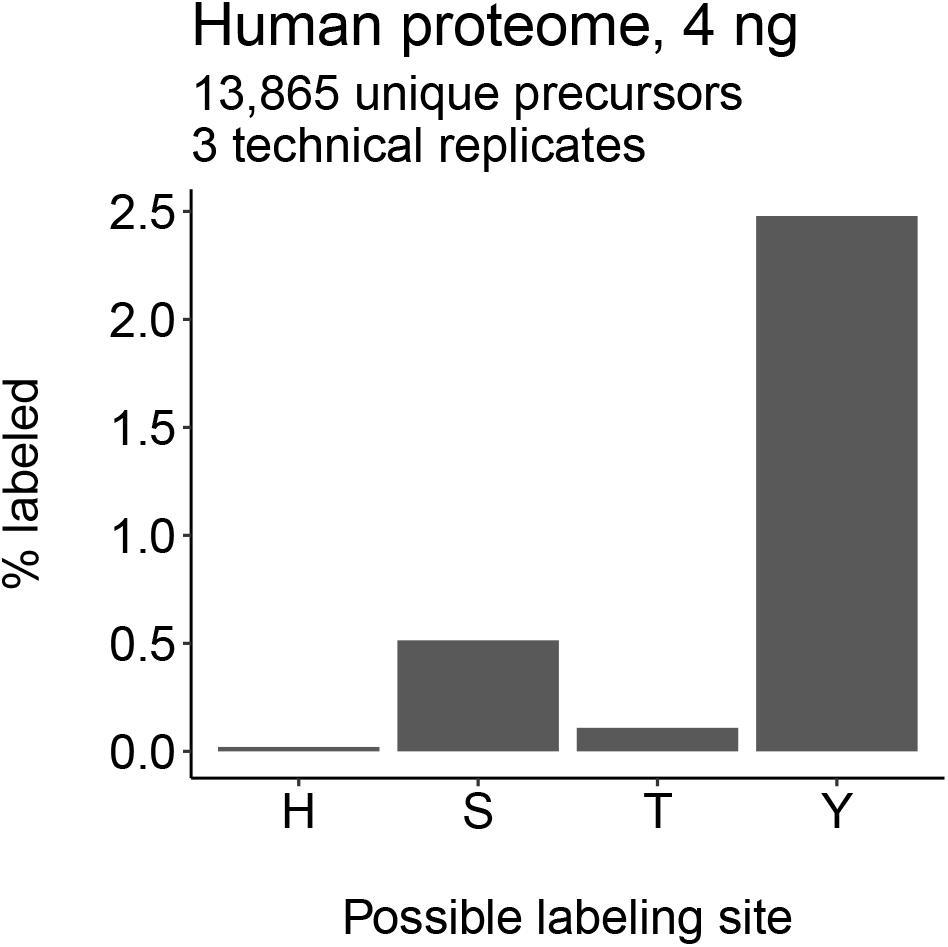
Off-target labeling by PSMtag. Three replicate injections of PSMtag labeled human proteomes (K-562 cell lines) of 4 ng were acquired by the Orbitrap Astral and analyzed by MaxQuant using fixed modification of PSMtag at lysine and N-terminus and variable modifications of PSMtag at histidine (H), serine (S), threonine (T), tyrosine (Y).

**Supplemental Fig. 5.**
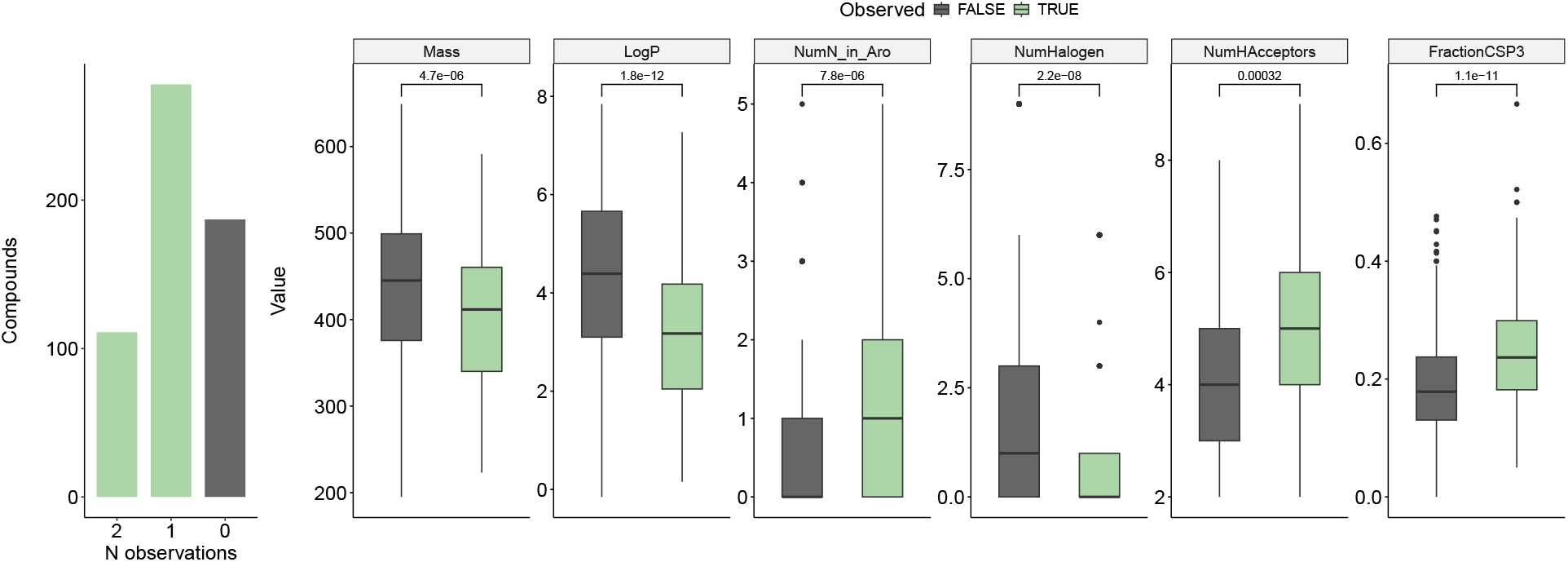
Tag-peptide conjugate observability and annotation. The number of tag candidates seen on both model peptides (111), on only one model peptide (278), and on neither model peptide (187). The number of tags seen on at least one peptide is the sum of the values for N observations = 1 and 2 (389). Chemical descriptors that are significantly differential (*ρ* displayed for Wilcoxon Rank-Sum Test between groups) with respect to observability are shown as box plots.

**Supplemental Fig. 6.**
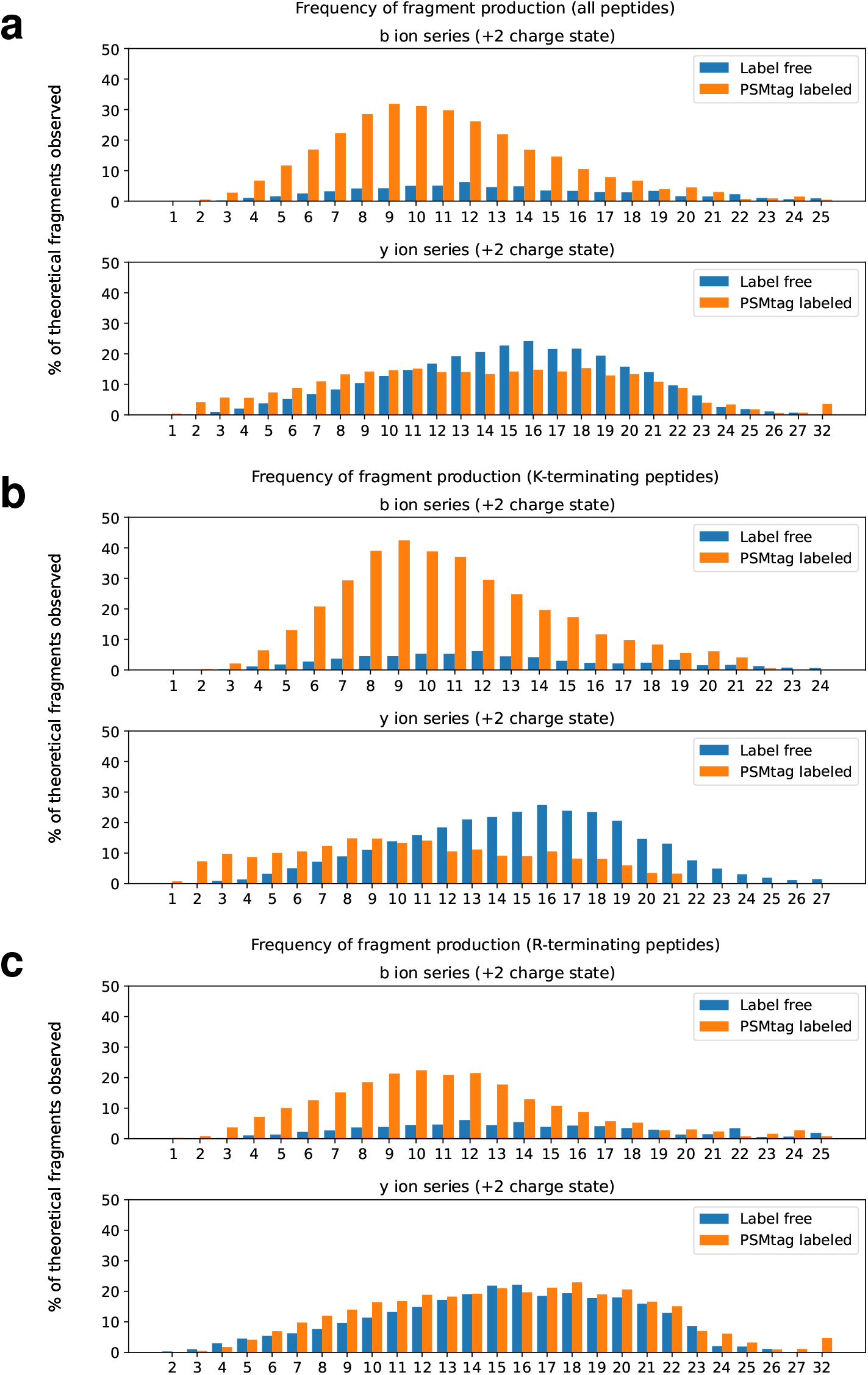
Fragment ion series with two positive charges for tagged and label-free peptides. **a**, The frequency of observing b-ions and y-ions with two positive charges for all peptides confidently assigned. **b**, The subset terminating in lysine. **c**, The subset terminating in arginine. Data was acquired on the Orbitrap Exploris 240 platform using collision energies and FAIMS CV values optimized to maximize peptide identifications from the different sample types, label-free and labeled.

**Supplemental Fig. 7.**
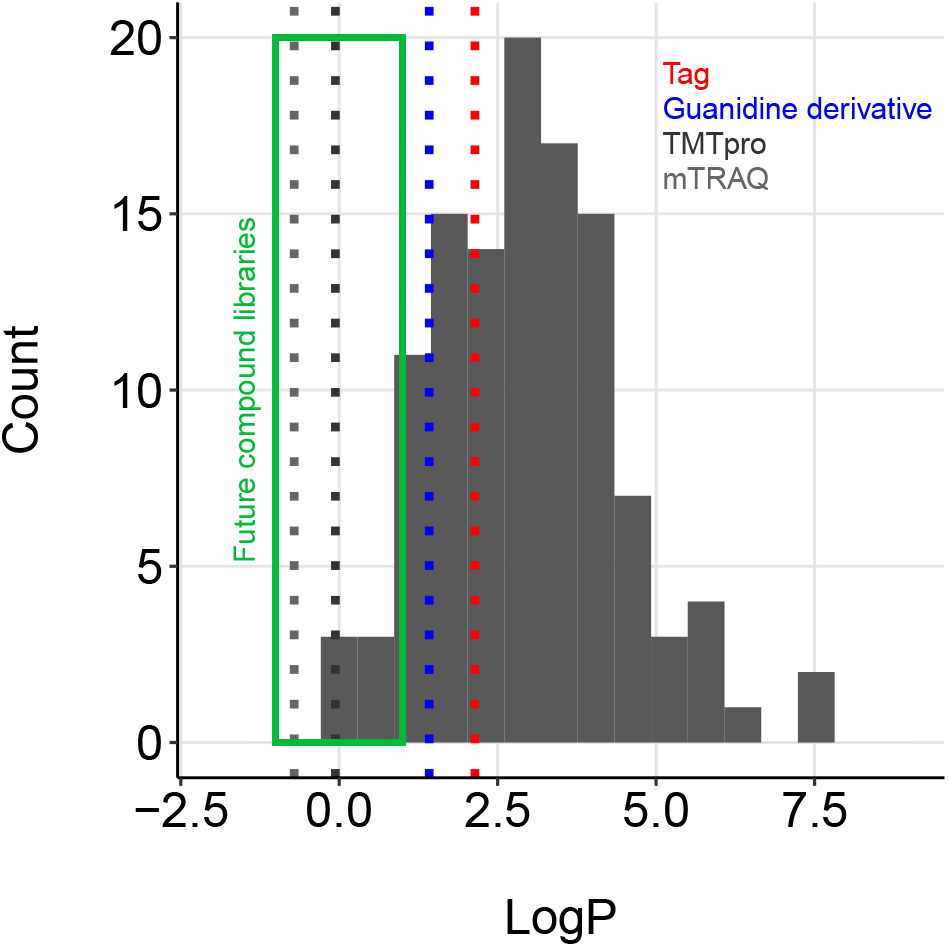
The distribution of logP values for mass tag candidates and commercial mass tags. The logP values for compounds were estimated from SMILES strings using the python package *rdkit*.

**Supplemental Fig. 8.**
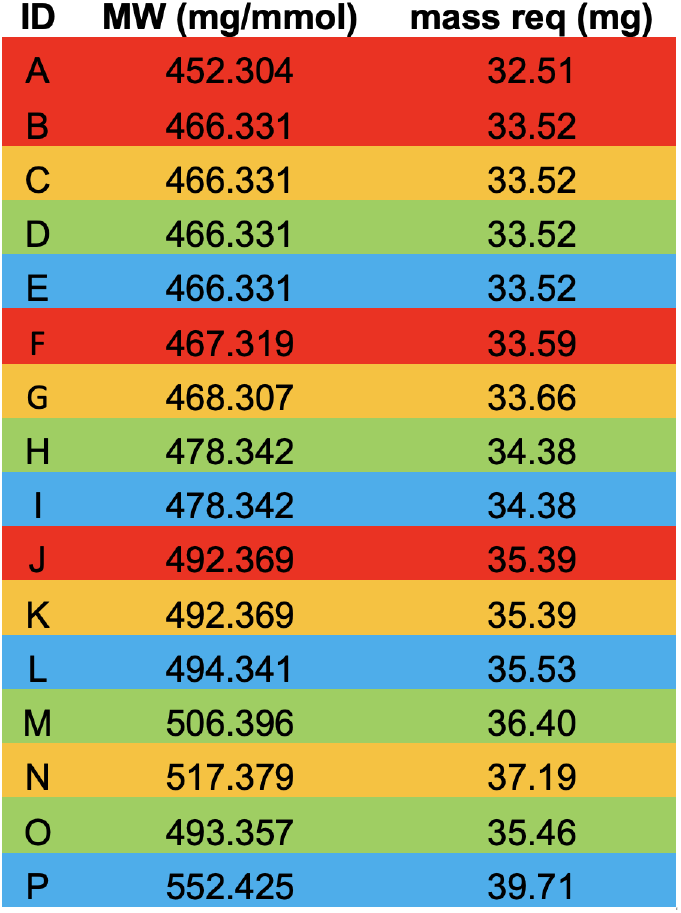
The pooling-scheme for increasing LC-MS/MS throughput of combinatorially-synthesized mass tag candidates. To increase throughput of analyzing mass tag-model peptide conjugates by LC-MS/MS those candidates with different masses were combined.

**Supplemental Fig. 9.**
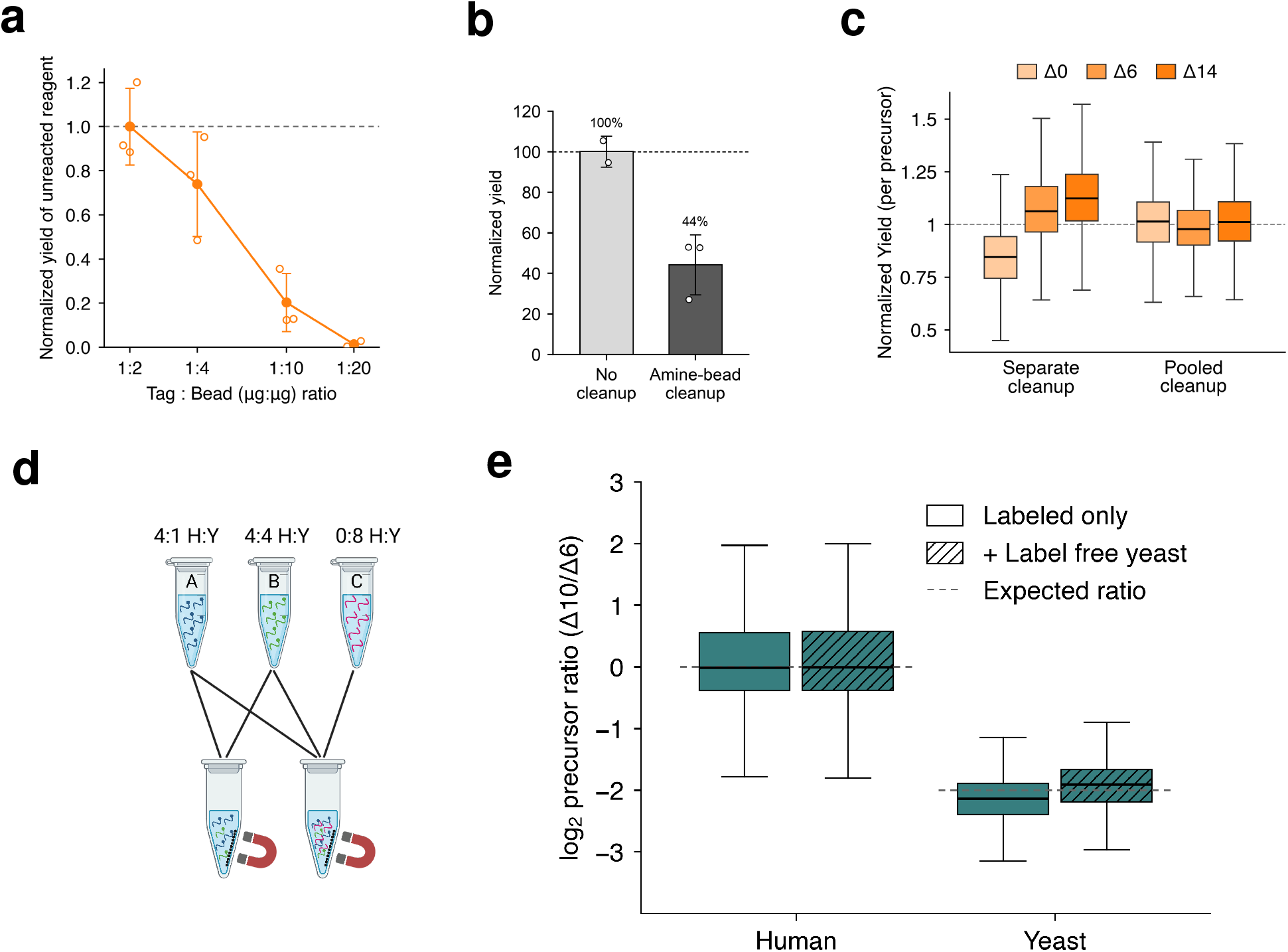
Optimization of amine-bead removal of unreacted PSMtag reagent after peptide labeling. **a**, Residual PSMtag remaining following quenching with amine-functionalized beads using various tag:bead ratios. Values are calculated as the peak area normalized to the highest ratio of tag to beads that was designed to result in incomplete removal (n=3). **b**, The peptide recovery after amine-bead cleanup, normalized to the mean peptide levels from the sample samples with no cleanup. **c**, Recovery yield characterized in separate vs. pooled cleanups. The yield was normalized per-precursor to the mean intensity across all three channels within the two groups. **d**, The experimental design for testing to what extent cross-labeling of unlabeled, highly-abundant material may occur during bead cleanup. **e**, Testing to what extent cross-labeling of unlabeled, highly-abundant material may impact quantitative accuracy and precision. Yeast precursor intensities were normalized to median intensity of human precursors in their respective channels. Precision and accurate was maintained for ratios were calculated between A and B for the combined quench with and without channel C.

## Synthesis Scheme

The tag described to this study is referred to as “Tag 6” in the descriptions on the following pages. Reagents for plex channels 1-9 are denoted as T0, T2, T4, T6, T8, T10, T12, T14, T16, corresponding to their additional nominal mass.

### LCMS method

Column: Agilent Zorbax SB-C18, 3.5 uM, 4.6×50 mm, flow rate 1 mL/min, 0.1% Formic acid ACN/H2O. Gradient: 0.5 min at 2% ACN, linear ramp to 100% ACN over the next 7 min, hold at 100% ACN for 2.5 min. Return to 2% ACN over the next 0.1 min.

**Synthesis schemes for isotopologs and solution phase synthesis of PSMtag.**

Preparation of **methyl 2-(1-acetyl-5-bromoindolin-3-yl)acetate**

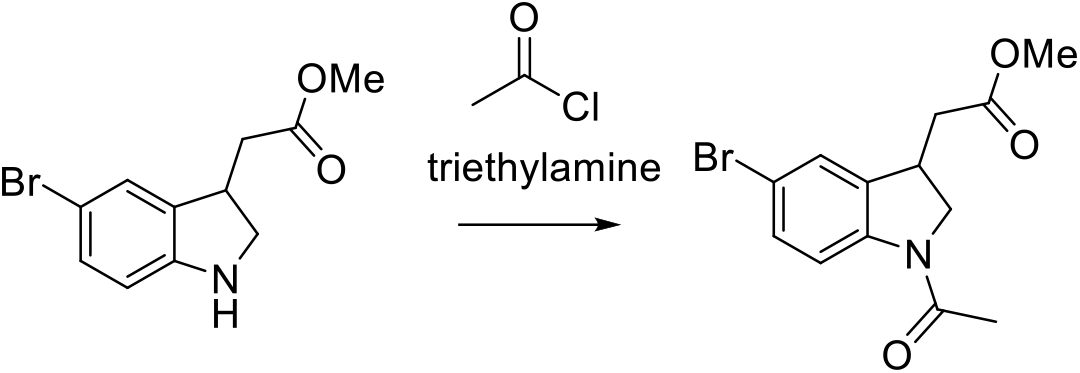

To a stirred mixture of methyl 2-(5-bromo-2,3-dihydro-1H-indol-3-yl)acetate hydrochloride (0.5 g, 1.63 mmol), triethylamine (455 μL, 3.26 mmol) in CH_2_Cl_2_ (8 mL) at 0° C acetyl chloride (139 μL, 1.96 mmol) in CH_2_Cl_2_ (2 mL) was added dropwise over 10 min. The reaction mixture was stirred for 2 hours under a nitrogen atmosphere. After completion of the reaction as indicated by TLC, the reaction was quenched with cold water, and saturated NaHCO_3_. (10 ml) The organic layer was extracted with CH_2_Cl_2_ (3 x 30 mL). The combined organic layers were washed brine solution (20 mL) and dried over Na_2_SO_4_. The solvent was removed under reduced pressure to give crude product. The crude residue was purified by column chromatography (Hexane/EtOAc, 5:5) to provide methyl 2-(1-acetyl-5-bromoindolin-3-yl)acetate (0.47 g, 92.3%) as off -white solid. **^1^H NMR** (300 MHz, CDCl_3_) δ 8.10 (d, *J* = 8.4 Hz, 1H), 7.34 (dd, *J* = 8.4 Hz, 2H), 4. 32 (t, *J* = 8.7 Hz, 1H), 3.78 −3.73(m, 5H), 2.85 – 2.78 (m, 1H), 2.61 – 2.52 (m, 1H), 2.21 (s, 3H); [M+H]^+^: 312.02.

Preparation of **5-bromo-2-(methoxy-^13^C)pyridine**

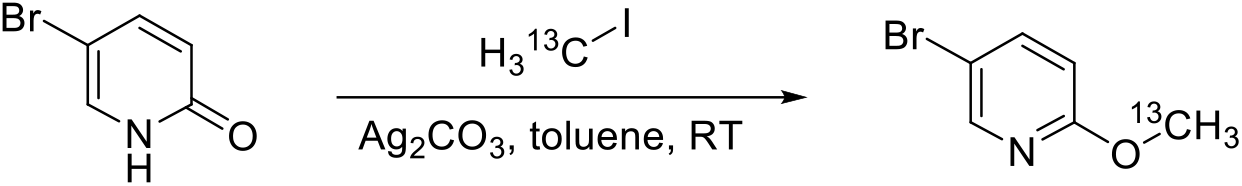

To the suspension of 5-Bromo-1*H*-Pyridin-2-one (550.0 mg, 1 Eq, 3.161 mmol) and silver carbonate (958.8 mg, 1.1 Eq, 3.477 mmol) in Toluene (10 mL) was added Iodo(13C)methane (497.0 mg, 217 μL, 1.1 Eq, 3.477 mmol), and stirred at RT for 3 days. The reaction mixture was filtered through celite to remove the silver salt, and the filer cake was flushed with DCM. The filtrate was concentrated under reduced pressure (∼10 bar) to remove the solvent. The residue was purified by silica gel flash chromatography using 0-3% DCM in hexane as the eluent to obtain 5-bromo-2-(methoxy-13C)pyridine (228.8 mg, 38.2% yield) as a colorless oil. **^1^H NMR** (300 MHz, CDCl_3_-*d*): δ 8.20 (d, *J* = 2.4 Hz, 1H), 7.63 (dd, *J* = 9 Hz, 2.7 Hz, 1H), 6.66 (dd, *J* = 9 Hz, 0.6 Hz, 1H), 3.90 (d, *J* = 146.1 Hz, 3H). **^13^C NMR** (75 MHz, CDCl_3_-*d*): δ 163.17, 147.69, 141.28, 112.84, 111.91, 53.97.

Preparation of **5-bromo-2-(methoxy-^18^O)pyridine**

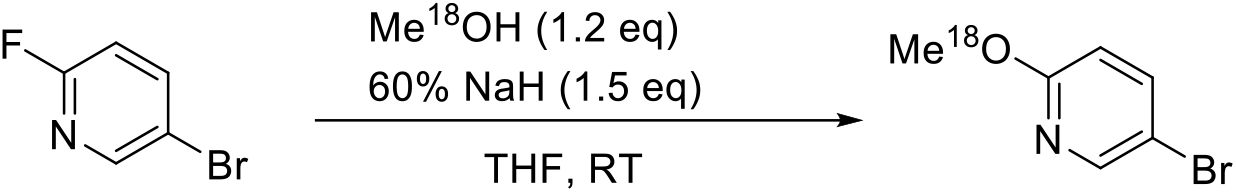

To a solution of methanol-^18^O (578 mg, 690 μL, 1.20 Eq, 17.0 mmol) in anhydrous THF (20 mL) under nitrogen with vigorous stirring was added the NaH (852.3 mg, 60% Wt, 1.5 Eq, 21.308 mmol) at 0 °C. After stirring at RT for 15 min, 5-bromo-2-fluoropyridine (2500.0 mg, 1 Eq, 14.205 mmol) was added dropwise. The resulting mixture was stirred at RT for 4h, then poured into saturated aq. NH_4_Cl and extracted with EtOAc (5×10 mL). The combined organic layers were dried over Na_2_SO_4_, filtered, and concentrated under reduced pressure to obtain a crude product, which was purified by silica gel flash chromatography by using 0-3% DCM in Hexanes as the eluent to afford 5-bromo-2-(methoxy-^18^O)pyridine (2405.1 mg, 89.1 % yield) as a colorless oil. **^1^H NMR** (300 MHz, CDCl_3_-*d*): δ 8.20 (d, *J* = 2.7 Hz, 1H), 7.63 (dd, *J* = 8.7 Hz, 2.7 Hz, 1.5 Hz, 1H), 6.66 (d, *J* = 9.0 Hz, 1H), 3.90 (S, 3H). **^13^C NMR** (75 MHz, CDCl_3_-*d*): δ 163.25, 147.76, 141.22, 112.78, 111.87, 53.85.\

Preparation of **5-bromo-2-(methoxy-^18^O)pyridine-1-^15^*N***

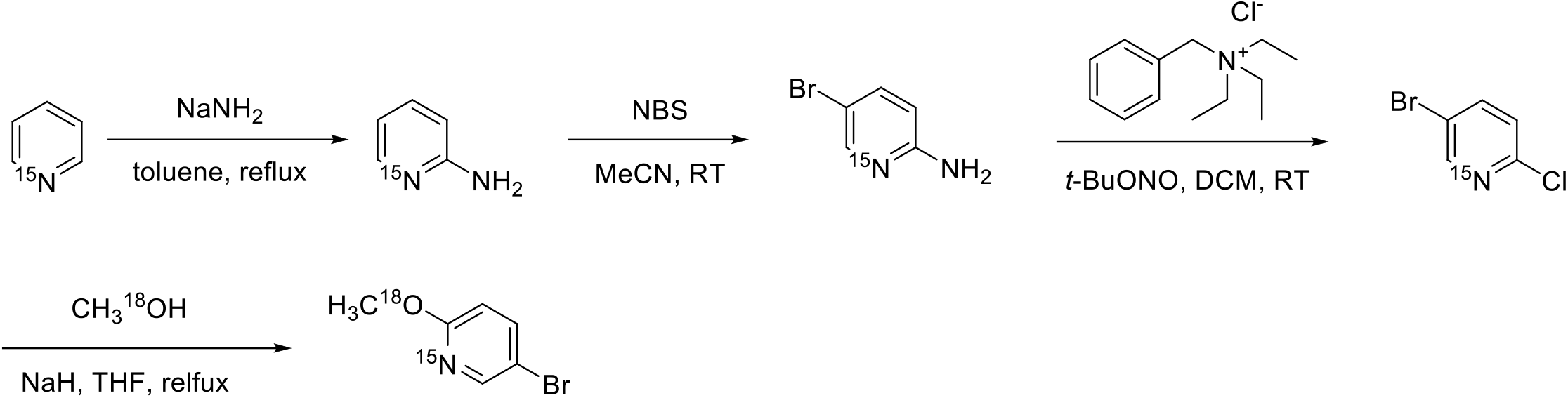

Preparation of **pyridin-1-^15^*N*-2-amine**

To the suspension of sodium amide (1.22 g, 99% Wt, 2.5 Eq, 30.9 mmol) in Toluene (4 mL) in a 40 mL reaction vial at 115 °C with vigorous stirring under Argon atmospheric was slowly added Pyridine-^15^*N* (990 mg, 1.01 mL, 1 Eq, 12.4 mmol) over 5 min, kept heating at 115 °C overnight. After cooling to RT, the residue was slowly quenched by 2 mL of water dropwise, then heated to 70 °C for 20 min. Observed a lot of bubbles during the water addition. Cooled to RT again, separated the lower aqueous solution, and extracted with toluene (6×2 mL). Combined the organic layers, dried over Na_2_SO_4_, filtered, and removed the solvent under reduced pressure to obtain an orange-brown solid as the crude product. The crude product was sublimated under reduced pressure at 90-110°C to obtain pyridin-1-^15^*N*-2-amine (898.3 mg, 76.4 % yield) as a light yellow solid. **^1^H NMR** (300 MHz, CDCl_3_-*d*): δ 8.12 – 7.99 (m, 1H), 7.47 – 7.36 (m, 1H), 6.68 – 6.59 (m, 1H), 6.49 (dd, *J* = 8.4 Hz, 0.6 Hz, 1H), 4.45 (br s, 2H). **^13^C NMR** (75 MHz, CDCl_3_-*d*): δ 158.65, 148.40, 137.85, 114.23, 108.72. **HRMS** m/z calc for C_5_H_7_N^15^N (M+H)^+^: 96.06; found: 95.89.

Preparation of **5-bromopyridin-1-^15^*N*-2-amine**

To the solution of pyridin-1-^15^*N*-2-amine (890.0 mg, 1 Eq, 9.358 mmol) in dry Acetonitrile (20 mL) in an ice water bath was added a solution of NBS (1.699 g, 1.02 Eq, 9.545 mmol) in Acetonitrile (10 mL) dropwise. Slowly warmed up to RT, and stirred the reaction mixture for 2 h at RT. Removed the solvent under reduced pressure, then diluted with sat. NaHCO_3_ aq (35 mL), extracted with EtOAc (4×15 mL). Combined the organic layers, dried over Na_2_SO_4_, filtered, and removed the solvent under reduced pressure. The residue was purified by silica gel flash chromatography using 0-15% EtOAc in DCM as the eluent to obtain 5-bromopyridin-1-^15^*N*-2-amine (1.7007 g, 104.4 % yield) as a yellow solid. **^1^H NMR** (300 MHz, CDCl_3_-*d*): δ 8.09 (dd, *J* = 10.8 Hz, 2.4 Hz, 1H), 7.49 (dd, *J* = 8.4 Hz, 2.4 Hz, 1H), 6.41 (d, *J* = 9.0 Hz, 1H), 4.58 (br s, 2H). **^13^C NMR** (75 MHz, CDCl_3_-*d*): δ 157.30, 148.83, 140.36, 110.23, 108.50. **HRMS** m/z calc for C_5_H_6_BrN^15^N (M+H)^+^: 173.97; found: 174.03.

Preparation **of 5-bromo-2-chloropyridine-1-^15^*N***

To a solution of 5-bromopyridin-1-^15^*N*-2-amine (1000 mg, 1 Eq, 5.747 mmol) in dry DCM (20 mL) was added benzyl(triethyl)ammonium chloride (5.891 g, 4.5 Eq, 25.86 mmol), followed by the addition of tert-Butyl nitrite (5.926 g, 6.84 mL, 10 Eq, 57.47 mmol) in an ice-water bath under an argon atmosphere. After stirring the solution at RT overnight, the reaction mixture was quenched by sat. aqueous NaHCO_3_ solution (35 mL), then extracted with DCM (3×10mL). Combined the organic layers, dried over Na_2_SO_4_, filtered, and removed the solvent under reduced pressure. The residue was purified by silica gel flash chromatography using 0-5% DCM in hexane as the eluent to obtain 5-bromo-2-chloropyridine-1-^15^*N* (484.3 mg, 43.57 % yield) as a white solid. **^1^H NMR** (300 MHz, CDCl_3_-*d*): δ 8.46 (dd, *J* = 11.7 Hz, 2.7 Hz, 1H), 7.77 (dd, *J* = 8.4 Hz, 2.7 Hz, 1H), 7.24 (d, *J* = 8.4 Hz, 1H). **^13^C NMR** (75 MHz, CDCl_3_-*d*): δ 150.93, 150.39, 141.40, 125.83, 119.32.

Preparation of **5-bromo-2-(methoxy-^18^O)pyridine-1-^15^*N***

To a solution of methanol-18O (84.47 mg, 101 μL, 1.2 Eq, 2.482 mmol) in anhydrous THF (6 mL) under nitrogen with vigorous stirring was added NaH (124.1 mg, 60% Wt, 1.5 Eq, 3.102 mmol) at 0 °C. After stirring at 0 °C for 15 min, a solution of 5-bromo-2-chloropyridine-1-^15^*N* (400.0 mg, 1 Eq, 2.068 mmol) in THF (2 mL) was added dropwise. The resulting mixture was stirred at 85 °C for 41h. After cooling to RT, the reaction mixture was poured into saturated aq. NH_4_Cl (30 mL) and extracted with Et_2_O (2×10 mL). Combined the organic layers, dried over Na_2_SO_4_, filtered, and removed the solvent under reduced pressure. The residue was purified by silica gel flash chromatography by using 0-5% DCM in hexanes to afford 5-bromo-2-(methoxy-^18^O)pyridine-1-^15^*N* (317.6 mg, 80.40 % yield) as a colorless oil. The sample solidified at −20°C during storage. **^1^H NMR** (300 MHz, CDCl_3_-*d*): δ 8.19 (ddd, *J* = 11.4 Hz, 2.7 Hz, 0.6, 1H), 7.63 (dd, *J* = 8.7 Hz, 2.7 Hz, 1H), 6.66 (dd, *J* = 9 Hz, 0.9 Hz, 1H), 3.90 (s, 3H). **^13^C NMR** (75 MHz, CDCl_3_-*d*): δ 163.26, 147.73, 141.22, 112.76, 111.90, 53.85.

Intermediate for **T6**, **T8**, and **T10**:

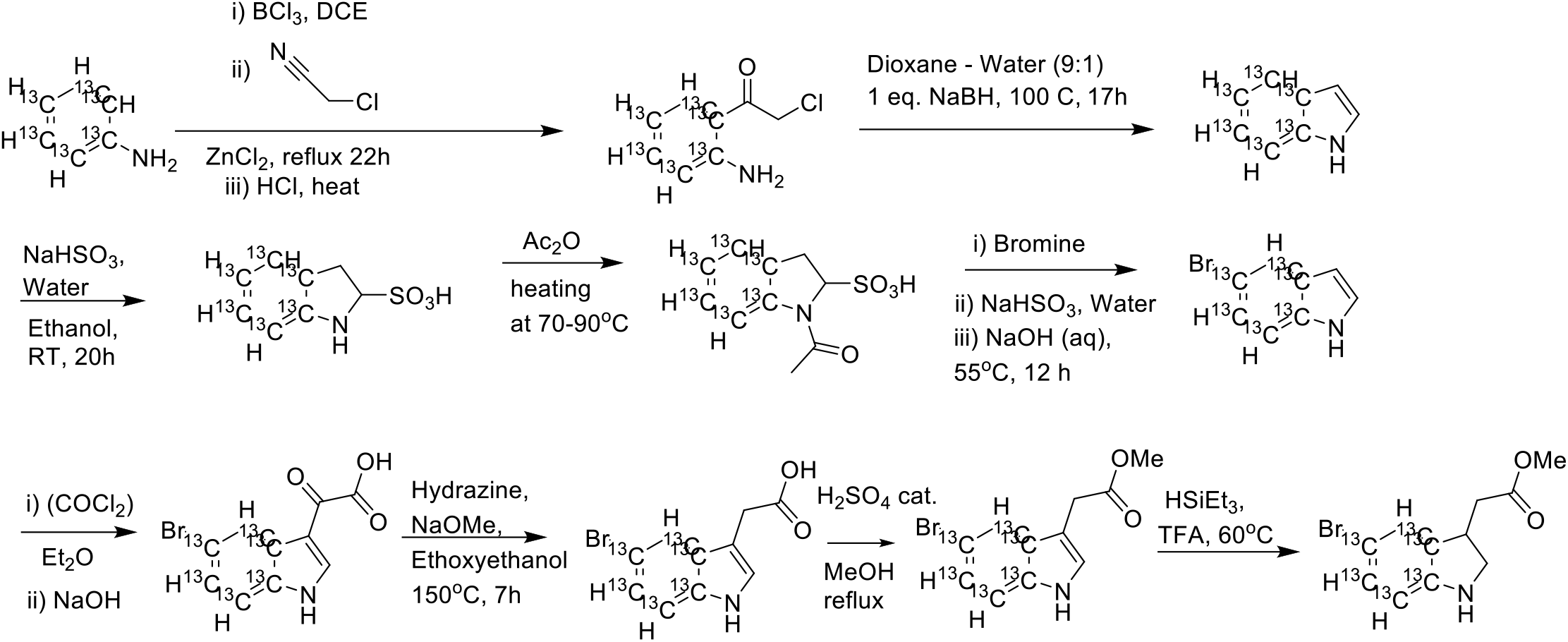

**1-(2-aminophenyl-1,2,3,4,5,6-^13^C6)-2-chloroethan-1-one** was prepared according to the protocol described in Wager, C.A.B; Miller, S.A; J. Label Cmpd Radiopharm 2006; 49; 615-622, LCMS RT 5.23 min, (M+H)^+^: Expected 176.1; observed 176.0

**1H-indole-3a,4,5,6,7,7a-^13^C6** was prepared according to the protocol described in Wager, C.A.B; Miller, S.A; J. Label Cmpd Radiopharm 2006; 49; LCMS RT 5.47 min, (M+H)^+^ Expected MW: 124.1; observed 124.2

**5-bromo-1H-indole-3a,4,5,6,7,7a-^13^C6** was prepared via a sulfonate according to the protocol described in Chinese Patent Suzhou Hande Chuanghong Biochemical Technology Co., Ltd.;Mao Zhongping;Ma

Dongxu;Jiang Xiangsheng CN110396059, 2019. LCMS RT 6.08 min, (M-H)^-^: Expected 200.0/202.0; observed 200.2/202.1

**2-(5-bromo-1H-indol-3-yl-3a,4,5,6,7,7a-^13^C6)-2-oxoacetic acid** was prepared according to the protocol described in the Japanese Patent (2022) 7125719 B2, WO2018164214 A1 2018-09-13, LCMS RT 4.75 min, (M-H)^-^: Expected 272.0, 274.0, observed 272.0, 274.0

**2-(5-bromo-1H-indol-3-yl-3a,4,5,6,7,7a-^13^C6)acetic acid** was prepared according to the protocol described in the Japanese Patent (2022) 7125719 B2, WO2018164214 A1 2018-09-13, LCMS RT 5.08 min, (M-H)^-^: Expected 258.0, 260.0; observed 258.1, 260.1

**Preparation of Methyl 2-(5-bromo-1H-indol-3-yl-3a,4,5,6,7,7a-^13^C6)acetate methyl 2-(5-bromo-1H-indol-3-yl-3a,4,5,6,7,7a-13C6)acetate**

The indole carboxylic acid was dissolved in methanol and catalytic amount of sulfuric acid, and the resultant solution was heated at reflux overnight and then evaporated to dryness and used directly in the next step. LCMS RT 6.00 min, (M-H)^-^: expected 272.0, 274.0; observed 272.1, 274.3

**Methyl 2-(5-bromoindolin-3-yl-3a,4,5,6,7,7a-^13^C6)acetate** was prepared according to the protocol described in Phytochemistry 68 (2007) 2512–2522. LCMS RT 5.37 min, (M+H)^+^: expected 276.0, 278.0; observed 276.0, 278.0

Intermediate for **T12**, **T14**, and **T16:**

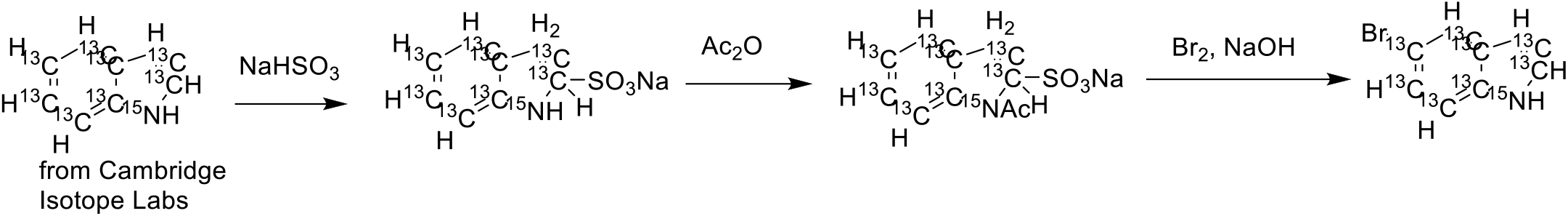

**5-bromo-1H-indole-^13^C8-^15^N** was prepared via a sulfonate according to the protocol described in Chinese Patent Suzhou Hande Chuanghong Biochemical Technology Co., Ltd.;Mao Zhongping;Ma Dongxu;Jiang Xiangsheng CN110396059, 2019.

Intermediate for **T12**:

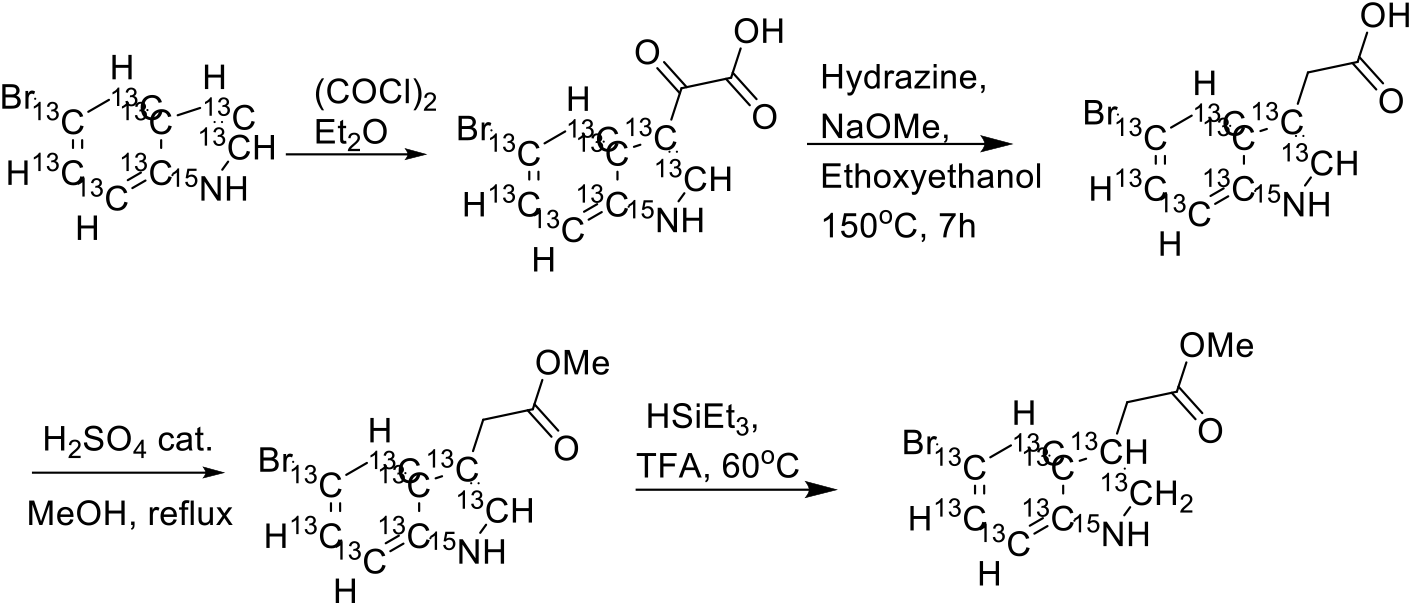

**2-(5-Bromo-1H-indol-3-yl-^13^C8-^15^N)-2-oxoacetic acid** was prepared according to the protocol described in the Japanese Patent (2022) 7125719 B2, WO2018164214 A1 2018-09-13. LCMS RT 4.78 min, (M-H)^-^: 275.0, 277.0; observed 275.2, 277.1

**2-(5-Bromo-1H-indol-3-yl-^13^C8-^15^N)acetic acid** was prepared according to the protocol described in the Japanese Patent (2022) 7125719 B2, WO2018164214 A1 2018-09-13. LCMS RT 5.15 min, (M-H)^-^: Expected 261.0, 263.0, observed 261.1, 263.1

Preparation of **Methyl 2-(5-bromo-1H-indol-3-yl-^13^C8-^15^N)acetate**.

The indole carboxylic acid was dissolved in methanol and catalytic amount of sulfuric acid, and the resultant solution was heated at reflux overnight. The mixture was evaporated to dryness and used directly in the next reaction. LCMS RT 5.90 min, (M-H)^-^: Expected 275.0, 277.0; observed 275.1, 277.0

**methyl 2-(5-bromoindolin-3-yl-13C8-15N)acetate** was prepared according to the protocol described in Phytochemistry 68 (2007) 2512–2522. LCMS RT 5.30 min, (M+H)^+^: Expected 279.0, 281.0, observed 279.0, 281.0

Intermediate for **T14** and **T16:**

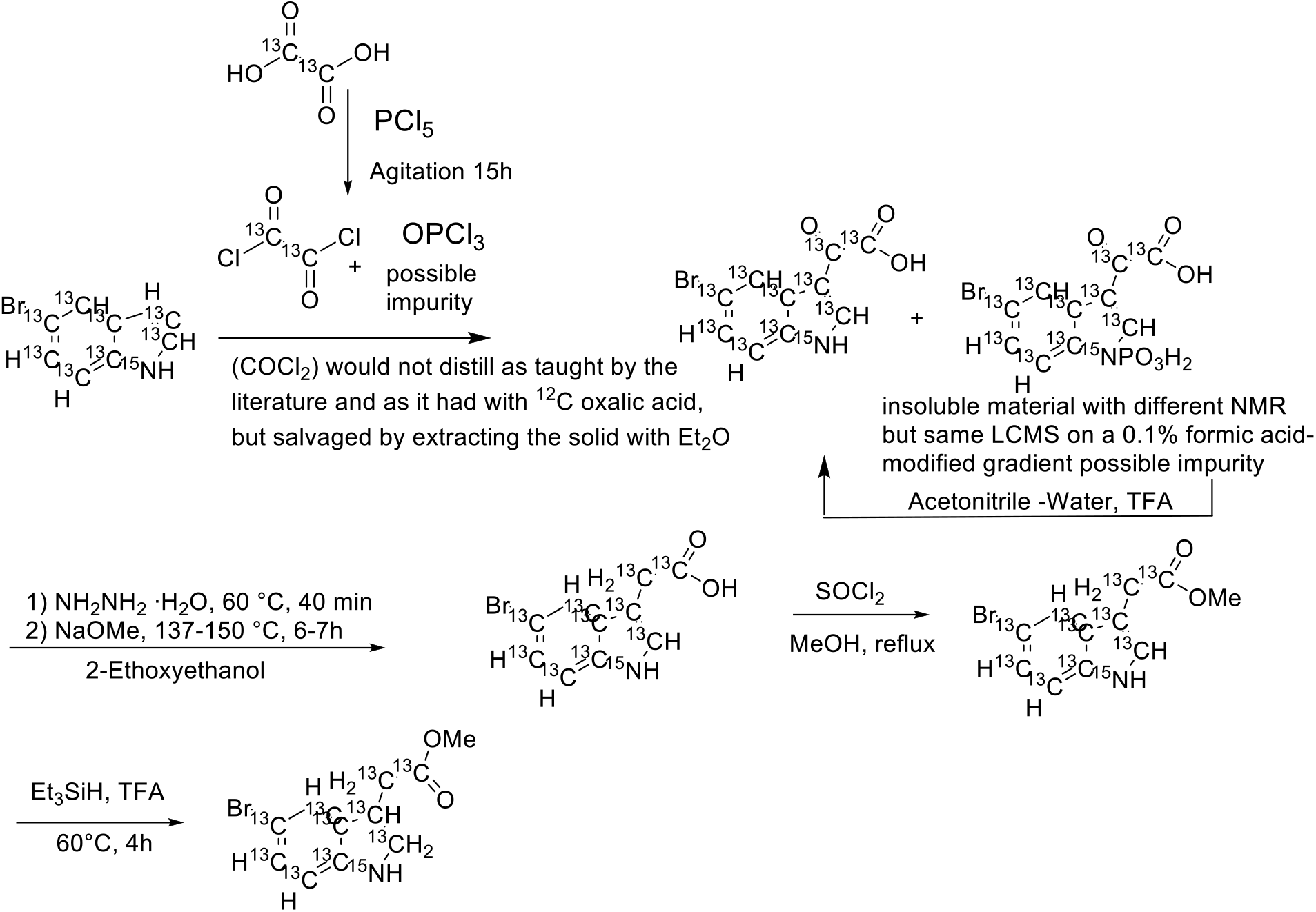

Preparation of **2-(5-bromo-1H-indol-3-yl-^13^C8-^15^N)-2-oxoacetic-^13^C2 acid**

^13^C-Oxalic acid (purchased from Cambridge isotopes lab) and Phosphorus pentachloride (5.4 g) was mixed in a 5 ml sealed vial and gently agitated with a vortex at RT for 15 hours. Hickman head distillation set up for distillation of labelled Oxalyl chloride failed, even though the reaction mixture liquified and this procedure worked multiple times reliably using ^12^C-oxalic acid. Oxalyl chloride is volatile but nothing was collected in the Hickman head receiver. Ether 5 ml was added into the reaction mixture to extract the ^13^C oxalyl chloride, and this was cooled to 0 C and added to 350 mg of 5-bromo-1H-indole-^13^C8-^15^N, in 10 mL diethyl ether. The resulting reaction mixture was allowed to warm to room temperature and stirred overnight. A precipitate formed and was collected by filtration. Both the precipitate and the filtrate showed an LCMS profile consistent with the desired product. The precipitate was hypothesized to contain an N-P bond resulting from residual POCl3 from the oxalyl chloride formation step, and the precipitate was treated with 10% water, 70% acetonitrile, 20% TFA, resulting in a solution. The mixture was stirred 4h and evaporated to dryness. 110 mg (23%) LCMS RT 4.88 min, (M-H)^-^: Expected 277.0, 279.0; observed 277.2, 279.2

**2-(5-bromo-1H-indol-3-yl-^13^C8-^15^N)acetic-^13^C2 acid** was prepared according to the protocol described in Japanese Patent (2022) 7125719 B2 WO2018164214 A1 2018-09-13 LCMS RT 5.09 min, (M-H)^-^: expected 263.0, 265.0; observed 263.1, 265.1

**Methyl 2-(5-bromo-1H-indol-3-yl-^13^C8-^15^N)acetate-^13^C2**. The indole carboxylic acid was dissolved in methanol and catalytic amount of sulfuric acid, and the resultant solution was heated at reflux overnight and then evaporated to dryness and used directly in the next step. LCMS RT 5.90 min, (M-H)^-^: expected MW: 277.0, 279.0; observed 277.1, 279.1

**Methyl 2-(5-bromoindolin-3-yl-^13^C8-^15^N)acetate-^13^C2** was prepared according to the protocol described in Phytochemistry 68 (2007) 2512–2522. LCMS RT 5.51 min, (M+H)^+^: Expected MW: 281.0, 283.0, observed (ES+) 283.1

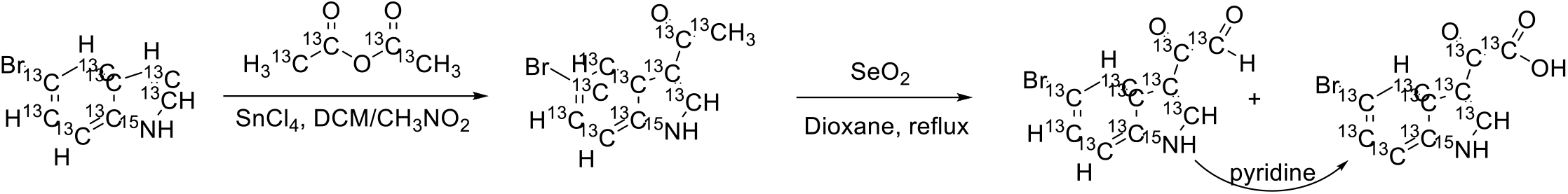

Preparation of **1-(5-bromo-1H-indol-3-yl-^13^C8-^15^N)ethan-1-one-1,2-^13^C2**

To a stirring solution of 5-bromo-1H-indole-^13^C8-^15^N (400 mg, 1 Eq, 1.95 mmol) in DCM (4 mL) under argon at 0 °C was added tin(IV) chloride (610 mg, 275 μL, 1.2 Eq, 2.34 mmol) in a single portion via syringe, and the mixture was stirred at 0 °C for 5 min. A light yellow precipitate formed. After the ice bath was removed, the mixture was stirred at room temperature for 30 min, at which point the mixture became a green solution. A solution of acetic anhydride-1,1,2,2-^13^C4 (207 mg, 185 μL, 1 Eq, 1.95 mmol) in nitromethane (3 mL) was added dropwise. The mixture was stirred for 2 h at room temperature. After being quenched with ice (10 g), the mixture was extracted with ethyl acetate (5×6 mL). The organic phase was dried over Na_2_SO_4_ and concentrated at reduced pressure to give the crude product. The crude product was slurried in Et_2_O and filtered to collect the off-white solid. The solid and filtrate were purified separately by on C18 silica using 3-100% MeCN in water with 0.1% FA as the eluent to afford 1-(5-bromo-1H-indol-3-yl-^13^C8-^15^N)ethan-1-one-1,2-^13^C2 (331.1 mg, 1.330 mmol, 68.1 %). LCMS rt 5.09 min, [M+H]^+^ expected 249.0/251.0, observed 249.2/251.2

Preparation of **2-(5-bromo-1H-indol-3-yl-^13^C8-^15^N)-2-oxoacetic-^13^C2 acid**

To a suspension of 1-(5-bromo-1H-indol-3-yl-^13^C8-^15^N)ethan-1-one-1,2-^13^C2 (330 mg, 1 Eq, 1.33 mmol) in 1,4-Dioxane (30 mL) was added selenium dioxide (221 mg, 1.5 Eq, 1.99 mmol), and the mixture was heated at reflux overnight. LC-MS indicated predominant formation of 2-(5-bromo-1H-indol-3-yl-13C8-15N)-2-oxoacetaldehyde-13C2 to the reaction mix was added 1 mL of pyridine, and the mixture was heated at reflux overnight. After cooling to room temperature, the mixture was filtered to remove the SeO2, and the filtrate was concentrated under reduced pressure to obtain the crude product as a pink solid. The crude product was dissolved in 100 mL of 1N NaOH aq, extracted with ethyl acetate (3 × 15 mL). The aqueous layer was acidified with 6 N HCl solution to pH=3 and extracted with ethyl acetate (5 × 20 mL). The aqueous layer was then acidified with 12 N HCl solution to pH=1 and extracted with ethyl acetate (2 × 20 mL). All the organic layers were combined, dried over anhydrous Na_2_SO_4_ and concentrated under reduced pressure to obtain 2-(5-bromo-1H-indol-3-yl-^13^C8-^15^N)-2-oxoacetic-^13^C2 acid (320 mg, 1.15 mmol, 86.5 %) as a yellow solid. LCMS rt 4.63 min, ES-expected 277.0/279.0, observed 277.0/279.0.

Preparation of T0:

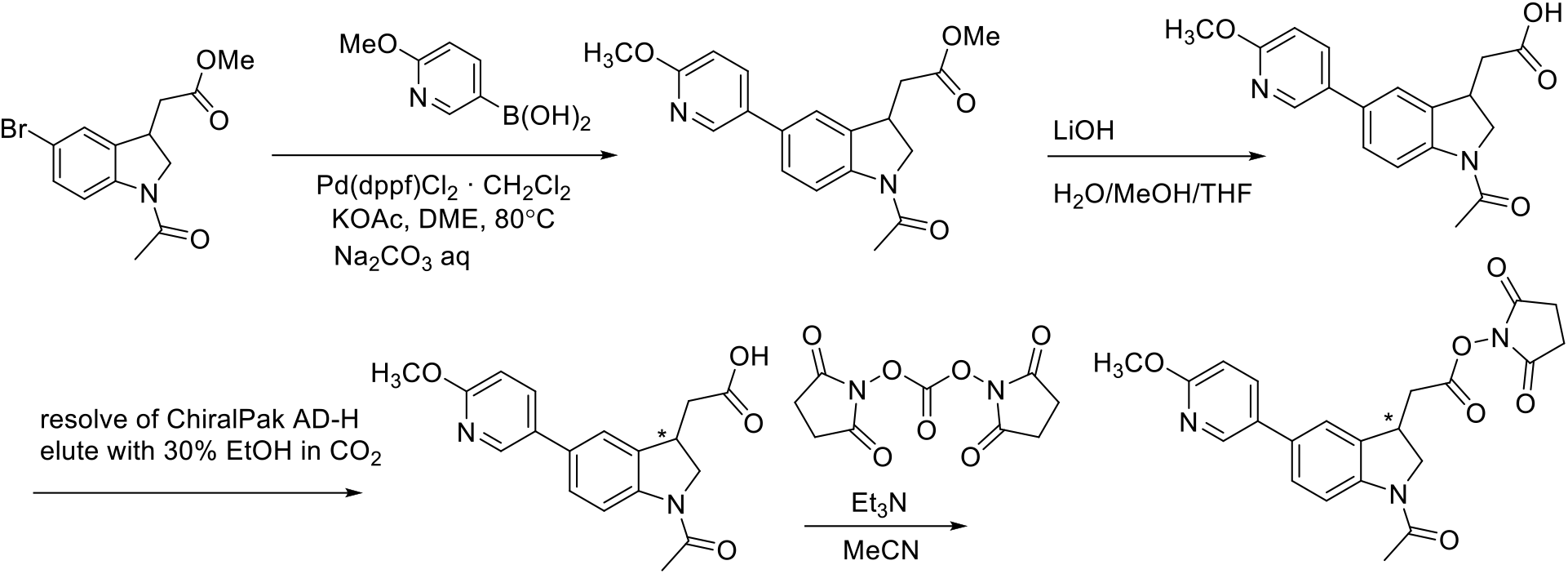

**Preparation of Methyl 2-(1-acetyl-5-(6-methoxypyridin-3-yl)indolin-3-yl)acetate**

In a 50 mL glass vial containing a magnetic stir bar, methyl 2-(1-acetyl-5-bromoindolin-3-yl)acetate **2** (0.25 g, 0.801 mmol) and 2-methoxy-5-pyridinylboronic acid **3** (0.147g, 0.961 mmol) were dissolved in 4 mL of DME (dimethoxyethane) and 1.2 mL of water. The vial was purged with nitrogen for 10 minutes. [1,1’-Bis(diphenylphosphino)ferrocene]dichloropalladium(II)ComplexWithDichloromethane (0.0327 g, 0.0401 mmol), and sodium carbonate (0.170 g, 1.60 mmol) were then added, and the vial was purged with nitrogen for an additional 10 minutes. The resultant solution was stirred at room temperature for 5 minutes, then the vial was capped, heated to 80°C, and stirred for 60 minutes. LCMS analysis of the reaction mixture showed a major peak corresponding to the desired product, indicating that the starting material was completely consumed. The reaction mixture was filtered through a celite pad, and the solvent was concentrated under reduced pressure using a rotary evaporator. The crude residue was purified by column chromatography (Hexane/EtOAc, 2:8) to provide Methyl 2-(1-acetyl-5-(6-methoxypyridin-3-yl)indolin-3-yl)acetate (0.240 g, 88%) as off-white solid. **^1^H NMR** (300 MHz, CDCl_3_) δ 8.33 (s, 1H) 8.26 (d, *J* = 8.1 Hz, 1H), 7.74 (d, *J* = 8.4 Hz, 1H), 7.39 (d, *J* = 8.7 Hz, 1H), 7.29 (s, 1H), 6.80 (d, *J* = 9.0 Hz, 1H), 4. 37 (t, *J* = 9.3 Hz, 1H), 3.97 (s, 3H) 3.90-3.74 (m, 5H), 2.92 – 2.87 (m, 1H), 2.66 – 2.57 (m, 1H), 2.25 (s, 3H)

**Preparation of 2-(1-acetyl-5-(6-methoxypyridin-3-yl)indolin-3-yl)acetic acid**

Methyl 2-(1-acetyl-5-(6-methoxypyridin-3-yl)indolin-3-yl)acetate (0.24 g, 0.705 mmol) was dissolved in methanol (6 mL) and water (2 mL), and LiOH (0.57g, 2.12 mmol) was added. The reaction mixture was stirred at room temperature for 1 hour. LCMS analysis indicated that the starting material was consumed, and the desired acid peak was observed. The methanol was removed under reduced pressure using a rotary evaporator, and 3 mL of water was added. The solution was neutralized with 4 N HCl until the pH reached 4. A solid formed, which was filtered and dried under vacuum to give 2-(1-acetyl-5-(6-methoxypyridin-3-yl)indolin-3-yl)acetic acid (0.206 g, 90%) as off -white solid. **^1^H NMR** (300 MHz, DMSO-D6) δ 8.24 (s, 1H) 8.06 (d, *J* = 8.1 Hz, 1H), 7.95 (d, *J* = 8.7 Hz, 1H), 7.59 (s, 1H), 7.46 (d, *J* = 8.1 Hz, 1H), 6.87 (d, *J* = 8.7 Hz, 1H), 4. 34 (t, *J* = 9.6 Hz, 1H), 3.86-3.70 (m, 5H), 2.97 – 2.91 (m, 1H), 2.61 – 2.55 (m, 1H), 2.15 (s, 3H); LCMS rt 4.87 min, [M+H]^+^: expected 327.1, observed 327.1

**Preparation of 2,5-Dioxopyrrolidin-1-yl 2-(1-acetyl-5-(6-methoxypyridin-3-yl)indolin-3-yl)acetate (racemic)**

2-(1-acetyl-5-(6-methoxypyridin-3-yl)indolin-3-yl)acetic acid **5** (10 mg, 0.031 mmol), N,N’-Disuccinimidyl Carbonate (8.7 mg, 0.034 mmol), and TEA (13 μL, 0.096 mmol) were dissolved in 0.5 mL of CH₃CN. The reaction was stirred for 1 hour at room temperature. LCMS indicated the consumption of the starting material. The reaction mixture was then diluted with 10 mL of DCM and 10 mL of 0.1 M KH_2_PO₄ solution, followed by extraction with DCM (3 × 20 mL). The combined organic layers were washed brine solution (20 mL) and dried over Na_2_SO_4_. The solvent was removed under reduced pressure to give crude product. The crude residue was purified by column chromatography (DCM/EtOAc, 5:5) to provide **2**. The product was dissolved in 5 mL of acetonitrile, lyophilized, and give 2,5-dioxopyrrolidin-1-yl 2-(1-acetyl-5-(6-methoxypyridin-3-yl)indolin-3-yl)acetate (0.0105 g, 80%) as off -white solid. **^1^H NMR** (300 MHz, CDCl_3_) δ 8.35 (s, 1H) 8.28 (d, *J* = 8.4 Hz, 1H), 7.76 (dd, *J* = 8.4 Hz, 1H), 7.40 (d, *J* = 8.1 Hz, 1H), 7.37 (s, 1H), 6.81 (d, *J* = 8.7 Hz, 1H), 4. 37 (t, *J* = 9.6 Hz, 1H), 3.98 (s, 3H) 3.97-3.88 (m, 2H), 3.2 – 3.12 (m, 1H), 2.94 – 2.91 (m, 1H), 2.90 – 2.88 (m, 4H), 2.26 (s, 3H); LCMS rt 5.25 min, [M+H]^+^: expected 424.1, observed 424.0

**Resolution:** ChrialPak AD-H 21×250 mm, with 30% ethanol in CO_2_ and a flow rate of 70 mL/min. Sample dissolved at 2 mg/mL in 50:50 dichloromethane:methanol and loaded at 2 mL per injection to afford 22.4 mg of peak 1, >99.5% purity and ee, 23.2 mg of peak 2 with >99.5% purity and 99% ee.

**Peak 1: 2,5-Dioxopyrrolidin-1-yl 2-(1-acetyl-5-(6-methoxypyridin-3-yl)indolin-3-yl)acetate**

To a suspension of 2-(1-acetyl-5-(6-methoxypyridin-3-yl)indolin-3-yl)acetic acid (22.4 mg, 1 Eq, 68.6 μmol) in MeCN (0.5 mL) in a 40 mL vial was added N.N’-Disuccinimidyl Carbonate (21.1 mg, 0.02 mL, 1.2 Eq, 82.4 μmol) followed by N.N’-Disuccinimidyl Carbonate (21.1 mg, 0.02 mL, 1.2 Eq, 82.4 μmol). The reaction mixture became a colorless clear solution after adding the E3N. The mixture was stirred at room temperature for 1h, and LC-MS showed ∼13% of acid remaining. 3.6 mg of N.N’-Disuccinimidyl Carbonate was added, stirred at RT for 30min. LC-MS showed ∼40% of SM remained. 15 mg of N.N’-Disuccinimidyl Carbonate was added, stirred at RT for 1h. The mixture was diluted with 1 mL of brine and 1 mL of NaHCO3 aq in the vial, extracted with DCM (4*1 mL, centrifuged to get a better separation in the vial). A pipet was used to remove the DCM out of the vial and all the organic layers were combined in a new vial, dried over Na2SO4, rotavap to dry. The residue was purified on silica gel (dry loading) eluting with 0-100% DCM in hexane then 0-35% EtOAc in DCM to get the pure product 10.7 mg as a white solid after freeze-drying. LCMS rt 5.53 min, [M+H]^+^: expected 424.1, observed 424.3.

**Peak 2: 2,5-Dioxopyrrolidin-1-yl 2-(1-acetyl-5-(6-methoxypyridin-3-yl)indolin-3-yl)acetate**

2-(1-Acetyl-5-(6-methoxypyridin-3-yl)indolin-3-yl)acetic acid (23 mg, 0.070 mmol), N,N’-Disuccinimidyl Carbonate (20 mg, 0.077 mmol), and TEA (31 μL, 0.22 mmol) were dissolved in 0.5 mL of CH₃CN. The reaction was stirred for 1 hour at room temperature. LCMS indicated the consumption of the starting material. The reaction mixture was then diluted with 10 mL of water 1 mL of saturated NaHCO_3_ solution, followed by extraction with DCM (3 × 20 mL). The combined organic layers were washed brine solution (20 mL) and dried over Na_2_SO_4_. The solvent was removed under reduced pressure to give crude product. The crude residue was purified by column chromatography (DCM/EtOAc, 5:5) to provide **T0-peak2**. The product was dissolved in 5 mL of acetonitrile, lyophilized, and afforded **T0-peak2** (0.015 g, 51%) as off -white solid. **^1^H NMR** (300 MHz, CDCl_3_) δ 8.35 (s, 1H) 8.28 (d, *J* = 8.1 Hz, 1H), 7.77 (dd, *J* = 8.4 Hz, 1H), 7.42 (d, *J* = 8.1 Hz, 1H), 7.37 (s, 1H), 6.81 (d, *J* = 8.7 Hz, 1H), 4. 37 (t, *J* = 8.7 Hz, 1H), 3.98 (s, 3H) 3.97-3.88 (m, 2H), 3.2 – 3.12 (m, 1H), 2.94 – 2.91 (m, 1H), 2.90 – 2.88 (m, 4H), 2.27 (s, 3H); LCMS rt 5.38 min, [M+H]^+^: expected 424.1, observed 424.3.

**T2:**

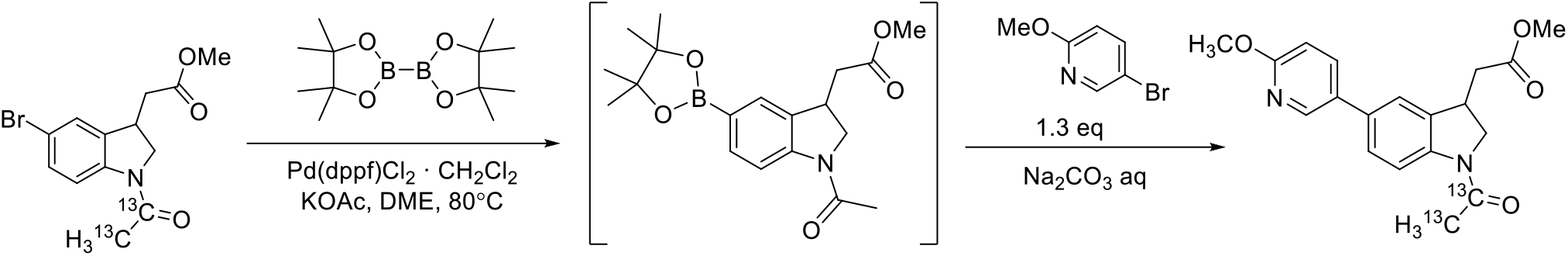

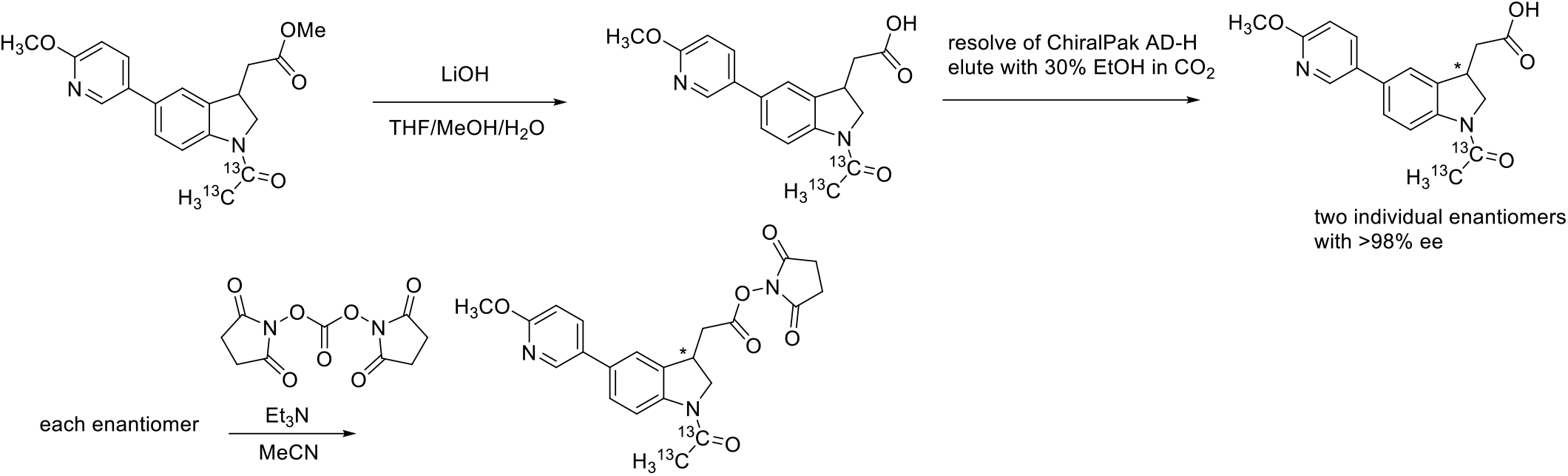

**Preparation of Methyl 2-(1-(acetyl-^13^C2)-5-(6-methoxypyridin-3-yl)indolin-3-yl)acetate**

To the solution of methyl 2-(1-(acetyl-^13^C2)-5-bromoindolin-3-yl)acetate (150.0 mg, 1 Eq, 477.5 μmol) in DME (3.6 mL) was added KOAc (93.72 mg, 2 Eq, 955.0 μmol) and Bis(pinacolato)diborane (181.9 mg, 1.5 Eq, 716.2 μmol), degassed and recharged with Argon for 3 times, then added Pd(dppf)Cl_2_·CH_2_Cl_2_ (38.99 mg, 0.1 Eq, 47.75 μmol). The mixture was heated at 80 °C under Argon atmosphere for 70min. After cooling to RT, a solution of Na_2_CO_3_ (101.2 mg, 2 Eq, 955.1 μmol) in Water (1.8 mL) was added to the reaction mixture, followed by the solution of 5-bromo-2-methoxypyridine (114.5 mg, 1.3 Eq, 609.0 μmol) in DME (1.8 mL). After heating at 80 °C under Argon atmosphere for 2.5 hours, the solvent was removed under reduced pressure. The residue was purified by silica gel flash chromatography by using 0-40% EtOAc in hexane as the eluent to obtain the desired product (128.9 mg, 78.8% yield) as a yellow foamy solid. LCMS rt 5.67 min, [M+H]^+^ expected 343.1, observed 343.1.

**Preparation of 2-(1-(Acetyl-^13^C2)-5-(6-methoxypyridin-3-yl)indolin-3-yl)acetic acid**

To the solution of Methyl 2-(1-(acetyl-13C2)-5-(6-methoxypyridin-3-yl)indolin-3-yl)acetate (124.6 mg, 1 Eq, 363.9 μmol) in Methanol (0.5 mL), THF (1 mL), and Water (0.5 mL) was added LiOH (34.87 mg, 4 Eq, 1.456 mmol). The reaction mixture was stirred at RT for 3h, then acidified by 3M HCl aq to adjust the pH=2∼3 to precipitate the product. The suspension was diluted with 2 mL of water, sonicated for 1min, centrifuged, and removed the clear liquid on the upper layer by pipet. Repeated the water wash twice (2×2mL), then used MeCN (3×2mL) instead of water, repeated 3 times. The solid was suspended in 1:1 MeCN/water, freeze-dried to obtain 2-(1-(Acetyl-^13^C2)-5-(6-methoxypyridin-3-yl)indolin-3-yl)acetic acid (98.1 mg, 82.1% yield) as an off-white solid. LCMS rt 4.80 min, [M+H]^+^ expected 329.1, observed 329.1.

**Resolution:** ChrialPak AD-H 21×250 mm, with 30% ethanol in CO_2_ and a flow rate of 70 mL/min. Sample dissolved at 2 mg/mL in 50:50 dichloromethane:methanol and loaded at 2 mL per injection to afford 21.8 mg of peak 1, >99.5% purity and ee, 26.1 mg of peak 2 with >99.5% purity and 98.98% ee.

**Preparation of 2,5-Dioxopyrrolidin-1-yl 2-(1-(acetyl-^13^C2)-5-(6-methoxypyridin-3-yl)indolin-3-yl)acetate**

Peak 1: To the suspension of 2-(1-(Acetyl-^13^C2)-5-(6-methoxypyridin-3-yl)indolin-3-yl)acetic acid (21.8 mg, 1 Eq, 66.4 μmol, Peak-1) in MeCN (0.5 mL) in a 40mL vial was added Et_3_N (20.8 mg, 28.7 μL, 3.1 Eq, 206 μmol) followed by *N,N’*-Disuccinimidyl Carbonate (29.9 mg, 1.8 Eq, 117 μmol). The mixture was stirred at room temperature for 3h. The reaction mixture was quenched with 0.1M KH_2_PO_4_ aq (10mL), extracted with DCM (4×2 mL). The combined organic layers were dried over Na_2_SO_4_, filtered and removed the solvent under reduced pressure. The residue was purified by silica gel flash chromatography using 0-100% DCM in hexane then 0-40% EtOAc in DCM to obtain 2,5-Dioxopyrrolidin-1-yl 2-(1-(acetyl-^13^C2)-5-(6-methoxypyridin-3-yl)indolin-3-yl)acetate (18.0 mg, 63.7% yield) as a white solid after freeze-drying. LCMS rt 5.38 min, [M+H]^+^ 426.1 expected, 426.2 observed.

Peak 2: 2-(1-(acetyl-13C2)-5-(6-methoxypyridin-3-yl)indolin-3-yl)acetic acid (26.1 mg, 0.0795 mmol), N,N’-Disuccinimidyl Carbonate (22.4 mg, 0.0875 mmol), and TEA (34.3 μL, 0.246 mmol) were dissolved in 0.5 mL of CH₃CN. The reaction was stirred for 1 hour at room temperature. LCMS indicated the consumption of the starting material. The reaction mixture was then diluted with 10 mL of water and 1 mL of saturated NH_4_Cl solution, followed by extraction with DCM (3 × 20 mL). The combined organic layers were washed brine solution (20 mL) and dried over Na_2_SO_4_. The solvent was removed under reduced pressure to give crude product. The crude residue was purified by column chromatography (DCM/EtOAc, 5:5) to provide **T2-peak2**. The product was dissolved in 5 mL of acetonitrile, lyophilized to give **T2-peak2** (0.09 g, 27%) as off -white solid. **^1^H NMR** (300 MHz, CDCl_3_) δ 8.35 (s, 1H) 8.28 (d, *J* = 8.4 Hz, 1H), 7.77 (dd, *J* = 8.4 Hz, 1H), 7.41 (d, *J* = 8.7 Hz, 1H), 7.37 (s, 1H), 6.81 (d, *J* = 8.7 Hz, 1H), 4. 37 (t, *J* = 9.3 Hz, 1H), 3.98 (s, 3H) 3.97-3.88 (m, 2H), 3.2 – 3.13 (m, 1H), 2.94 – 2.8 (m, 5H), 2.25 (bd, *J* = 128.7 Hz, 3 H); LCMS rt 5.29 min, [M+H]^+^: expected 426.1, observed 426.2

**T4:**

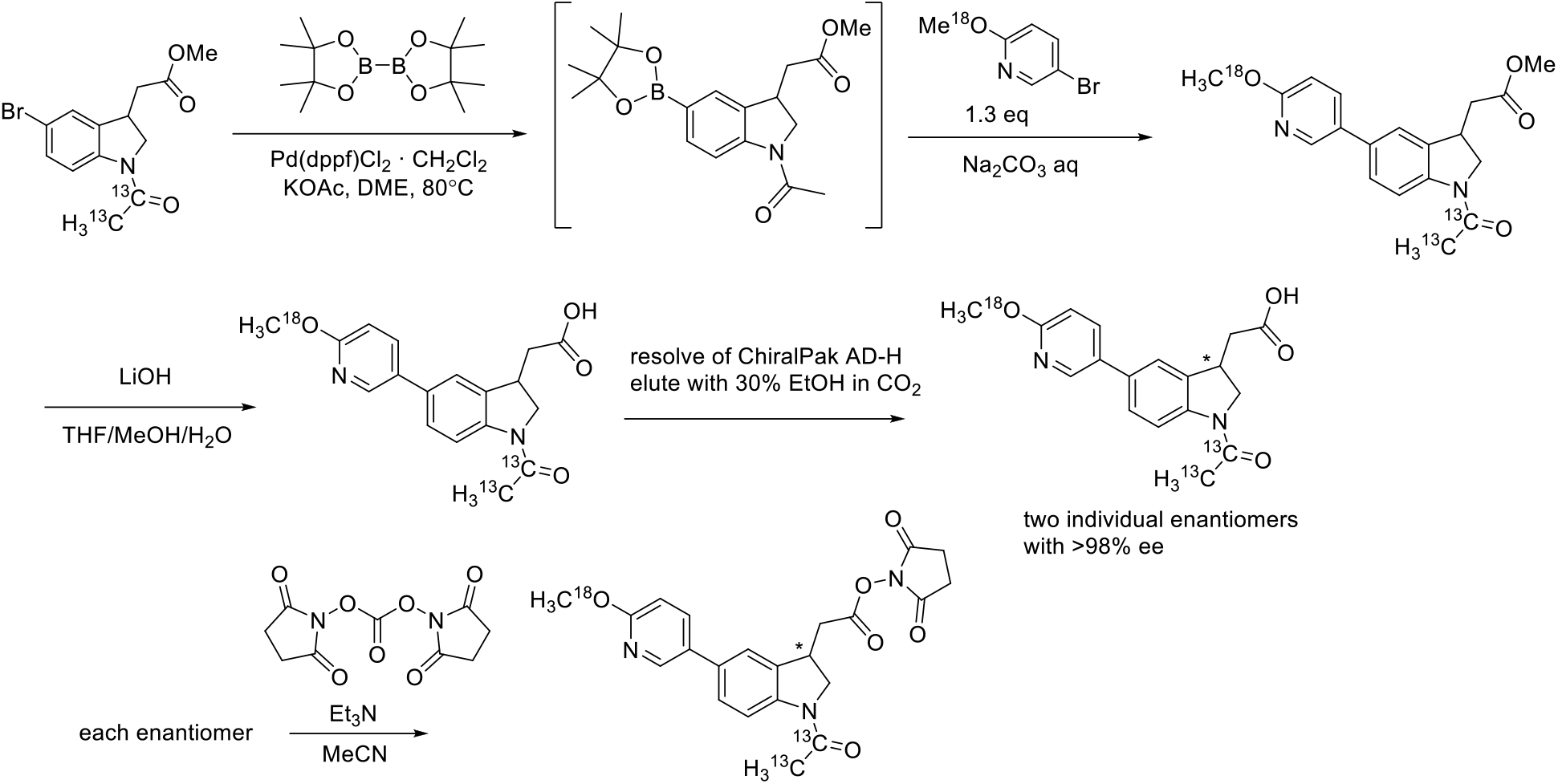

Preparation of **5-bromo-2-(methoxy-^18^O)pyridine**

To a solution of methanol-^18^O (578 mg, 690 μL, 1.20 Eq, 17.0 mmol) in anhydrous THF (20 mL) under nitrogen with vigorous stirring was added the NaH (852.3 mg, 60% Wt, 1.5 Eq, 21.308 mmol) at 0 °C. After stirring at RT for 15 min, 5-bromo-2-fluoropyridine (2500.0 mg, 1 Eq, 14.205 mmol) was added dropwise. The resulting mixture was stirred at RT for 4h, then poured into saturated aq. NH_4_Cl and extracted with EtOAc (5×10 mL). The combined organic layers were dried over Na_2_SO_4_, filtered, and concentrated under reduced pressure to obtain a crude product, which was purified by silica gel flash chromatography by using 0-3% DCM in Hexanes as the eluent to afford 5-bromo-2-(methoxy-^18^O)pyridine (2405.1 mg, 89.1 % yield) as a colorless oil. **^1^H NMR** (300 MHz, CDCl_3_-*d*): δ 8.20 (d, *J* = 2.7 Hz, 1H), 7.63 (dd, *J* = 8.7 Hz, 2.7 Hz, 1.5 Hz, 1H), 6.66 (d, *J* = 9.0 Hz, 1H), 3.90 (S, 3H). **^13^C NMR** (75 MHz, CDCl_3_-*d*): δ 163.25, 147.76, 141.22, 112.78, 111.87, 53.85.

**Preparation of Methyl 2-(1-(acetyl-^13^C2)-5-(6-(methoxy-^18^O)pyridin-3-yl)indolin-3-yl)acetate**

To the solution of methyl 2-(1-(acetyl-^13^C2)-5-bromoindolin-3-yl)acetate (150.0 mg, 1 Eq, 477.5 μmol) in DME (3.6 mL) was added KOAc (93.72 mg, 2 Eq, 955.0 μmol) and Bis(pinacolato)diborane (144.5 mg, 1.2 Eq, 573.0 μmol), degassed and recharged with Argon for 3 times, then added Pd(dppf)Cl_2_·CH_2_Cl_2_ (38.99 mg, 0.1 Eq, 47.75 μmol). The mixture was heated at 80 °C under Argon atmosphere for 90min. After cooling to RT, a solution of Na_2_CO_3_ (101.2 mg, 2 Eq, 955.1 μmol) in Water (1.8 mL) was added to the reaction mixture, followed by the solution of 5-bromo-2-methoxypyridine (120.0 mg, 1.3 Eq, 631.5 μmol) in DME (1.8 mL). After heating at 80 °C under Argon atmosphere for 1h, the solvent was removed under reduced pressure. The residue was purified by silica gel flash chromatography by using 0-40% EtOAc in hexane as the eluent to obtain the desired product (139 mg, 84.5% yield) as a yellow foamy solid. LCMS rt 5.41 min, [M+H]^+^ 346.2 expected, 346.2 observed.

**Preparation of 2-(1-(Acetyl-13C2)-5-(6-(methoxy-^18^O)pyridin-3-yl)indolin-3-yl)acetic acid**

To the solution of Methyl 2-(1-(acetyl-^13^C2)-5-(6-(methoxy-^18^O)pyridin-3-yl)indolin-3-yl)acetate (139 mg, 1 Eq, 404 μmol) in Methanol (0.5 mL), THF (1 mL), and Water (0.5 mL) was added LiOH (38.7 mg, 4 Eq, 1.61mmol). The reaction mixture was stirred at RT for 5h, then acidified by 3M HCl aq to adjust the pH=4∼5 to precipitate the product. The suspension was diluted with 2 mL of water, sonicated for 1min, centrifuged, and removed the clear liquid on the upper layer by pipet. Repeated the water wash twice (2×2mL), then used MeCN (3×2mL) instead of water, repeated 3 times. The solid was suspended in 1:1 MeCN/water, freeze-dried to obtain 2-(1-(Acetyl-^13^C2)-5-(6-(methoxy-^18^O)pyridin-3-yl)indolin-3-yl)acetic acid (111 mg, 83.5% yield) as an off-white solid. LCMS rt 4.77 min, [M+H]^+^ 331.1 expected, 331.1 observed

**Resolution:** ChrialPak AD-H 21×250 mm, with 30% ethanol in CO_2_ and a flow rate of 70 mL/min. Sample dissolved at 2 mg/mL in 50:50 dichloromethane:methanol and loaded at 2 mL per injection to afford 33.4 mg of peak 1, >99.5% purity and ee, 35.3 mg of peak 2 with >99.5% purity and 98.72% ee.

**Preparation of 2,5-Dioxopyrrolidin-1-yl 2-(1-(acetyl-^13^C2)-5-(6-(methoxy-^18^O)pyridin-3-yl) indolin-3-yl)acetate**

Peak 1: To the suspension of 2-(1-(acetyl-^13^C2)-5-(6-(methoxy-^18^O)pyridin-3-yl)indolin-3-yl)acetic acid (26.3 mg, 1 Eq, 79.6 μmol, Peak-1) in MeCN (0.5 mL) in a 40mL vial was added Et_3_N (22 mg, 30 μL, 2.7 Eq, 215 μmol). The solvent was removed under reduced pressure and dried in high vacuum overnight. The solid was suspended in MeCN (0.5 mL) in a 20mL vial, and Et_3_N (16.9 mg, 23.3 μL, 2.1 Eq, 167 μmol) was added, followed by *N,N’*-Disuccinimidyl Carbonate (29.2 mg, 1.4 Eq, 114 μmol). The mixture was stirred at room temperature for 1h. The reaction mixture was quenched with 0.1M KH_2_PO_4_ aq (10mL), extracted with DCM (4×2 mL). The combined organic layers were dried over Na_2_SO_4_, filtered and removed the solvent under reduced pressure. The residue was purified by silica gel flash chromatography using 0-100% DCM in hexane then 0-40% EtOAc in DCM to obtain 2,5-Dioxopyrrolidin-1-yl 2-(1-(acetyl-^13^C2)-5-(6-(methoxy-^18^O)pyridin-3-yl) indolin-3-yl)acetate (20.9 mg, 61.4% yield) as a white solid after freeze-drying. LCMS rt 5.29 min, [M+H]^+^ 428.2 expected, 428.1 observed.

Peak 2: 2-(1-(acetyl-13C2)-5-(6-(methoxy-^18^O)pyridin-3-yl)indolin-3-yl)acetic acid (34mg, 0.10 mmol), N,N’-Disuccinimidyl Carbonate (29 mg, 0.11 mmol), and TEA (44 μL, 0.32 mmol) were dissolved in 0.5 mL of CH₃CN. The reaction was stirred for 1 hour at room temperature. LCMS indicated the consumption of the starting material. The reaction mixture was then diluted with 10 mL of DCM and 10 mL of 0.1 M KH_2_PO₄ solution, followed by extraction with DCM (3 × 20 mL). The combined organic layers were washed brine solution (20 mL) and dried over Na_2_SO_4_. The solvent was removed under reduced pressure to give crude product. The crude residue was purified by column chromatography (DCM/EtOAc, 5:5) to provide **T4-peak2**. The product was dissolved in 5 mL of acetonitrile, lyophilized, and give **T4-peak2** (0.035 g, 80%) as off -white solid. **^1^H NMR** (300 MHz, CDCl_3_) δ 8.35 (s, 1H) 8.28 (d, *J* = 8.4 Hz, 1H), 7.77 (dd, *J* = 8.4 Hz, 1H), 7.41 (d, *J* = 8.7 Hz, 1H), 7.38 (s, 1H), 6.81 (d, *J* = 8.7 Hz, 1H), 4. 37 (t, *J* = 8.7 Hz, 1H), 4.05-3.88 (m, 5H), 3.2 – 3.13 (m, 1H), 2.94 – 2.8 (m, 5H), 2.25 (bd, *J* = 128.4 Hz, 3 H); LCMS 5.57 min, [M+H]^+^: 428.2 expected, 428.1 observed

**T6:**

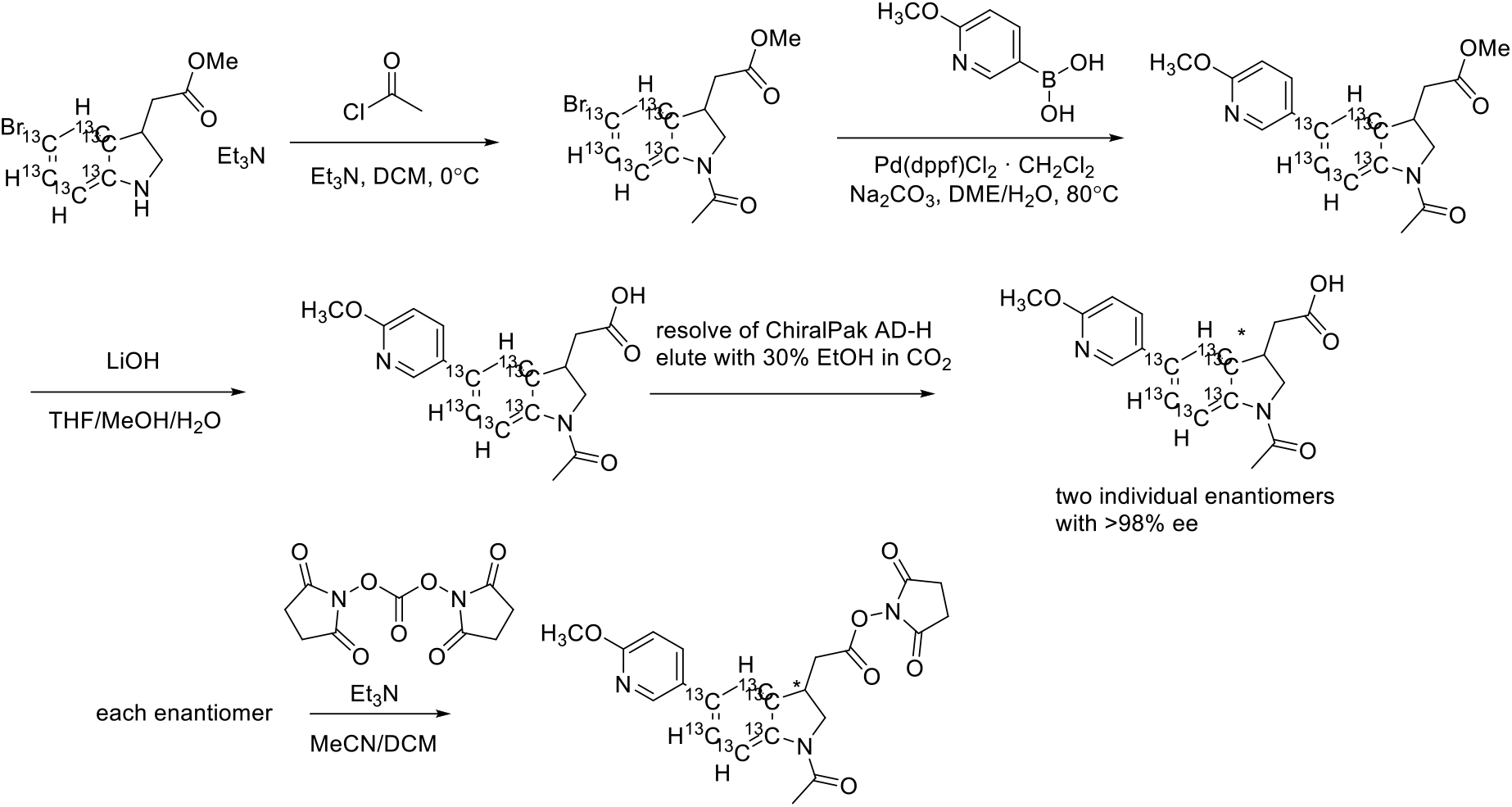

Preparation of **Methyl 2-(1-acetyl-5-bromoindolin-3-yl-3a,4,5,6,7,7a-^13^C6)acetate**

To a solution of methyl 2-(5-bromoindolin-3-yl-3a,4,5,6,7,7a-^13^C6)acetate (200 mg, 1 Eq, 724 μmol) in DCM (5 mL) in an ice bath was added triethylamine (147 mg, 202 μL, 2 Eq, 1.45 mmol) followed by Acetyl chloride (62.6 mg, 56.3 μL, 1.1 Eq, 797 μmol). After stirring for 2h, the reaction mixture was quenched with sat. NaHCO_3_ aq (2 mL). The aqueous layer was separated and washed with DCM (3’2 mL). The combined organic layers were dried over Na_2_SO_4_, filtered and removed the solvent under reduced pressure. The residue was purified by silica gel flash chromatography using 0-40% DCM in hexane then 0-40% EtOAc in Hexane to obtain methyl 2-(1-acetyl-5-bromoindolin-3-yl-3a,4,5,6,7,7a-^13^C6)acetate (95.6 mg, 41.5 % yield) as an off-white solid. LCMS rt 5.57 min, 318.0/320.0 expected, 318.2/320.2 observed.

Preparation of **Methyl 2-(1-acetyl-5-(6-methoxypyridin-3-yl)indolin-3-yl-3a,4,5,6,7,7a-^13^C6)acetate**

To the suspension of methyl 2-(1-acetyl-5-bromoindolin-3-yl-3a,4,5,6,7,7a-^13^C6)acetate (142.6 mg, 1 Eq, 448.3 μmol), 2-Methoxy-5-pyridinylboronic Acid (82.27 mg, 1.2 Eq, 537.9 μmol) and Na_2_CO_3_ (95.02 mg, 2 Eq, 896.5 μmol) in DME (1.2 mL) and Water (0.4 mL) was added [1,1’-Bis(diphenylphosphino)ferrocene]dichloropalladium (II) Complex with Dichloromethane (18.30 mg, 0.05 Eq, 22.41 μmol), then degassed and refilled by Argon for 3 times. The mixture was heated at 80 °C for 1h, then removed the solvent under reduced pressure. The residue was purified by silica gel flash chromatography using 0-40%EtOAc in Hexane to obtain Methyl 2-(1-acetyl-5-(6-methoxypyridin-3-yl)indolin-3-yl-3a,4,5,6,7,7a-^13^C6)acetate (151 mg, 97.27% yield) as a very light yellow foamy semi-solid. LCMS rt 5.57 min, 346.2 expected, 346.1 observed.

Preparation of **2-(1-Acetyl-5-(6-methoxypyridin-3-yl)indolin-3-yl-3a,4,5,6,7,7a-^13^C6)acetic acid**

To the solution of Methyl 2-(1-acetyl-5-(6-methoxypyridin-3-yl)indolin-3-yl-3a,4,5,6,7,7a-^13^C6)acetate (142.0 mg, 1 Eq, 410.0 μmol) in Methanol (0.5 mL), THF (1 mL), and Water (0.5 mL) was added LiOH (42.8 mg, 4.4 Eq, 1.78mmol). The reaction mixture was stirred at RT for 3h, then acidified by 3M HCl aq to adjust the pH=4∼5 to precipitate the product. The suspension was diluted with 4 mL of water, sonicated for 1min, centrifuged, and removed the clear liquid on the upper layer by pipet. Repeated the water wash twice (2×4mL), then used MeCN (3×2mL) instead of water, repeated 3 times. The solid was suspended in 1:1 MeCN/water, freeze-dried to obtain 2-(1-Acetyl-5-(6-methoxypyridin-3-yl)indolin-3-yl-3a,4,5,6,7,7a-^13^C6)acetic acid (116 mg, 85.1% yield) as a white solid. LCMS rt 4.85 min, [M+H]^+^ 333.1 expected, 333.1 observed.

**Resolution:** ChrialPak AD-H 21×250 mm, with 30% ethanol in CO_2_ and a flow rate of 70 mL/min. Sample dissolved at 2 mg/mL in 50:50 dichloromethane:methanol and loaded at 2 mL per injection to afford 27 mg of peak 1, >99.5% purity and ee, 28.9 mg of peak 2 with >99.5% purity and ee.

Preparation of **2,5-Dioxopyrrolidin-1-yl 2-(1-acetyl-5-(6-methoxypyridin-3-yl)indolin-3-yl-3a,4,5,6,7,7a-^13^C6)acetate**

Peak 1: To the suspension of 2-(1-Acetyl-5-(6-methoxypyridin-3-yl)indolin-3-yl-3a,4,5,6,7,7a-^13^C6)acetic acid (27 mg, 1 Eq, 81 μmol, Peak-1) in MeCN (0.5 mL) in a 20mL vial was added Et_3_N (22 mg, 30 μL, 2.6 Eq, 215 μmol). The solvent was removed under reduced pressure and dried in high vacuum for 30 min. The solid was suspended in MeCN (0.5 mL) in a 20mL vial, and Et_3_N (22 mg, 30 μL, 2.6 Eq, 215 μmol) was added, followed by *N,N’*-Disuccinimidyl Carbonate (29.7 mg, 1.4 Eq, 116 μmol). The mixture was stirred at room temperature for 1h. The reaction mixture was quenched with 0.1M KH_2_PO_4_ aq (10mL), extracted with DCM (4×2 mL). The combined organic layers were dried over Na_2_SO_4_, filtered and removed the solvent under reduced pressure. The residue was purified by silica gel flash chromatography using 0-100% DCM in hexane then 0-40% EtOAc in DCM to obtain 2,5-Dioxopyrrolidin-1-yl 2-(1-acetyl-5-(6-methoxypyridin-3-yl)indolin-3-yl-3a,4,5,6,7,7a-^13^C6)acetate (19.7 mg, 56.5% yield) as a white solid after freeze-drying. LCMS rt 5.48 min, [M+H]^+^ expected 430.2 observed 430.3

Peak 2: 2-(1-acetyl-5-(6-methoxypyridin-3-yl)indolin-3-yl-3a,4,5,6,7,7a-13C6)acetic acid (28mg, 00.084 mmol), N,N’-Disuccinimidyl Carbonate (24 mg, 0.092 mmol), and TEA (36 μL, 0.26 mmol) were dissolved in 0.5 mL of CH₃CN. The reaction was stirred for 1 hour at room temperature. LCMS indicated the consumption of the starting material. The reaction mixture was then diluted with 10 mL of DCM and 10 mL of 0.1 M KH_2_PO₄ solution, followed by extraction with DCM (3 × 20 mL). The combined organic layers were washed brine solution (20 mL) and dried over Na_2_SO_4_. The solvent was removed under reduced pressure to give crude product. The crude residue was purified by column chromatography (DCM/EtOAc, 5:5) to provide **T6-peak2**. The product was dissolved in 5 mL of acetonitrile and lyophilized to give **T6-peak2** (0.021 g, 58%) as off -white solid. **^1^H NMR** (300 MHz, CDCl_3_) δ 8.35 (s, 1H), 8.25 (bd, *J* = 167.4 Hz, 1 H), 7.79 – 7.63 (m, 2H), 7.17 – 7.10 (m, 1H), 6.81 (dd, *J* = 8.4 Hz, 1H), 4. 37 (t, *J* = 8.4 Hz, 1H), 4.0-3.88 (m, 5H), 3.2 – 3.13 (m, 1H), 2.94 – 2.8 (m, 5H), 2.28 (s, 3H); LCMS rt 5.55 min [M+H]^+^: expected 430.2 observed 430.3

**T8:**

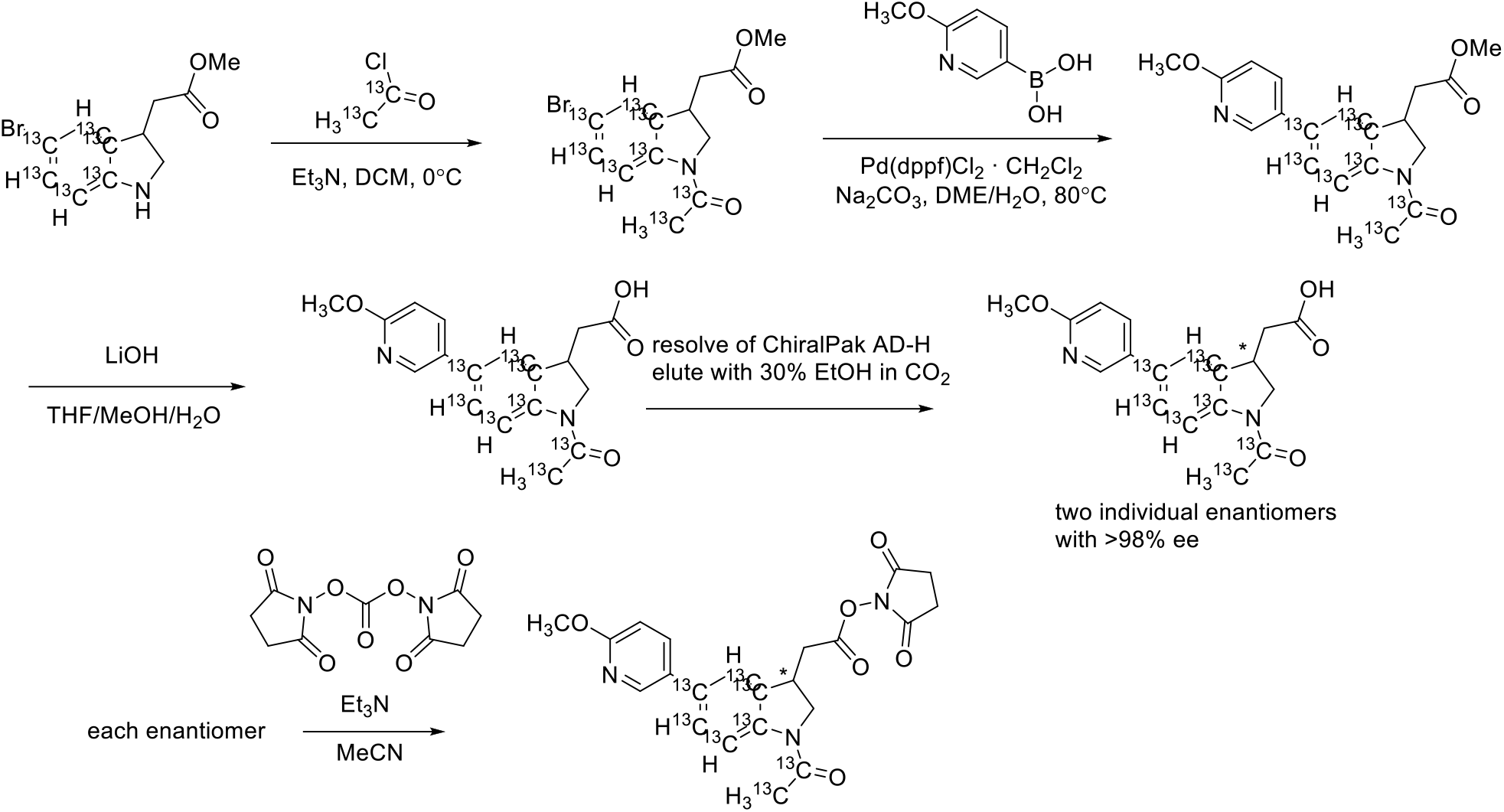

**Methyl 2-(1-(acetyl-^13^C2)-5-bromoindolin-3-yl-3a,4,5,6,7,7a-^13^C6)acetate** was prepared according to the protocol used for methyl 2-(1-acetyl-5-bromoindolin-3-yl-3a,4,5,6,7,7a-^13^C6)acetate. 271.5 mg (67%) LCMS rt 5.82 min, [M+H]^+^ expected 320.0/322.0 observed 320.3/322.3

**Methyl 2-(1-(acetyl-^13^C2)-5-(6-methoxypyridin-3-yl)indolin-3-yl-3a,4,5,6,7,7a-^13^C6)acetate** was prepared according to the protocol used for methyl 2-(1-acetyl-5-(6-methoxypyridin-3-yl)indolin-3-yl-3a,4,5,6,7,7a-^13^C6)acetate. 99.0 mg (76%) LCMS rt 5.71 min, [M+H]^+^ expected 349.2 observed 349.1

**2-(1-(Acetyl-^13^C2)-5-(6-methoxypyridin-3-yl)indolin-3-yl-3a,4,5,6,7,7a-^13^C6)acetic acid** was prepared according to the protocol used for 2-(1-Acetyl-5-(6-methoxypyridin-3-yl)indolin-3-yl-3a,4,5,6,7,7a-^13^C6)acetic acid 66.0 mg (76%) LCMS rt 4.87 min, [M+H]^+^ expected 335.2 observed 335.0

**Resolution:** ChrialPak AD-H 21×250 mm, with 30% ethanol in CO_2_ and a flow rate of 70 mL/min. Sample dissolved at 2 mg/mL in 50:50 dichloromethane:methanol and loaded at 2 mL per injection to afford 23.6 mg of peak 1, >99.5% purity and ee, 24.7 mg of peak 2 with >99.5% purity and ee.

Peak1: 2,5-Dioxopyrrolidin-1-yl 2-(1-(acetyl-^13^C2)-5-(6-methoxypyridin-3-yl)indolin-3-yl-3a,4,5,6,7,7a-^13^C6)acetate was prepared according to the protocol used for 2,5-Dioxopyrrolidin-1-yl 2-(1-acetyl-5-(6-methoxypyridin-3-yl)indolin-3-yl-3a,4,5,6,7,7a-13C6)acetate. 21.2 mg (70%) LCMS rt min, [M+H]^+^ 5.59 expected 432.2 observed 432.3

Peak 2: 2-(1-(acetyl-13C2)-5-(6-methoxypyridin-3-yl)indolin-3-yl-3a,4,5,6,7,7a-13C6)acetic acid (24mg, 0.072 mmol), N,N’-Disuccinimidyl Carbonate (20 mg, 0.079 mmol), and TEA (31 μL, 0.22 mmol) were dissolved in 0.5 mL of CH₃CN. The reaction was stirred for 1 hour at room temperature. LCMS indicated the consumption of the starting material. The reaction mixture was then diluted with 10 mL of DCM and 10 mL of saturated NaHCO_3_ solution, followed by extraction with DCM (3 × 20 mL). The combined organic layers were washed brine solution (20 mL) and dried over Na_2_SO_4_. The solvent was removed under reduced pressure to give crude product. The crude residue was purified by column chromatography (DCM/EtOAc, 5:5) to provide **T8-peak2**. The product was dissolved in 5 mL of acetonitrile and lyophilized to give **T8-peak2** (0.010 g, 32%) as off -white solid. **^1^H NMR** (300 MHz, CDCl_3_) δ 8.35 (s, 1H), 8.25 (bd, *J* = 168.6 Hz, 1 H), 7.79 – 7.60 (m, 2H), 7.17 – 7.10 (m, 1H), 6.81 (d, *J* = 9.0 Hz, 1H), 4. 37 (t, *J* = 9.6 Hz, 1H), 3.98 (s, 3H) 3.97-3.88 (m, 2H), 3.2 – 3.13 (m, 1H), 2.94 – 2.8 (m, 5H), 2.25 (bd, *J* = 128.7 Hz, 3 H); LCMS rt 5.26 min, [M+H]^+^: expected 432.2 observed 432.1

**T10:**

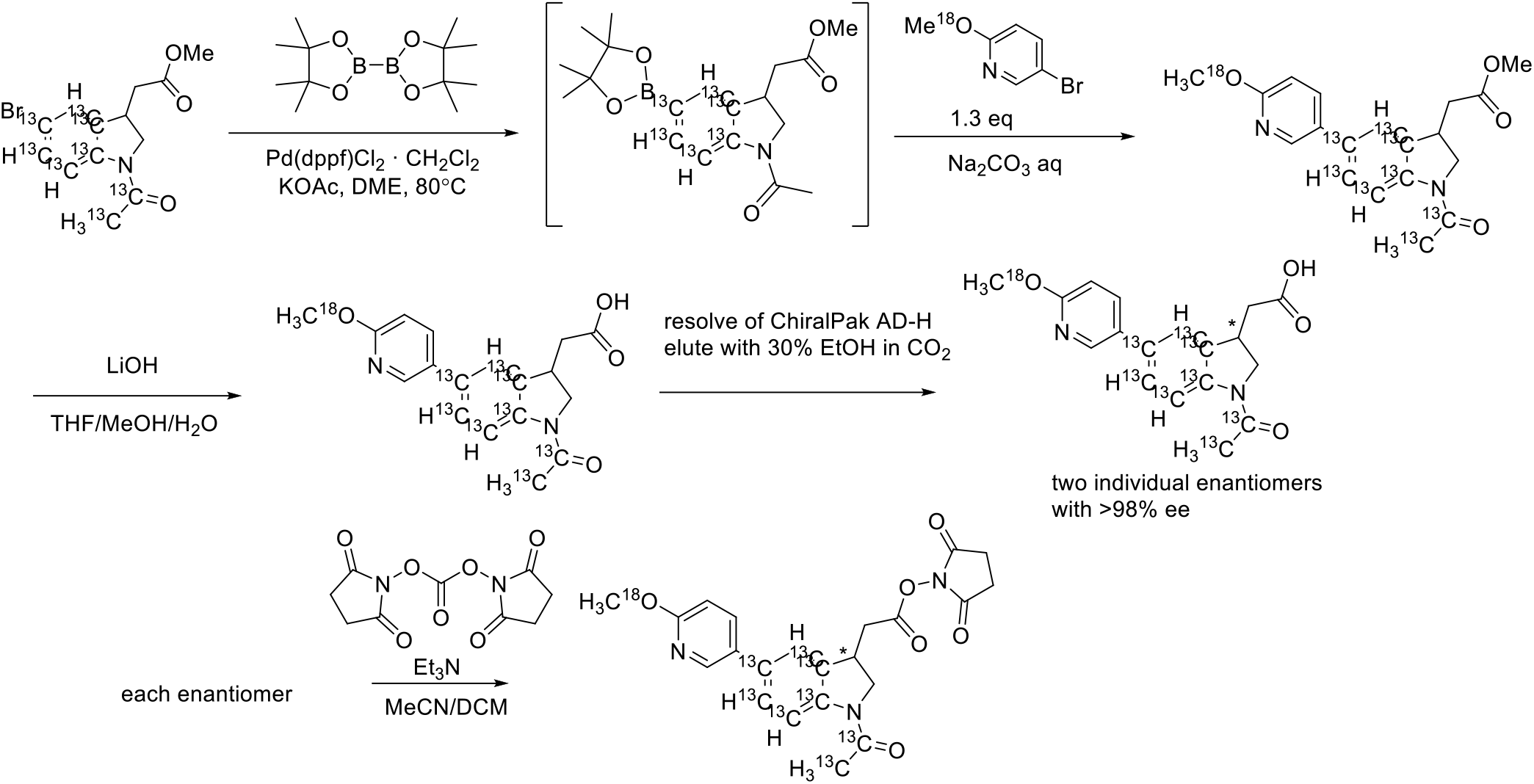

**Methyl 2-(1-(acetyl-^13^C2)-5-(6-(methoxy-^18^O)pyridin-3-yl)indolin-3-yl-3a,4,5,6,7,7a-^13^C6)acetate** was prepared according to the protocol used for methyl 2-(1-(acetyl-^13^C2)-5-(6-(methoxy-^18^O)pyridin-3-yl)indolin-3-yl)acetate. 120 mg (75%) LCMS rt 5.38 min, [M+H]^+^ expected 351.1 observed 351.1

**2-(1-(Acetyl-^13^C2)-5-(6-(methoxy-^18^O)pyridin-3-yl)indolin-3-yl-3a,4,5,6,7,7a-^13^C6)acetic acid** was prepared according to the protocol used for 2-(1-Acetyl-5-(6-methoxypyridin-3-yl)indolin-3-yl-3a,4,5,6,7,7a-^13^C6)acetic acid. 76 mg (79%) LCMS rt 4.95 min, [M+H]^+^ expected 337.2 observed 337.1

**Resolution:** ChrialPak AD-H 21×250 mm, with 30% ethanol in CO_2_ and a flow rate of 70 mL/min. Sample dissolved at 2 mg/mL in 50:50 dichloromethane:methanol and loaded at 2 mL per injection to afford 32 mg of peak 1, >99.5% purity and ee, 33.9 mg of peak 2 with >99.5% purity and ee.

Peak 1: 2,5-Dioxopyrrolidin-1-yl 2-(1-(acetyl-^13^C2)-5-(6-(methoxy-^18^O)pyridin-3-yl)indolin-3-yl-3a,4,5,6,7,7a-^13^C6)acetate was prepared according to the protocol used for 2,5-Dioxopyrrolidin-1-yl 2-(1-(acetyl-^13^C2)-5-(6-methoxypyridin-3-yl)indolin-3-yl-3a,4,5,6,7,7a-^13^C6)acetate. 31.8 mg (77%) LCMS rt 5.55 min, [M+H]^+^ expected 434.2 observed 434.2

Peak 2: 2-(1-(acetyl-13C2)-5-(6-(methoxy-^18^O)pyridin-3-yl)indolin-3-yl-3a,4,5,6,7,7a-13C6)acetic acid (33 mg, 0.072 mmol), N,N’-Disuccinimidyl Carbonate (28 mg, 0.11 mmol), and TEA (42 μL, 0.30 mmol) were dissolved in 0.5 mL of CH₃CN. The reaction was stirred for 1 hour at room temperature. LCMS indicated the consumption of the starting material. The reaction mixture was then diluted with 10 mL of DCM and 10 mL of 0.1 M KH_2_PO₄ solution, followed by extraction with DCM (3 × 20 mL). The combined organic layers were washed brine solution (20 mL) and dried over Na_2_SO_4_. The solvent was removed under reduced pressure to give crude product. The crude residue was purified by column chromatography (DCM/EtOAc, 5:5) to provide **T10-peak2**. The product was dissolved in 5 mL of acetonitrile and lyophilized to give **T10-peak2** (0.037 g, 87%) as off -white solid. **^1^H NMR** (300 MHz, CDCl_3_) δ 8.35 (s, 1H), 8.25 (bd, *J* = 172.2 Hz, 1 H), 7.79 – 7.63 (m, 2H), 7.17 – 7.10 (m, 1H), 6.81 (dd, *J* = 8.4 Hz, 1H), 4. 37 (t, *J* = 8.4 Hz, 1H), 4.0-3.88 (m, 5H), 3.2 – 3.13 (m, 1H), 2.94 – 2.8 (m, 5H), 2.25 (bd, *J* = 128.4 Hz, 3 H); LCMS rt 5.50 min, [M+H]^+^: expected 434.2 observed 434.3

**T12:**

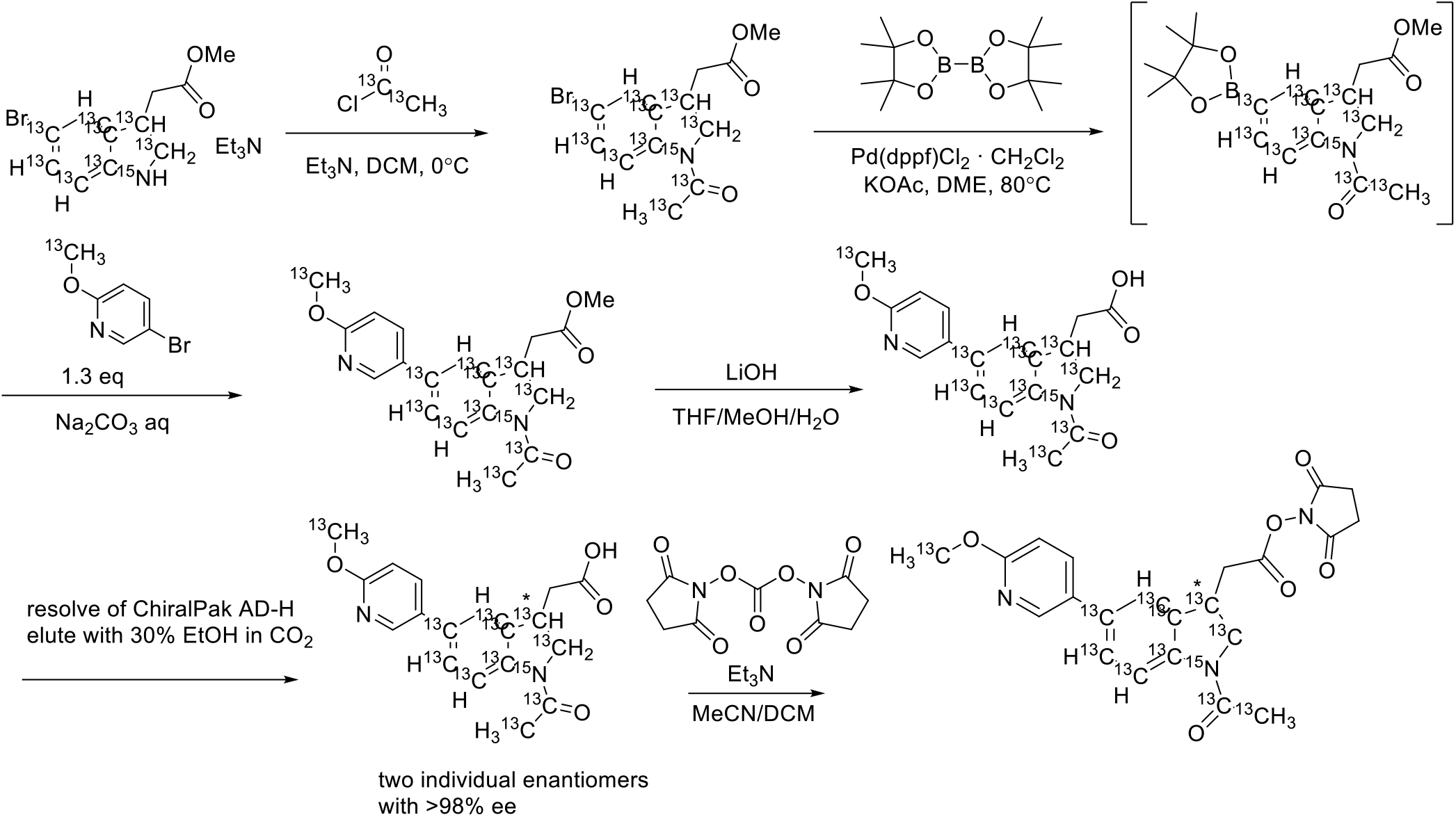

**Methyl 2-(1-(acetyl-^13^C2)-5-bromoindolin-3-yl-^13^C8-^15^N)acetate** was prepared according to the protocol used for methyl 2-(1-acetyl-5-bromoindolin-3-yl-3a,4,5,6,7,7a-^13^C6)acetate 331.8 mg (92%) LCMS rt 5.79 min, [M+H]^+^ expected 323.0/325.0. observed 323.3/325.3

**Methyl 2-(1-(acetyl-^13^C2)-5-(6-(methoxy-^13^C)pyridin-3-yl)indolin-3-yl-^13^C8-^15^N)acetate** was prepared according to the protocol used for methyl 2-(1-(acetyl-^13^C2)-5-(6-(methoxy-^18^O)pyridin-3-yl)indolin-3-yl)acetate. 93 mg (58%) LCMS rt 5.60 min, [M+H]^+^ expected 353.2 observed 353.3

**2-(1-(Acetyl-^13^C2)-5-(6-(methoxy-^13^C)pyridin-3-yl)indolin-3-yl-^13^C8-^15^N)acetic acid** was prepared according to the protocol used for 2-(1-acetyl-5-(6-methoxypyridin-3-yl)indolin-3-yl-3a,4,5,6,7,7a-^13^C6)acetic acid 62.2 mg (75%) LCMS rt 4.89 min, [M+H]^+^ expected 339.2 observed 339.2

**Resolution:** ChrialPak AD-H 21×250 mm, with 30% ethanol in CO_2_ and a flow rate of 70 mL/min. Sample dissolved at 2 mg/mL in 50:50 dichloromethane:methanol and loaded at 2 mL per injection to afford 25.2 mg of peak 1, >99.5% purity and ee, 27.3 mg of peak 2 with >99.5% purity and 98.24% ee.

Peak 1: 2,5-dioxopyrrolidin-1-yl 2-(1-(acetyl-^13^C2)-5-(6-(methoxy-^13^C)pyridin-3-yl)indolin-3-yl-^13^C8-^15^N)acetate was prepared according to the protocol used for 2,5-Dioxopyrrolidin-1-yl 2-(1-(acetyl-^13^C2)-5-(6-methoxypyridin-3-yl)indolin-3-yl-3a,4,5,6,7,7a-^13^C6)acetate 26.3 mg (81%) LCMS rt 5.51 min, [M+H]^+^ expected 436.2 observed 436.2 Peak 2: 2-(1-(acetyl-13C2)-5-(6-(methoxy-^13^C)pyridin-3-yl)indolin-3-yl-^13^C8-^15^N)acetic acid (27 mg, 0.080 mmol), N,N’-Disuccinimidyl Carbonate (23 mg, 0.088 mmol), and TEA (35 μL, 0.25 mmol) were dissolved in 0.5 mL of CH₃CN. The reaction was stirred for 1 hour at room temperature. LCMS indicated the consumption of the starting material. The reaction mixture was then diluted with 10 mL of DCM and 10 mL of 0.1 M KH_2_PO₄ solution, followed by extraction with DCM (3 × 20 mL). The combined organic layers were washed brine solution (20 mL) and dried over Na_2_SO_4_. The solvent was removed under reduced pressure to give crude product. The crude residue was purified by column chromatography (DCM/EtOAc, 5:5) to provide **T12-peak2**. The product was dissolved in 5 mL of acetonitrile and lyophilized to give **T12-peak2** (0.0253 g, 73 %) as off -white solid. **^1^H NMR** (300 MHz, CDCl_3_) δ 8.35 (s, 1H), 8.25 (bd, *J* = 169.5 Hz, 1 H), 7.79 – 7.63 (m, 2H), 7.17 – 7.10 (m, 1H), 6.81 (dd, *J* = 8.4 Hz, 1H), 4. 37 (t, *J* = 9.3 Hz, 1H), 4.29-3.90 (m, 5H), 3.2 – 3.13 (m, 1H), 2.94 – 2.8 (m, 5H), 2.25 (bd, *J* = 128.1 Hz, 3 H); LCMS rt 5.44 min, [M+H]^+^: expected 436.2, observed 436.1

**T14:**

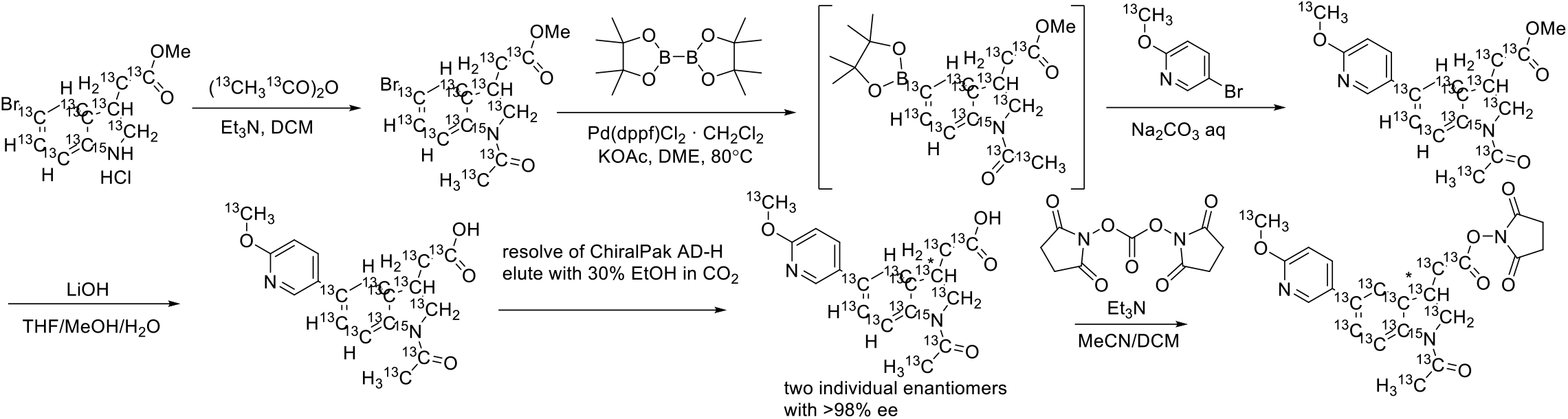

**methyl 2-(1-(acetyl-^13^C2)-5-(6-(methoxy-^13^C)pyridin-3-yl)indolin-3-yl-^13^C8-^15^N)acetate-^13^C2** was prepared according to the protocol used for methyl 2-(1-(acetyl-^13^C2)-5-(6-(methoxy-^18^O)pyridin-3-yl)indolin-3-yl)acetate 97.1 mg (89%) LCMS RT: 5.38 min, [M+H]^+^ 355.2 expected, observed 355.1

**2-(1-(acetyl-^13^C2)-5-(6-(methoxy-^13^C)pyridin-3-yl)indolin-3-yl-^13^C8-^15^N)acetic-^13^C2 acid** was prepared according to the protocol used for 2-(1-acetyl-5-(6-methoxypyridin-3-yl)indolin-3-yl-3a,4,5,6,7,7a-^13^C6)acetic acid 79.1 mg LCMS rt 4.84 min, [M+H]^+^: expected 341.2, observed 341.0

**Resolution:** ChrialPak AD-H 21×250 mm, with 30% ethanol in CO_2_ and a flow rate of 70 mL/min. Sample dissolved at 2 mg/mL in 50:50 dichloromethane:methanol and loaded at 2 mL per injection to afford 36.5 mg of peak 1, >99.5% purity and ee, 33.3 mg of peak 2 with >99.5% purity and ee.

Peak 1: prepared according to the protocol used for 2,5-Dioxopyrrolidin-1-yl 2-(1-(acetyl-^13^C2)-5-(6-methoxypyridin-3-yl)indolin-3-yl-3a,4,5,6,7,7a-^13^C6)acetate 33.3 mg (71%) LCMS rt 5.25 min, [M+H]^+^: expected 438.2, observed 437.8

Peak 2: 2-(1-(acetyl-^13^C2)-5-(6-(methoxy-^13^C)pyridin-3-yl)indolin-3-yl-^13^C8-^15^N)acetic-^13^C2 acid (33 mg, 0.097 mmol), N,N’-Disuccinimidyl Carbonate (27 mg, 011 mmol), and TEA (42 μL, 0.30 mmol) were dissolved in 0.5 mL of CH₃CN. The reaction was stirred for 1 hour at room temperature. LCMS indicated the consumption of the starting material. The reaction mixture was then diluted with 10 mL of DCM and 10 mL of 0.1 M KH_2_PO₄ solution, followed by extraction with DCM (3 × 20 mL). The combined organic layers were washed brine solution (20 mL) and dried over Na_2_SO_4_. The solvent was removed under reduced pressure to give crude product. The crude residue was purified by column chromatography (DCM/EtOAc, 5:5) to provide **T14-peak2**. The product was dissolved in 5 mL of acetonitrile and lyophilized to give **T14-peak2** (0.027 g, 64 %) as off -white solid. **^1^H NMR** (300 MHz, CDCl_3_) δ 8.36 (s, 1H), 8.25 (bd, *J* = 167.1 Hz, 1 H), 7.76 – 7.67 (m, 2H), 7.19 – 7.12 (m, 1H), 6.83 (dd, *J* = 7.8 Hz, 1H), 4. 23 - 4.152 (m, 3H), 3.80-3.70 (m, 2H), 3.3 – 3.17 (m, 1H), 2.90 – 2.70 (m, 5H), 2.25 (bd, *J* = 128.7 Hz, 3 H); LCMS rt 5.60 min [M+H]^+^: expected 438.2, observed 437.7

**T16:**

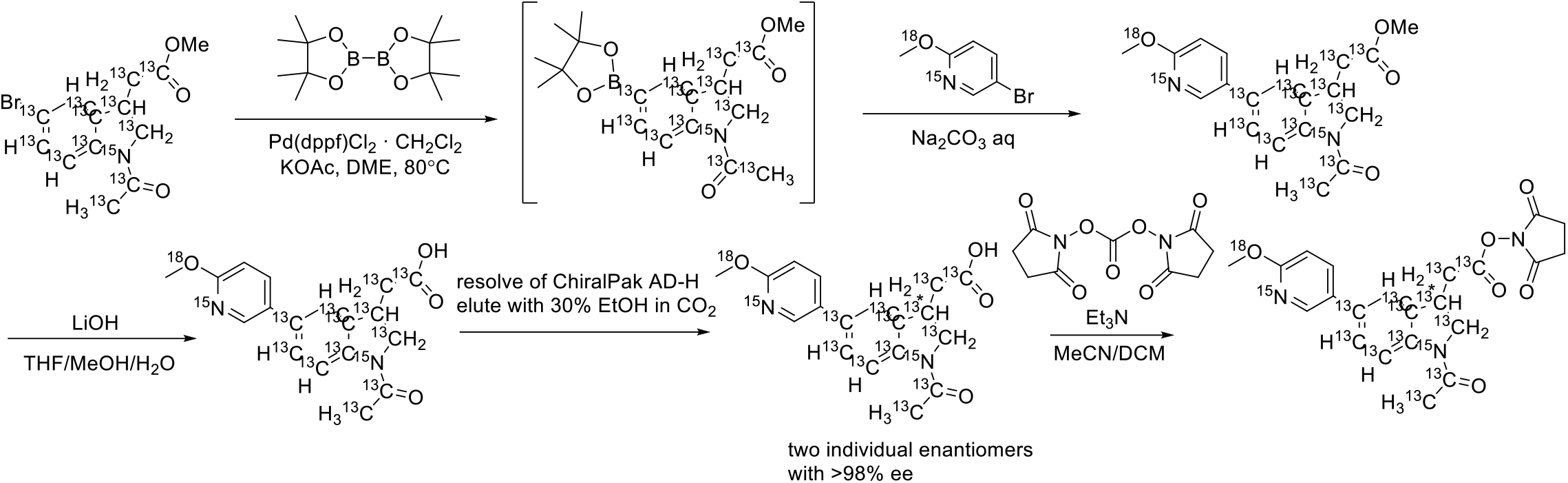

**Methyl 2-(1-(acetyl-^13^C2)-5-(6-(methoxy-^18^O)pyridin-3-yl-1-^15^N)indolin-3-yl-^13^C8-^15^N)acetate-^13^C2** was prepared according to the protocol used for methyl 2-(1-(acetyl-^13^C2)-5-(6-(methoxy-^18^O)pyridin-3-yl)indolin-3-yl)acetate 85.4 mg (78%) [M+H]^+^ 357.2 expected, observed 357.0

**2-(1-(acetyl-^13^C2)-5-(6-(methoxy^18^O)pyridin-3-yl-1-15N)indolin-3-yl-^13^C8-^15^N)acetic-^13^C2 acid** was prepared according to the protocol used for 2-(1-acetyl-5-(6-methoxypyridin-3-yl)indolin-3-yl-3a,4,5,6,7,7a-^13^C6)acetic acid. 72.7 mg (89%) LCMS RT: 5.37min, [M+H]^+^ 343.2 expected, observed 343.2

**Resolution**: ChrialPak AD-H 21×250 mm, with 30% ethanol in CO_2_ and a flow rate of 70 mL/min. Sample dissolved at 2 mg/mL in 50:50 dichloromethane:methanol and loaded at 2 mL per injection to afford 26.5 mg of peak 1, >99.5% purity and ee, 26.8 mg of peak 2 with >99.5% purity and 98.36% ee.

**2,5-Dioxopyrrolidin-1-yl 2-(1-(acetyl-^13^C2)-5-(6-(methoxy-^18^O)pyridin-3-yl-1-^15^N)indolin-3-yl-^13^C8-^15^N)acetate-^13^C2**

Peak 1: prepared according to the protocol used for 2,5-Dioxopyrrolidin-1-yl 2-(1-(acetyl-^13^C2)-5-(6-methoxypyridin-3-yl)indolin-3-yl-3a,4,5,6,7,7a-^13^C6)acetate 26.2 mg (77%) LCMS rt 5.48 min, [M+H]^+^ 440.2 expected, 439.9 observed

Peak 2: 2-(1-(acetyl-^13^C2)-5-(6-(methoxy-^18^O)pyridin-3-yl-1-^15^N)indolin-3-yl-^13^C8-^15^N)acetic-^13^C2 acid (26 mg, 0.076 mmol), N,N’-Disuccinimidyl Carbonate (22 mg, 084 mmol), and TEA (33 μL, 0.24 mmol) were dissolved in 0.5 mL of CH₃CN. The reaction was stirred for 1 hour at room temperature. LCMS indicated the consumption of the starting material. The reaction mixture was then diluted with 10 mL of DCM and 10 mL of 0.1 M KH_2_PO₄ solution, followed by extraction with DCM (3 × 20 mL). The combined organic layers were washed brine solution (20 mL) and dried over Na_2_SO_4_. The solvent was removed under reduced pressure to give crude product. The crude residue was purified by column chromatography (DCM/EtOAc, 5:5) to provide **T16-peak2**. The product was dissolved in 5 mL of acetonitrile and lyophilized to give **T16-peak2** (0.0261 g, 78 %) as off -white solid. **^1^H NMR** (300 MHz, CDCl_3_) δ 8.36 (s, 1H), 8.25 (bd, *J* = 167.1 Hz, 1 H), 7.76 – 7.67 (m, 2H), 7.19 – 7.12 (m, 1H), 6.83 (dd, *J* = 7.8 Hz, 1H), 4. 20 - 4.10 (m, 4H), 3.80-3.70 (m, 1H), 3.3 – 3.14 (m, 1H), 2.90 – 2.68 (m, 5H), 2.26 (bd, *J* = 128.7 Hz, 3 H); LCMS rt 5.51 min, [M+H]^+^ 440.2 expected, 439.9 observed.

## Supplementary Note 1: Mechanisms for greater b-ion production

Although increased intensity of a precursor can lead to improved scoring by virtue of better MS/MS fragment sensitivity, PSMtag boosts peptide scores even in the case of decreased precursor intensity as seen in Fig. 1f. We suggested enhanced b-ion generation accounts for this increase, as shown in Fig. 2e. This observation has been noted for mTRAQ-labeled peptides and attributed to stabilization of b-ions via increased gas-phase basicity of mTRAQ relative to the unlabeled N-terminus^20^. This justification is supported by the mobile proton model (MPM) that describes the relationship of a proton’s mobility with the proton affinity of peptide functional groups^66,67^. Interestingly, any one site of protonation on PSMtag should not substantially increase the proton affinity of the N-terminus, yet we observe a similar stabilization. It is possible however, that the increased number of similarly basic sites on the N-terminus (from one label-free to three with PSMtag) provide a greater likelihood of detecting a b-ion due to the increased probability of retaining a charge. When C-terminal lysine residues are labeled, the gas-phase basicity of the C-terminus is lowered, allowing for greater proton mobility in accordance with the MPM. This model also offers an explanation for why the optimized collision energy for labeled peptides is lower than label-free overall.

## Supplementary Note 2: Limitations and opportunities in exploring chemical properties correlated with positive tag performance

Ordered by their correlation to Score, clusters in Fig. 1 and Extended Data Fig. 1 emerge that may be interpreted. For example, NumHAcceptors, NumAmide, Charge, and Mass are all negatively correlated with Score. Larger molecules in this screen contain more amide groups, which are hydrogen acceptors, which allow more Charge to be held during the electrospray ionization under acidic conditions used in this study. We cannot determine if any individually relate negatively to Score given their physical interconnectedness in this screen. Relatedly, a hypothesis requiring further investigation is that higher mass candidates may result in the collision energy, which is calculated by the MS to be proportional to mass, applied to the model peptides to result in over-fragmentation of the peptide backbone and thus poorer Score.

A select set of differential chemical descriptors related to whether or not the tag-candidates was observed at all are shown in Supplementary Fig. 5. Additional hypotheses related to synthesizability biases of different tag candidates may be generated from this comparison, but a downside to combinatorial synthesis in this context is the inability to quantify the yield of any one product. We cannot decouple synthesis yield from features like ionization propensity or adsorptive losses. This limits the ability of using data metrics like total or apex intensity of compound-peptide precursors for screening the mass tag candidates. Additionally, impact on compound ionization is only one beneficial metric. While increased MS1 peak intensities are desirable for sensitivity, MS2 fragmentation is crucial for identification. There can occasionally be a tradeoff between ionizability and fragmentability as described by the mobile proton model.

## Supplementary Note 3: Characterizing how amine-functionalized paramagnetic beads can remove excess NHS-ester tag

Following a ladder of ratios tested as shown in Supplementary Fig. 9a, unreacted PSMtag was removed below our limit of detection with a 20:1 mass ratio as described in the methods. Some peptide loss occurs with the addition of amine-bead-based quenching, 44% on average (Supplementary Fig. 9b). We expect the losses are driven by nonspecific interactions with the introduction of additional surface area of beads. We mitigate specific interactions of peptides with the amine-functionalized surfaces by addition of TEAB, similar to elution requirements for HILIC-based approaches. Additional loss may be caused by incomplete removal of sample from the reaction vessel as several *µ*L are left behind to avoid transferring the beads with the cleaned samples. As we tested with small sample amounts and volumes, the lost volume can total a relatively large fraction of starting sample amount. To mitigate this, one can incorporate a two-step elution. Further optimization is ongoing.

Initial evaluations were performed for all samples separately prior to combining for 5- or 9-plex injections. We recently tested the impact of a combined quench approach in which all samples are added to a single vessel for quenching and elution. Combined quenching mitigates the variability in sample loss across channels, improving reproducibility and limiting batch effects. This is demonstrated in Supplementary Fig. 9c comparing the relative recovery of peptides from samples quenched separately, recovered, and then combined (”separate”) to samples that were quenched in the same vessel with a proportional amount of the amine-functionalized beads and extracted together (”pooled”). A risk of a combined quench is the potential that excess tag from any channel may continue to label an under-labeled sample before the excess is fully quenched by the amine beads. An under-labeled sample is never desirable, however, a single, under-labeled sample may be omitted in a large scale study. Many under-labeled samples would indicate a systematic issue in sample preparation that would necessitate repeating an experiment. Thus, the primary concern for the pooled quench approach is if cross-channel labeling would skew protein quantitation for biological interpretation. To mimic such a scenario, a mixed proteome experiment with human- and yeast-derived peptides was carried out similar to our quantitative benchmark described above. Two samples, A and B, were labeled with PSMtag and a third unlabeled sample comprised of excess yeast was added to the vessel with amine-beads used for the combined quench, as depicted in Supplementary Fig. 9d. If substantial labeling of sample C yeast peptides competes with the amine-functionalized surface of the beads during the quenching time, we may expect deviation of the fold-change calculated from samples A and B. We stressed the system by adding greater excess of the unlabeled yeast in sample C relative to samples A and B, a more challenging case that may be likely from true samples. As Supplementary Fig. 9e demonstrates, our quantitative accuracy and precision is maintained for both the control and test injections, suggesting cross-channel labeling is minimal and has no significant effect on quantitative performance of PSMtag.

## Notes

https://www.parallelsq.org/psmtag

https://github.com/ParallelSquared/tag

### Competing Interest Statement

H.S., M.Y., S.S., D.B.S., M.A., W.V., A.Y., M.A.L., B.P. and N.S. are listed as inventors on a patent application for the tags described in this paper. R.T.K. have a financial interest in MicrOmics Technologies, whose LC workflow was used for the 1130 SPD demonstration. PTI has a collaborative research agreement with Thermo Fisher Scientific, the manufacturer of mass spectrometry instrumentation used in this research.

### Summary of Updates

Added ORCID to some authors. Added ORCID to some authors.

